# An evolutionary quantitative genetics model for phenotypic (co)variance under limited dispersal, with an application to socially synergistic traits

**DOI:** 10.1101/393538

**Authors:** Charles Mullon, Laurent Lehmann

## Abstract

Darwinian evolution consists of the gradual transformation of heritable traits due to natural selection and the input of random variation by mutation. Here, we use a quantitative genetics approach to investigate the coevolution of multiple quantitative traits under selection, mutation, and limited dispersal. We track the dynamics of trait means and variance-covariances between traits that experience frequency-dependent selection. Assuming a multivariate-normal trait distribution, we recover classical dynamics of quantitative genetics, as well as stability and evolutionary branching conditions of invasion analyses, except that due to limited dispersal, selection depends on indirect fitness effects and relatedness. In particular, correlational selection that associates different traits *within*-individuals depends on the fitness effects of such associations *between*-individuals. We find that these kin selection effects can be as relevant as pleiotropy for the evolution of correlation between traits. We illustrate this with an example of the coevolution of two social traits whose association within-individuals is costly but synergistically beneficial between-individuals. As dispersal becomes limited and relatedness increases, associations between-traits between-individuals become increasingly targeted by correlational selection. Consequently, the trait distribution goes from being bimodal with a negative correlation under panmixia to unimodal with a positive correlation under limited dispersal.

## 1 Introduction

Understanding how heritable quantitative traits are molded by natural selection and mutation has been a long-standing goal of evolutionary biology. This research endeavour has led to an abundant theoretical literature that seeks to understand the roles of ecology and genetics in the gradual transformation of quantitative phenotypes. Notwithstanding this abundance, models of gradual evolution usually follow one of two approaches, depending on whether the focus is put on ecological or genetic processes.

One approach consists in investigating the invasion success of a rare phenotypic mutant (i.e., an evolutionary invasion analysis, e.g. Michod, 1979, Eshel and Feldman, 1984, Parker and Smith, 1990, Eshel et al., 1997; also referred to as “Adaptive Dynamics”, e.g., Dercole and Rinaldi, 2008, for a textbook treatment) and places emphasis on ecology (or on how organisms interact with one another via effects on resources and the environment). In most practical applications, this emphasis comes at the expense of genetics realism. In particular, trait dynamics inferred from invasion analyses most often assume that mutations have weak quantitative effects and are so rare (relative to the strength of selection) that at most two alleles can segregate in the population. In this case, a sensitivity analysis of the invasion fitness of a rare mutant in a resident monomorphic population that is at its ecological equilibrium (e.g., Michod, 1979, Eshel and Motro, 1981, Eshel and Feldman, 1984, Taylor, 1989, Parker and Smith, 1990, Charlesworth, 1994) can be used to understand gradual trait evolution and the ecological transformations due to this evolution (Metz et al., 1996, Geritz et al., 1998, Rousset, 2004, Dercole and Rinaldi, 2008, Metz, 2011). Evolutionary invasion analysis is therefore particularly well-suited to investigate the evolution of traits under ecological feedbacks and the frequency-dependent selection that emerges due to such feedbacks (e.g., Kisdi and Geritz, 2009, Lion, 2018, and references therein). This approach has revealed that in the presence of trade-offs, gradual evolution under ecological feedbacks often leads to the emergence of polymorphism. Here, the population evolves under directional selection to express a trait value such that any rare mutant has an advantage over the common resident (Eshel and Motro, 1981, Eshel, 1983, Taylor, 1989, Christiansen, 1991, Abrams et al., 1993b). As a result, the population subsequently splits into two lineages of distinct phenotypes, or morphs, in a process referred to as evolutionary branching (Geritz et al., 1998; see Rueffler et al., 2006, Kisdi and Geritz, 2009, for reviews).

By contrast to invasion analysis, evolutionary quantitative genetics models of gradual evolution tend to be more preoccupied with the genetic basis of traits (Roff, 1997, Lynch and Walsh, 1998). Importantly, quantitative genetics models envisage that substantial heritable phenotypic variation segregates in the population. The continuum-of-alleles model, in particular, posits that quantitative traits are determined by a continuum of possible alleles produced by mutation (e.g., Kimura, 1965b, Latter, 1970, Fleming, 1979, Bürger, 1986). A quantitative genetics approach aims to investigate the roles of selection and mutation in the gradual evolution of a phenotypic distribution of arbitrary complexity. Due to the complication of dealing with multiple phenotypic variants, however, analytical explorations of quantitative genetics models usually come at the expense of generality. Notably, the vast majority of quantitative genetics models of traits under frequency-dependent selection, which either is implicit or emerges from ecological interactions, focuses on the evolution of mean phenotypic values in the population, assuming that heritable phenotypic variation is constant (i.e., additive genetic variances and covariances are fixed, e.g., Lande, 1976, 1981, Iwasa et al., 1991, Gomulkiewicz and Kirkpatrick, 1992, Abrams et al., 1993a, Iwasa and Pomiankowski, 1995, Day and Taylor, 1996, Tazzyman and Iwasa, 2009, Nuismer et al., 2010, Connallon, 2015).

But phenotypic variance should be especially sensitive to frequency-dependent selection. This is because such selection either favors or disfavors rare variants that differ from the most common, and thus either increases or decreases trait variance (Slatkin, 1980, Taper and Chase, 1985, Taylor and Day, 1997, Day and Proulx, 2004, Sasaki and Dieckmann, 2011, Wakano and Iwasa, 2013, Wakano and Lehmann, 2014, Débarre et al., 2014, Débarre and Otto, 2016). In fact, recent quantitative genetics models investigating populations of individuals experiencing frequency-dependent interactions have revealed links between the dynamics of phenotypic variance and evolutionary branching (Sasaki and Dieckmann, 2011, Wakano and Iwasa, 2013, Wakano and Lehmann, 2014, Débarre et al., 2014, Débarre and Otto, 2016), thereby extending the links between the dynamics of the phenotypic mean in quantitative genetics models and directional selection in invasion analysis models (Charlesworth, 1990, Iwasa et al., 1991, Taper and Case, 1992, Abrams et al., 1993a; for reviews: Abrams, 2001, Lion, 2018). Specifically, evolutionary branching occurs in a quantitative genetics model when the phenotypic variance is predicted to grow without bound while the phenotypic mean is held constant, under the assumption that the phenotypic distribution is normal (this assumption allows to only have to consider the dynamics of the mean and variance of the phenotypic distribution, Wakano and Iwasa, 2013, Wakano and Lehmann, 2014, Débarre et al., 2014, Débarre and Otto, 2016). As evolutionary branching occurs, the variance may in fact converge to a bounded value (see Fig. 2e-f of Débarre and Otto, 2016), but these dynamics cannot be captured by models that assume that the phenotypic distribution is normal and thus unimodal (instead of a bi- or multi-modal distribution; see Sasaki and Dieckmann, 2011 and Appendix D of Débarre and Otto, 2016 for a relaxation of the unimodal assumption). In spite of this limitation, quantitative genetics approaches have been useful to investigate relevant factors for frequency-dependent selection and evolutionary branching, such as genetic drift (with fixed, Wakano and Iwasa, 2013, or fluctuating, Débarre and Otto, 2016, population size) or the interaction between multiple traits (Débarre et al., 2014).

One factor that is particularly relevant for frequency-dependent interactions is limited dispersal. This is because limited dispersal creates genetic structure, whereby individuals that interact and compete with one another are more likely to share identical alleles at loci determining social or competitive traits than individuals randomly sampled from the population, resulting in kin selection on traits (Hamilton, 1964, Michod, 1982, Frank, 1998, Rousset, 2004). Using an invasion analysis, a number of models have investigated the conditions that lead to disruptive selection (usually followed by evolutionary branching) due to frequency-dependent interactions among individuals under limited dispersal (Day, 2001, Ajar, 2003, Rousset, 2004, Mullon et al., 2016, Parvinen et al., 2018; see also Svardal et al., 2015 for evolutionary branching due to spatial and temporal heterogeneities in selection but without kin selection). Using a quantitative genetics approach, Wakano and Lehmann (2014) (WL14 hereafter) found branching conditions equivalent to those obtained from invasion analysis by studying the dynamics of the variance of a trait under limited dispersal. The analysis of frequency-dependent and disruptive selection under limited dispersal has helped reveal further connections between invasion analysis and fundamental branches of evolutionary theory. In particular, Ajar (2003), WL14 and Mullon et al. (2016) expressed disruptive selection coefficients in terms of relatedness coefficients, which are quantities central to population genetics, kin selection and social evolution theory (i.e., the evolution of traits that influence the fitness of their actor and recipient, such as helping or harming, e.g., Hamilton, 1964, Frank, 1998, Rousset, 2004, Wenseleers, 2010; see also Kisdi, 2016 for a kin selection perspective on evolutionary branching of dispersal).

In this paper, we incorporate two additional factors that have previously been omitted in the gradual evolution of quantitative traits when selection is frequency-dependent and dispersal is limited. First, we consider the joint evolution of multiple traits (whereas WL14 focuses on a single trait). This enables us to investigate how phenotypic covariances among traits within individuals are molded by frequency-dependent selection and pleiotropic mutations (i.e., when traits share a common genetic basis so that mutations have correlated effects across traits). Second, we model the coupled dynamics of the phenotypic means and (co)variances (whereas WL14 looks at the dynamics of the variance only once selection on means is negligible). This allows for a more complete picture of the dynamics of the phenotypic distribution. By expressing these dynamics in terms of relatedness coefficients, we further connect kin selection theory with the evolutionary quantitative genetics of multiple traits (Lande, 1979, Lande and Arnold, 1983, Phillips and Arnold, 1989, Brodie et al., 1995; in particular with the evolution of the **G**-matrix of additive genetic variance-covariance, Steppan et al., 2002, Arnold et al., 2008)

The rest of this paper is organized as follows. We describe the life-cycle and population structure under consideration in section 2. Our first result, presented in section 3.1, is an equation for the one-generational change of a multi-variate phenotypic distribution under limited dispersal, mutation, and selection. Next, in section 3.2, we present a closed dynamical system for the mean vector and variance-covariance matrix of the phenotypic distribution, under the assumption that the distribution in the whole population is normal. Further, we express this dynamical system in terms of effects on individual fitness and relatedness in section 3.3, and highlight some equilibrium properties of our dynamical system in section 3.4. In section 3.5, we apply our framework to study the coevolution of two traits that have socially synergistic effects between individuals. Finally, we discuss the implications of our results for understanding patterns of intra-specific variation, with special reference to social and competitive traits.

## 2 Model

We consider a population of haploid individuals, divided among an infinite number of groups, each of fixed size *N* (the total population size is therefore constant). Each individual bears a multidimensional phenotype that consists of *n* genetically determined quantitative traits. The discrete-time life cycle of this population is as follows. (1) Groups may go extinct (in which case all *N* adult individuals in a group die before reproduction) and do so independently of one another. (2) Adults reproduce clonally (producing offspring in sufficient number for the size of each group in the population to be *N* by the end of the life cycle) then either survive or die, which frees up breeding spots. (3) The phenotype of each individual independently mutates with probability *v*, causing a random quantitative deviation in trait values. (4) Each offspring either remains in its natal group, or disperses to another randomly chosen group (i.e., we consider the island model of dispersal, Wright, 1931, Rousset, 2004). (5) Offspring compete locally in each group to fill open breeding spots, if any.

This life-cycle allows for one, several, or all adults to die per life-cycle iteration (including through whole group extinction before reproduction). Generations can thus overlap but the expression of traits is assumed to be independent of age (e.g., the fertility or mortality of an individual is independent from its age and that of any other individual it interacts with). Dispersal can occur before or after density-dependent competition (as long as the number of adults in each groups remains constant), and in groups, so that more than one offspring from the same natal patch can establish in a non-natal patch. This life cycle is equivalent to that considered in Mullon et al. 2016, except that here, we allow for the constant input of mutations in the population (step 3 of the life cycle).

## 3 Results

### 3.1 Dynamics of the phenotypic distribution

In order to track phenotypic evolution in the population, we denote by *p*_*t*_(**z**) the phenotypic density distribution in the population at a demographic time point *t*, where **z** = (*z*_1_, *z*_2_, …, *z*_*n*_) ∈ ℝ^*n*^ is a vector collecting the variable *z*_*a*_ for each trait *a* 1, …, *n*. To capture group structure, we introduce the density distribution *ϕ*_*t*_ of phenotypic groups states in the population at time *t* (i.e., *ϕ*_*t*_ (**z**_1_, **z**_2_, …, **z**_*N*_) is the probability density distribution of groups in which individuals arbitrarily labelled 1 to *N* have phenotypes **z**_1_, **z**_2_, …, **z**_*N*_, respectively, see Appendix eq. A-3 for more details).

In Appendix A, we show that the recurrence equation for the phenotypic distribution in the population from demographic time step *t* to *t* +1 (one iteration of the life cycle) can be expressed as

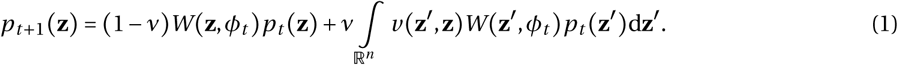

The first summand represents changes in the distribution due to reproduction and survival of individuals that have not mutated (with probability 1− *ν*), and the second summand, changes due to those that have (with probability *v*; and where *v* (**z**′, **z**) denotes the probability density function for the event that an individual mutates from **z**′ to **z** given that a mutation has occurred). The quantity *W*(**z**, *ϕ*_*t*_) in eq. (1) is a measure of fitness of phenotype **z**, which depends on the way phenotypes are distributed across groups (i.e., on *ϕ*_*t*_). When *W*(**z**, *ϕ*_*t*_)> 1, the frequency of **z** in the population increases due to selection, and conversely decreases when *W*(**z**, *ϕ*_*t*_) < 1.

To gain insights into the fitness measure *W*(**z**, *ϕ*_*t*_), note that recurrence eq. (1) has the same form as the classical recurrence of the phenotypic distribution in well-mixed populations under the continuum-of-alleles model (e.g., Kimura, 1965b, eqs. 1-2; Fleming, 1979, eq. 2.4; Bürger, 1986, eq. 1; Taylor and Day, 1997, eq. 1; Champagnat et al., 2006, eq. 4.1). In a well-mixed population of constant size, the fitness of phenotype **z** is equal to the *individual* fitness of a focal individual with phenotype **z**; namely, its expected number of successful offspring produced over one iteration of the life-cycle (including self through survival). Because individuals interact at random in a well-mixed population, such individual fitness function only depends on the population wide phenotypic distribution, *p*_*t*_(**z**) (i.e., the phenotype of any group neighbor to a focal individual, captured by *ϕ*_*t*_, is in fact independently and identically distributed according to *p*_*t*_(**z**)). The fitness of an individual with phenotype **z** in a population with trait distribution *p*_*t*_(**z**) can thus be written as *w*(**z**, *p*_*t*_(**z**)), and *W*(**z**, *ϕ*_*t*_)= *w* (**z**, *p*_*t*_ **(z)**) in eq. (1) (to distinguish between fitness at the phenotype and individual level, we generically denote the former by an upper case *W* and the latter by a lower case *w)*.

Defining individual fitness in terms of expected number of successful offspring is standard in social evolution theory (e.g., Hamilton, 1964, Rousset, 2004), and takes its roots in population dynamics: when *w*(**z**, *p*_*t*_(**z)**) > 1, the number of individuals with phenotype **z** increases and conversely decreases when *w* (**z**, *p*_*t*_ (**z**)) < 1 (e.g., eq. 2.2 of Nagylaki 1992). As such, it is sometimes referred to as “absolute” fitness. Many quantitative genetics models, by contrast, employ the notion of “relative” fitness to track changes in phenotypic frequencies. This can stem from two non-mutually exclusive modelling choices: (1) one in fact considers the effect of the phenotype on a vital rate, *f*(**z**, *p*_*t*_ (**z)**) (such as fecundity or offspring survival), that influences the number of offspring that enter competition before regulation, which requires normalisation by mean vital rate, *W* (**z**, *ϕ*_*t*_)*=w*(**z**, *p*_*t*_(**z))=***f*(**z**, *p*_*t*_(**z))/**[∫ *f*(**z**, *p*_*t*_ (**z)**) *p*_*t*_ (**z**)d**z]**; (2) the population size fluctuates, in which case it is necessary to normalise by mean fitness, *W*(**z**, *ϕ*_*t*_) = *w*(**z**, *p*_*t*_(**z))/**[∫ *w* (**z**, *p*_*t*_(**z)**) *p*_*t*_(**z**)d**z**]. In our model, because group size and therefore population size is constant, *W*(**z**, *ϕ*_*t*_) in eq. (1) can be viewed as an absolute measure of fitness.

In contrast to a well-mixed population, the fitness of an individual *w* in a dispersal-limited group-structured population depends on the way phenotypes are distributed across groups (so on *ϕ*_*t*_), and specifically on the collection of phenotypes carried by the individuals that belong to its own group. The fitness of an individual with phenotype **z** in a population with group distribution given by *ϕ*_*t*_ can thus be written as *w*_*µ*_(**z**, *ϕ*_*t*_), where *µ* is the collection of phenotypes carried by individuals in the focal group (formally, *µ* is a counting measure in our analysis – see Appendix A.1.2 – but for the purpose of the main text, it can simply be thought of as the phenotypic composition of the focal group). In terms of this individual fitness function, we find that the fitness at the level of the phenotype that is relevant for phenotypic dynamics, *W* (**z**, *ϕ*_*t*_) in eq. (1), is

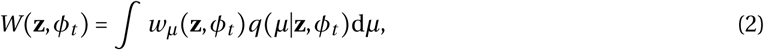

where the integral runs through every possible group state, *µ*, and *q(µ*|**z**, *ϕ*_*t*_) is the probability density function for the event that an individual randomly picked from the collection of all carriers of the **z** phenotype in the population at time *t* resides in a group in state *µ* (see eq. A-17 in Appendix A for derivation). According to eq. (2), *W*(**z**, *ϕ*_*t*_) is the average expected number of successful offspring of an individual with phenotype **z**, where the average is taken over all group states *µ* in which an individual with phenotype **z** can reside at time *t*.

An alternative interpretation for *W* (**z**, *ϕ*_*t*_) can be reached by noting that because there is an infinite number of possible alleles, all individuals with the same phenotype **z** belong to the same genetic lineage (as the same allele cannot appear twice via mutation). The function *q*(*µ*|**z**, *ϕ*_*t*_) in eq. (2) then corresponds to the probability that an individual sampled from this lineage at time *t* resides in a group in state *µ*. As such, *W* (**z**, *ϕ*_*t*_) can be interpreted as the average direct fitness of an individual randomly sampled from the lineage of individuals carrying phenotype **z** at time *t*. If on average individuals from the **z**-lineage produce more than one successful offspring at time *t*, this lineage will be larger at time *t+*1 and in a population of constant size, the frequency of individuals with phenotype **z** will increase. The fitness measure *W* (**z**, *ϕ*_*t*_) can thus be seen as the multiallelic version of the concept of a mutant’s *lineage fitness* used previously in invasion analyses (which turns out to be equal to the mutant’s growth rate when the mutant is rare in an otherwise monomorphic population, Mullon et al., 2016, Lehmann et al., 2016; see also Wild, 2011 for similar branching processes approach to social evolution in group-structured populations). We will therefore refer to *W*(**z**, *ϕ*_*t*_) as the lineage fitness (or average direct fitness) of phenotype **z**, keeping in mind that unlike in invasion analyses, *W*(**z**, *ϕ*_*t*_) here applies for any frequency of **z** (rare or common) and for any population composition (monomorphic or polymorphic).

### 3.2 Tracking the dynamics of the phenotypic distribution

The dynamical equation for the phenotypic distribution eq. (1) has no straightforward solution, even when the population is well-mixed (Kimura, 1965b, Fleming, 1979, Lande, 1979, Bürger, 1986). Under limited dispersal, this problem is further complicated by the necessity of simultaneously tracking the dynamics of group composition *ϕ*_*t*_. To proceed further in our analysis and track the dynamics of the phenotypic distribution, we therefore make additional assumptions.

#### 3.2.1 Weak selection, weak mutation, normal closure and quasi-equilibrium of local genetic associations

We first assume that selection is weak, in the sense that the phenotypic variance in the population is small (allowing for second-order approximation of lineage fitness, see Appendix B.1.1 for details, and Iwasa et al., 1991 for a similar approach for quantitative genetics of traits under frequency-dependent selection in well-mixed populations). This enables us to express lineage fitness in terms of time-dependent local genetic associations among individuals of the same group (i.e., relatedness coefficients), w hich c apture relevant m oments of the distribution of group composition *ϕ*_*t*_, and therefore avoids us having to keep track of the full distribution (see eq. B-1-B-4). Next, we assume that mutations are rare, so that we can ignore the joint effects of selection and mutation on the phenotypic distribution over one time period (see Appendix B.1.2).

Following previous authors (e.g., see Taylor and Day, 1997, Wakano and Iwasa, 2013, Débarre et al., 2014, Wakano and Lehmann, 2014, Débarre and Otto, 2016, for social traits), we further assume that the processes of selection and mutation are such that *p*_*t*_ (**z**) is approximately multivariate normal (allowing for moment closure, see Appendix B.2.1). The assumption of normality is a strong one but it is noteworthy that it does not require that the realized distribution of phenotypes within a focal group at any given demographic time period is normal. In addition, the assumption of normality has been shown to give accurate predictions for the change of mean and variance, which is our main goal, even when selection generates significant deviations from normality (in well-mixed populations, Turelli and Barton, 1994). Under the assumption of normality, the distribution *p*_*t*_ (**z**) is characterised by its mean vector 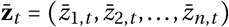, whose *a*-entry is the average value of trait *a* in the population at time period 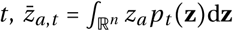; and its variance-covariance matrix ***G***_*t*_ whose (*a, b*)-entry is the (co)variance among traits *a* and *b* in the population at time period *t*, 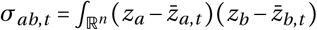*p*_*t*_(**z**)d**z**. The dynamics of *p*_*t*_(**z**) can therefore be tracked through the dynamics of its mean vector 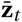 and variance-covariance matrix ***G***_*t*_.

But due to limited dispersal, the dynamics of 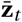 and ***G***_*t*_ still depend on time-dependent local genetic associations among individuals of the same group. To close evolutionary dynamics on 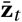 and ***G***_*t*_ and avoid tracking the dynamics of these genetic associations, we assume that selection is weak relative to dispersal so that genetic associations reach their steady state before significant changes have occurred in the phenotypic distribution, *p*_*t*_(**z**) (see section B.2.2 for details). This “quasi-equilibrium” assumption, which is frequently used in population genetic theory (e.g., Kimura, 1965a, Nagylaki, 1993, Kirkpatrick et al., 2002, Roze and Rousset, 2005, 2008), finally allows us to characterize the dynamics of *p*_*t*_ (**z**) entirely by the coupled dynamics of its mean vector 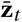 and variance-covariance matrix ***G***_*t*_.

#### 3.2.2 Dynamics of phenotypic mean vector and variance-covariance matrix

Under the above assumptions, we show in Appendix B that the coupled changes of the mean trait vector and variance-covariance matrix over one demographic time period are respectively given by

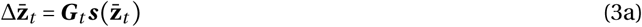

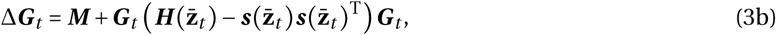

where 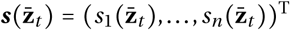 (.^T^ denotes the transpose of a vector or matrix) is a *n*×1 is vector of directional selection coefficients (or selection gradients), i.e., 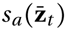 is the first-order, marginal, effect of an (infinitesimal) change in trait *a* away from the population mean 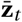 on lineage fitness 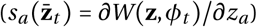. The *n* × *n* matrix ***M*** collects the effects of mutation; its (*a, b*)-entry,

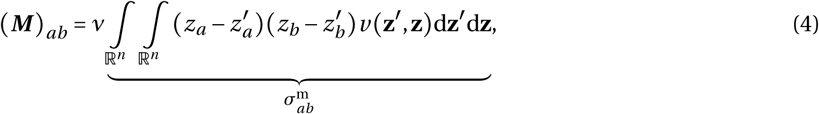

is the product of the mutation probability, *v*, with the (co)variance, 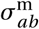, in mutational effects on traits *a* and *b* conditional on the appearance of a mutation (which captures the pleiotropic effects of mutations on *a* and *b*: when 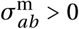, mutations tend to change *a* and *b* in a similar way; and when 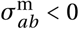 in opposite ways).

The *n* × *n* Hessian matrix 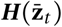 collects the second-order effects of traits on lineage fitness; its (*a, b)*-entry 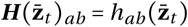 is the marginal effect of joint changes in traits *a* and *b* away from the population mean 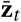 on lineage fitness 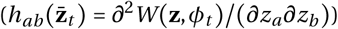. Finally, the notation 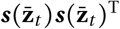 denotes the outer product between two column vectors, so that 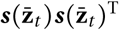 is *n* × *n* matrix with (*a, b*)-entry 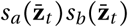.

#### 3.2.3 Directional, disruptive, and correlational selection coefficients

Dynamical equations (3) have the same form as in well-mixed populations (e.g., eqs. 1-2 of Phillips and Arnold, 1989, see also eq. 7 of Lande, 1979 and eqs. 6 and 15 of Lande and Arnold, 1983). In such models, the effects of selection depend on the marginal effects of traits on *individual* rather than *lineage* fitness. Nevertheless, the parallels between eq. (3) and previous works allow us to use the same vocabulary and interpretations on the evolution of phenotypic means and (co)variances (Brodie et al., 1995). First, the evolution of the mean of each trait (eq. 3a) depends on the vector of directional selection (or the selection gradient), 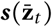, which points in the direction favored by selection in multivariate phenotypic space (Lande, 1979). The effect of directional selection on the mean of each trait, however, is constrained by the genetic variation available and these constraints are captured by ***G***_*t*_ in eq. (3a) (Lande, 1979).

Second, the evolution of the variance-covariance matrix ***G***_*t*_ (eq. 3b) depends on the effects of mutations (***M)***, of directional selection 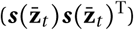, and of quadratic selection given by the matrix 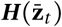 (Lande, 1979, Lande and Arnold, 1983, Phillips and Arnold, 1989). This matrix 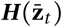 captures two relevant features of selection. First, the sign of its diagonal entry (*a, a)* indicates whether selection favors a decrease (when 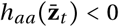) or an increase (when 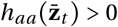) in the variance of trait *a* when this trait evolves in isolation of other traits (Phillips and Arnold, 1989), hence 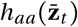 is referred to as the coefficient of disruptive selection on trait *a*. Second, the off-diagonal entry (*a, b)* tells us whether selection favors a positive (when 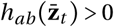) or negative (when 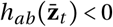) covariance or correlation among traits *a* and *b*. The off-diagonal entry 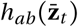 is therefore referred to as the coefficient of correlational selection among traits *a* and *b* (Lande and Arnold, 1983, Phillips and Arnold, 1989).

### 3.3 Selection in terms of individual fitness effects and relatedness coefficients

So far, the effects of limited dispersal on evolutionary dynamics (eqs. 1 and 3) have been hidden behind the notion of lineage fitness, *W*(**z**, *ϕ*_*t*_). To highlight more tangibly how selection depends on limited dispersal, we express the selection coefficients (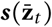 and 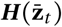) in terms of the effects of traits on individual fitness and relatedness. For this, let us first rewrite the individual fitness of a focal individual, that we label as individual “*i* “, as a function 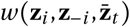 of three arguments: (1) the phenotype of the focal individual, **z_*i*_** = (*z*_*i*,1_, *z*_*i*,2_, …, *z*_*i,n*_); (2) the collection of phenotypes of its *N* − 1 neighbors **z**_− *i*_ = (**z**_1_, … **z**_*i*−1_, **z**_*i*+1_, …, **z**_*N*_) (where **z**_*j*_ = *z*_*j*,1_, *z*_*j*,2_, …, *z*_*j,n*_) is the phenotype of a neighbor indexed *j*); and (3) the average phenotype in the population 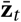 (see eq. 15 for an example of such a fitness function). This individual fitness function is equal to the fitness function *w*_*µ*_(**z**, *ϕ*_*t*_) that appears in eq. (2),

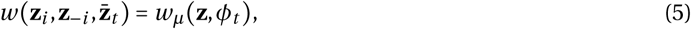

when focal phenotype is **z**_*i*_ = **z**, the state of the focal group is *µ* = {**z**_*i*_} ∪ **z**_−*i*_ = (**z**_1_, …, **z**_*N*_), and groups other than the focal one are considered to be monomorphic for the population average 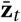 (i.e., we consider that all individuals in other groups express 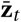 so that the distribution *ϕ*_*t*_ is delta peaked on 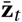; we can do this because the phenotypic distribution is assumed to be centered around 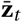 with small variance and individuals from different groups interact at random in the island model; see Iwasa et al., 1991 for a similar approach in panmictic populations).

We further introduce two neutral relatedness coefficients that will be relevant for selection: let 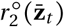 and 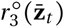 respectively be the probabilities that, in the absence of selection and when the population phenotypic average is 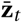, one and two neighbors of a focal individual carry a phenotype that is identical-by-descent to that of the focal (i.e., the set of individuals under consideration have a common ancestor). Alternatively, 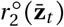 and 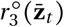 can be interpreted as the probabilities that, in the absence of selection and when the population phenotypic average is 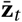, two and three individuals sampled in the same group carry identical-by-descent phenotypes. This interpretation is in line with the definition of relatedness in the infinite island model (see e.g., Rousset, 2004, Taylor et al., 2007, for further considerations on relatedness in the finite island model).

#### 3.3.1 Directional selection

We find that the selection gradient on a trait *a* can be expressed as

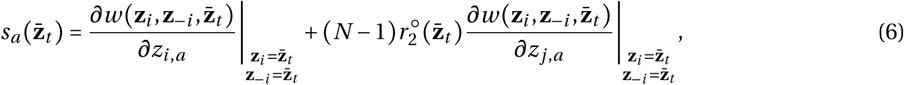

where 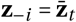 means that the derivative is evaluated when all neighbors express the mean phenotype 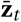 (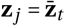 for all *j*≠*i*). The first derivative in eq. (6) captures the direct effect of trait *a*: the effect of a change in trait *a* in a focal individual on its own fitness. In a well-mixed population, this is all that matters for directional selection (i.e., 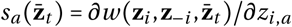 when the population size is constant, Phillips and Arnold, 1989^1^). The second derivative, which is weighted by pairwise relatedness 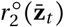, is the indirect effect of trait *a*: the effect focal fitness of a change in trait *a* in a neighbor of the focal (we arbitrarily chose this neighbor to be individual *j*≠*i)*. The selection gradient on trait *a*, eq. (6), is therefore the inclusive fitness effect of trait *a* (Hamilton, 1964, Rousset, 2004). Hence, in the absence of covariance among traits, the change in mean trait value is proportional to this trait’s inclusive fitness effect (substituting eq. 6 into 3a with the off-diagonal elements of **G**_*t*_ all zeros). This finding is in line with much previous modeling work on the quantitative genetics of spatially- or family-structured populations (for e.g., Cheverud, 1985, Queller, 1992a, b, Frank, 1998, McGlothlin et al., 2014, Wakano and Lehmann, 2014).

#### 3.3.2 Correlational and disruptive selection

We find that the correlational selection coefficient on two traits *a* and *b* (or the disruptive selection coefficient when *a* = *b*) can be expressed as the sum of two terms,

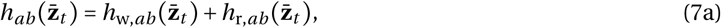

where the first term,

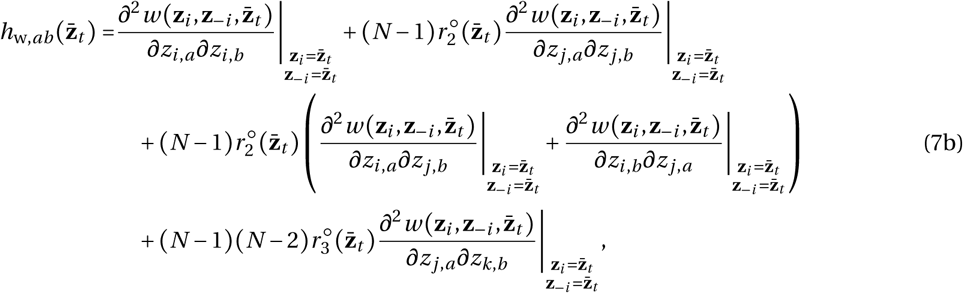

depends on the effects of joint changes in traits *a* and *b* within-(first line of eq. 7b) and between-individuals (second and third line of eq. 7b) on focal fitness. The first derivative on the first line of eq. (7b) is the effect of a joint change in traits *a* and *b* in a focal individual on its own fitness, which can be viewed as the *direct* synergistic effects of traits *a* and *b* (Figure 1a). In a well-mixed population, there are no other effects participating to correlational selection (i.e., 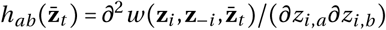, Phillips and Arnold, 1989).

**Figure 1:**
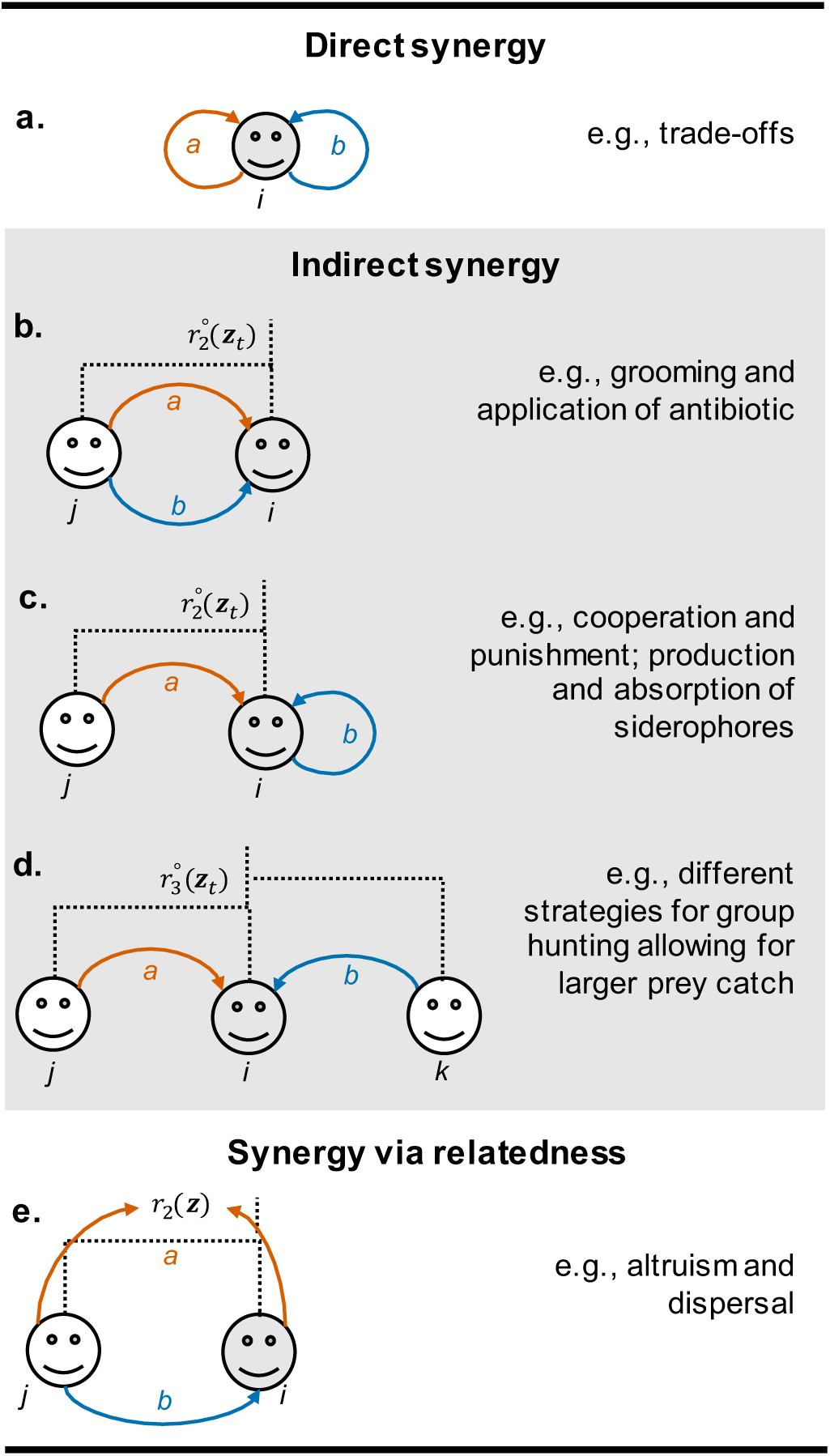
Within- and between-individual fitness effects relevant for correlational selection and examples of traits likely to be influenced by such effects. As revealed by eq. (7), there are five types of fitness effects due to perturbations in two traits *a* and *b* that are relevant for correlational selection when dispersal is limited: **a.** effect of a joint changes in *a* and *b* within the focal individual (first term of eq. 7b); **b.** effect of joint changes in *a* and *b* within neighbors of the focal (second term of eq. 7b, weighted by neutral pairwise relatedness, 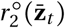); **c.** effect of joint changes in *a* and *b* between the focal (here *b*) and its neighbors (here, *a*; second line of eq. 7b, weighted by 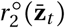); **d.** effect of joint changes in *a* and *b* between neighbors of the focal (third line of eq. 7b, weighted by neutral three-way relatedness, 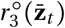); **e.** the effect of the indirect effect of one trait (here *b*) multiplied to the effect of the other (here *a*) on pairwise relatedness, which reflects the tendency of relatives to receive the effects of *b* (eq. 7c).

But when dispersal is limited (so that 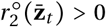 and 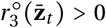, three *indirect* synergistic effects become relevant for correlational selection. These are the effect of a change in: (i) both traits in one neighbor of the focal (second derivative on the first line weighted by the neutral probability that the focal and this neighbor are identical-by-descent, 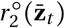, Figure 1b); (ii) in one trait in the focal and in the other in a neighbor (the two derivatives of the second line weighted by 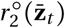, Figure 1c); and (iii) in one trait in a neighbor and in the other in another neighbor indexed as *k* (last derivative weighted by the neutral probability that the focal and these two neighbors are identical-by-descent, 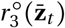, Figure 1d). Collectively, these terms capture the effects of non-random (due to limited dispersal) frequency-dependent interactions among individuals on correlational selection, revealing that under limited dispersal, selection favors the association of traits when these have positive effects between individuals.

The second term of eq. (7a), 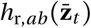, captures another type of synergistic effect relevant for correlational selection in group-structured populations. This term can be expressed as

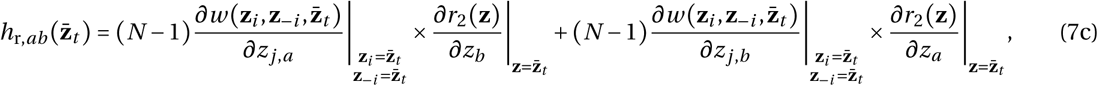

where 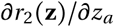 is the effect of trait *a* on the probability that a neighbor of a focal individual with phenotype **z** carries a phenotype that is identical-by-descent to that of the focal (and 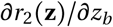 the effect of trait *b*). We refer to this as the effect of traits on relatedness. So eq. (7c) reveals that correlational selection depends on the product between the indirect effect of one trait (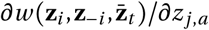 and 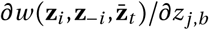), with the effect of the other trait on relatedness. Such synergy via relatedness (Figure 1e) reflects that in group structured populations, selection will favor an association among two traits when such an association results in indirect fitness benefits (e.g., trait *a* is cooperative, 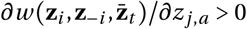) being preferentially directed towards relatives (e.g., trait *b* is the tendency to stay in natal group, 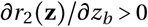.

Group-structure and limited dispersal may thus lead to significant changes to the way selection molds phenotypic correlations, especially when traits have synergistic effects that are either indirect (Figure 1b-d) or via relatedness (Figure 1e). This will be illustrated later when we study the coevolution of two social traits in section 3.5. Before doing so, let us remark that when a single traits evolves (*n* = 1) and the selection gradient on this trait is zero 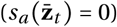, the change in phenotypic variance that we obtain (eq. 7 substituted into eq. 3b) reduces to previously derived expressions from quantitative genetics in the island model (eqs. 26 and 31 of Wakano and Lehmann, 2014). Further, eqs. (6)-(7) are consistent with evolutionary invasion analyses, i.e., with the first- and second-order effects of selection on the growth rate (or invasion fitness) of a rare mutant that arises in a monomorphic group-structured population and that differs from the resident in a single (eqs. 8 & 9 of Ajar, 2003) or multiple (eqs. 12 & 13 of Mullon et al., 2016) traits. We discuss further the correspondence between quantitative genetics, invasion analyses, and adaptive dynamics models in the next section, in which we study the equilibrium properties of the phenotypic distribution.

### 3.4 Equilibrium properties of the phenotypic distribution

Eq. (3) with eqs. (6)-(7) is a closed dynamical system that allows to track the evolution of the mean trait value and of the (co)variance between traits. In this section, we first investigate key features of the equilibrium of these phenotypic dynamics, and then discuss their connections with notions of evolutionary stability that come from invasion analyses and adaptive dynamics.

#### 3.4.1 Equilibrium mean trait values

We denote the mean trait vector and variance-covariance matrix of the equilibrium phenotypic distribution by 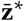 and ***G*\***, respectively. Such equilibrium simultaneously satisfies 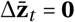 and Δ***G***_*t*_ = **0** (where **0** is used to denote a *n* vector and *n* × *n* matrix whose entries are all zero, respectively). Rather than solving both systems of equations simultaneously, we can use the fact that in eq. (3a), the matrix ***G*** is a positive-definite matrix with real-entries (since it is a variance-covariance matrix). From standard linear algebra theory (Hines, 1980, Leimar, 2005, 2009), it then follows that the equilibrium for the phenotypic means must satisfy

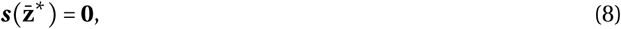

i.e., all selection gradients (eq. 6) vanish at 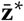, independently of the ***G*** matrix. An alternative argument to ignore the ***G*** matrix when determining the equilibrium trait vector **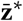** can be made from our assumption that (co)variances are small (weak selection). As a consequence, the dynamics of the ***G*** matrix are slower than those of the mean vector 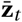 (see eq. B-20 in Appendix B.2). Trait means should therefore reach their equilibrium before the variance-covariance ***G*** matrix stabilizes.

We can further ask whether a population with a mean vector that is close to an equilibrium 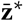 will eventually converge to it as a result of selection and mutation. From the fact that ***G*** is positive-definite, it can be shown (see Leimar, 2009, for e.g.) that a necessary condition for a population to converge to 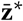 for all possible ***G*** matrices is that the Jacobian matrix 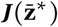 of the selection gradients with (*a, b*) entry

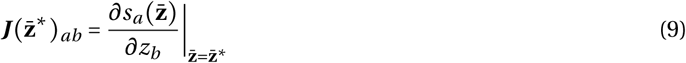

is negative-definite at 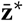, which means that the symmetric real part of 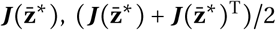 has only negative eigenvalues. This type of equilibrium is referred to as (strongly) convergence stable (Leimar, 2005, 2009).

#### 3.4.2 Equilibrium variance-covariance matrix

The dynamics of the variance-covariance matrix can then be studied at a convergence stable equilibrium 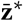 for mean trait values (eq. 8). In this case, the equilibrium ***G*** * for the variance-covariance matrix solves

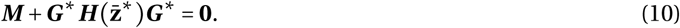

Eq. (10) has an admissible solution (i.e., such that ***G*\*** is positive-definite) if, and only if, the Hessian matrix, 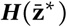, is negative-definite (Bhatia, 2015). This corresponds to the case under which selection is stabilizing at 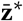. In fact, if 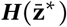 is negative-definite, then the population will remain unimodally distributed around the mean vector 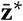 and exhibit a variance-covariance matrix,

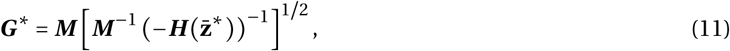

where the operation **X**^1/2^ denotes the square root of **X** such that all the eigenvalues of **X**^1/2^ are positive (Bhatia, 2015; see also eq. 21c of Lande, 1980).

#### 3.4.3 Connections with notions of stability from invasion analyses

Using a quantitative genetics approach, we have derived the conditions under which the multivariate phenotypic distribution of a dispersal limited population converges and remains at an equilibrium (eqs. 8-11). Here, we highlight the connections between these conditions and notions of evolutionary stability that have emerged from invasion analyses and adaptive dynamics under limited dispersal.

##### Singular strategy

First, the selection gradient eq. (6) substituted into condition (8) is equivalent to the definition of evolutionarily singular strategies/phenotypes under limited dispersal (i.e., phenotypes which when expressed by the whole population, the gradient of invasion fitness is zero, e.g., Rousset, 2004; see also Geritz et al., 1998, for general definition).

##### Convergence stability

Second, the condition for a mean trait vector to be an attractor of directional selection (condition 9 with eq. 6) is equivalent to the condition for a multi-trait phenotype to be convergence stable in invasion analysis (Mullon et al., 2016; see also Brown and Taylor, 2010, for a graphical approach to the coevolution of two traits in a genetically structured population; and Leimar, 2009, Geritz et al., 2016, for general considerations on multi-trait in invasion analysis). It is noteworthy that in spite of this equivalence, the phenotypic dynamics envisaged by a quantitative genetics model (given by eq. 3a, see also eq. 7 of Lande, 1979, or eq. 1 of Phillips and Arnold, 1989) differ from the dynamics inferred from invasion analysis (which are captured by the so-called “canonical equation”, eq. 1 of Dieckmann and Law, 1996, or eq. 3 of Leimar, 2009). In a quantitative genetics model, the mean trait vector changes as a result of selection acting on a standing genetic variation, which is large enough to be captured by a statistical distribution (***G***_*t*_ in eq. 3a). Under the “canonical equation”, traits evolve under a trait substitution sequence, whereby a selected mutant fixes before another mutant arises, so that the population “jumps” from one monomorphic state to another and in principle cannot sustain polymorphism (see Fig. 1c, upper right panel of Champagnat et al., 2006, for a useful depiction of a trait substitution sequence; see Van Cleve, 2015, for a review of trait substitution sequences with kin selection effects).

##### Uninvadability

Third, the condition that 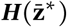 with eq. (7) is negative-definite for the population to remain unimodally distributed around 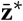 is consistent with the condition derived from invasion analyses for 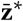 to be locally uninvadable (i.e., that any rare mutant that arises in a population for monomorphic for 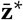 and that causes a slight deviation from 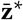 eventually vanishes, Mullon et al., 2016; see also Ajar, 2003 for a single evolving trait in dispersal limited population; and Leimar, 2009, Geritz et al., 2016, for general considerations on multi-trait analyses).

##### Evolutionary branching

Invasion analyses have revealed that a phenotype that is convergence stable is not necessarily uninvadable (Eshel and Motro, 1981, Eshel, 1983, Taylor, 1989, Christiansen, 1991, Abrams et al., 1993b). In fact, when a singular phenotype is convergence stable but invadable, disruptive selection can lead to evolutionary branching, whereby two lineages stably coexist in polymorphism (Metz et al., 1996, Geritz et al., 1998). When multiple traits are evolving, a sufficient condition for the initiation of evolutionary branching is that the Jacobian is negative-definite and the Hessian matrix is positive-definite at the singular phenotype 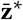 (note that this does not ensure that the resulting polymorphism is stable, Geritz et al., 2016, for further considerations). In the context of quantitative genetics, this means that the mean trait vector is held at 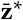 (as 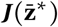 is negative-definite) while the dynamics of the variance-covariance matrix (eq. 3b) diverges to infinity (as 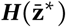 is positive-definite). In other words, at the onset of evolutionary branching, directional selection maintains the population mean vector at 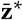 all the while disruptive selection favors extreme phenotypes, leading to the explosion of the variance-covariance matrix (in line with previous quantitative genetics approaches to study evolutionary branching, Wakano and Iwasa, 2013, Débarre et al., 2014, Wakano and Lehmann, 2014, Débarre and Otto, 2016).

#### 3.4.4 The molding of phenotypic correlations by selection and mutation

Invasion analyses can be used to infer on the phenotypic correlations or associations generated by disruptive selection (by studying the eigenvector associated with the greatest eigenvalue of 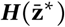, which gives the axis in phenotypic space along which selection is disruptive and along which the population becomes dimorphic, Mullon et al., 2016, Geritz et al., 2016). This approach, however, only incorporates the effect of selection and is limited to studying phenotypic correlations at the onset of evolutionary branching (inferring on the long term outcome of evolutionary branching requires studying invasion in dimorphic populations, which is typically much more involved mathematically, e.g., Geritz et al., 1998, Sasaki and Dieckmann, 2011). A quantitative genetics approach such as ours here allows two further considerations on phenotypic correlations (e.g., Lande, 1980, Jones et al., 2007). First, it allows to incorporate the influence of pleiotropy (through the distribution of mutational input, captured by the variance-covariance matrix ***M*** in eq. 3b). Second, eq. (11) allows to study equilibrium phenotypic correlations as a balance between mutation and stabilizing selection (and not only disruptive selection). We investigate this balance in more detail in the next section.

### 3.5 Application to the coevolution of two synergistic social traits

We now apply the quantitative genetics approach elaborated above to study the coevolution of two social traits under limited dispersal. Our main goal is to illustrate the potential significance of indirect synergistic effects for the molding of phenotypic correlations when dispersal is limited (Figure 1b-d).

#### 3.5.1 Two public goods model

We model the coevolution of two nonnegative quantitative traits, labelled 1 and 2, that each capture participation to a different public good. For examples, in group living mammals, one trait could be the time/energy invested into foraging for the group’s offspring, and the other, investment into defending the group by standing sentry against predators (e.g., Carter et al., 2014); in microorganisms, each trait could be the production of a specific amino-acid that is released into the external environment from which it can then be absorbed and used by group members (e.g., Özkaya et al., 2017).

##### Benefits and costs

We assume that both public goods are shared equally among group members, and that individuals receive extra benefits from obtaining both goods together. The benefits, *B* (**z**_*i*_, **z**_−*i*_), received by a focal individual (with traits **z**_*i*_ in a group composed of **z**_−*i*_) can then be written in terms of the group trait averages 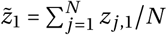 and 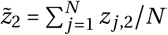 as

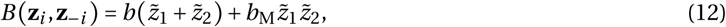

where the parameter *b* > 0 tunes the independent benefit of each public good produced (assumed to be the same for both goods for simplicity); and parameter *b*_M_ >0, the multiplicative benefits of receiving both goods together. Conversely, participation to both public goods simultaneously is assumed to be extra costly, for instance because the different goods call upon different biological functions that are costly to co-maintain, so that the cost *C*(**z**_*i*_) paid by a focal individual (with traits **z**_*i*_) can be written as

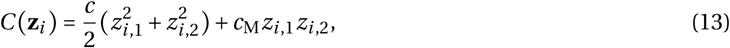

where the parameter *c* > 0 tunes the independent cost of each trait, and parameter *c*_M_ > 0, the multiplicative costs of the traits. The fecundity of a focal individual, *f* (**z**_*i*_, **z**_−*i*_), is then the difference between the benefits received and the costs paid,

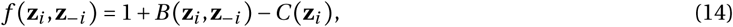

where 1 is the baseline fecundity when no one in the group participates to either public good (*z*_*i*,1_ = *z*_*i*,2_ = 0 for all *i*).

These benefits (eq. 12) and costs (eq. 13) entail that it is best for a focal individual to express a negative *within*-individual association between traits (if expressed at all), and simultaneously be in a group in which traits are positively associated *between*-individuals. Such a configuration is possible when the population is well-mixed (so that there are no genetic correlations – or no relatedness – among individuals of the same group), but difficult when individuals of the same group are related due to limited dispersal. As relatedness increases, associations within- and between-individuals become aligned due to the co-inheritance of linked traits (in fact, the covariance between-traits between-individuals is equal to the product of pairwise relatedness with total covariance in the absence of selection; i.e., the between-individuals covariance of traits *a* and *b* is equal to 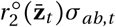, see eq. C-25 in Appendix C). We therefore expect limited dispersal to be relevant to the coevolution of the two traits of our model and to the way selection associates these traits within individuals.

##### Fitness

Before proceeding to the analysis, let us give the individual fitness function of a focal individual 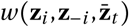. For this model, we assume that there is no group extinction, that offspring disperse independently from one another before local density regulation, and that all adults die after reproduction (so that the population follows a Wright-Fisher life cycle). In this case, individual fitness is,

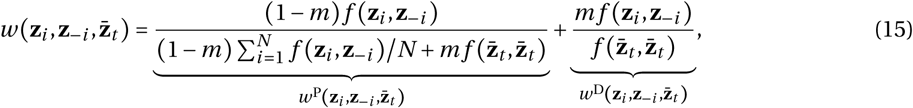

where 0 < *m* ≤ 1 is the dispersal probability. Individual fitness is the addition of two terms: (1) the expected number of offspring of the focal that successfully establish in their natal group, 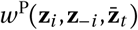, which is the ratio of the number of philopatric offspring of the focal to the total number of offspring that enter the competition in the focal group; and (2) the expected number of offspring of the focal that successfully settle in other groups, 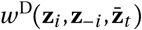, which is the ratio of offspring the focal sends in a non-focal group to the expected number of offspring in such a group (fitness function of the form eq. 15 is standard under limited dispersal, e.g., Rousset, 2004, Ohtsuki, 2010).

##### Relatedness

The final pieces of information that are necessary to apply our framework are the neutral relatedness coefficients, 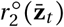 and 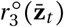, and the effect of each trait on pairwise relatedness 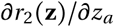. These expressions, which have been derived elsewhere for the Wright-Fisher life-cycle considered here (e.g., Rousset, 2004, Ajar, 2003, Ohtsuki, 2010, Wakano and Lehmann, 2014), are given in Appendix B.2.2 (eqs. B-21-B-22).

#### 3.5.2 Analysis

We now proceed to analyze the evolution of both social traits using the approach established in section 3.2. We first focus on the equilibrium properties of the phenotypic distribution.

##### Convergence of mean trait values

Substituting eq. (15) and pairwise relatedness coefficient (eq. B-21) into eq. (6), we obtain that the selection gradient vector is

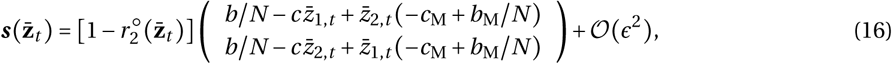

where *ϵ* is a small parameter capturing the magnitude of the effect of the public good on fecundity (i.e., *ϵ* is the largest of the parameters *b, b*_M_, *c, c*_M_). Solving eq. (16) for zero then yields the unique singular strategy

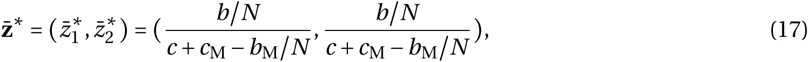

which unsurprisingly decreases with costs *c* and *c*_M_, and increases with “direct” benefits *b*/*N* and *b*_M_/*N* (as an individual recoups a share 1/*N* of its participation to each public good). Note that this singular strategy does not depend on dispersal (or relatedness). This is due our assumptions that group size is fixed and that generations are non-overlapping (in which case indirect fitness benefits of interacting with relatives are “cancelled” by the fitness costs of kin competition, e.g., Taylor, 1992, Rousset, 2004).

To establish whether the phenotypic distribution will converge to have mean 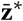, we substitute eq. (16) into the symmetric part of the Jacobian matrix eq. (9), which we evaluate at the equilibrium eq. (17). It is straightforward to show that the two eigenvalues of the resulting matrix are given by

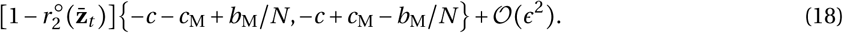

Both are negative provided

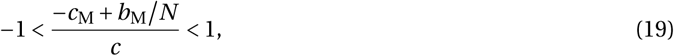

i.e., when the difference between the multiplicative costs, *c*_M_, and direct multiplicative benefits, *b*_M_/*N*, is small compared to the independent cost, *c*. In that case, the population will evolve to have mean given by eq. (17) and produce an equal amount of each public good (Figure 2a). Otherwise, the population will evolve to express a single trait and thus produce a single public good (depending on initial conditions, Figure 2b). Eqs. (17) and (19) reveal that limited dispersal does not influence the evolution of the mean of the phenotypic distribution. But what about the shape of the distribution around this mean?

**Figure 2:**
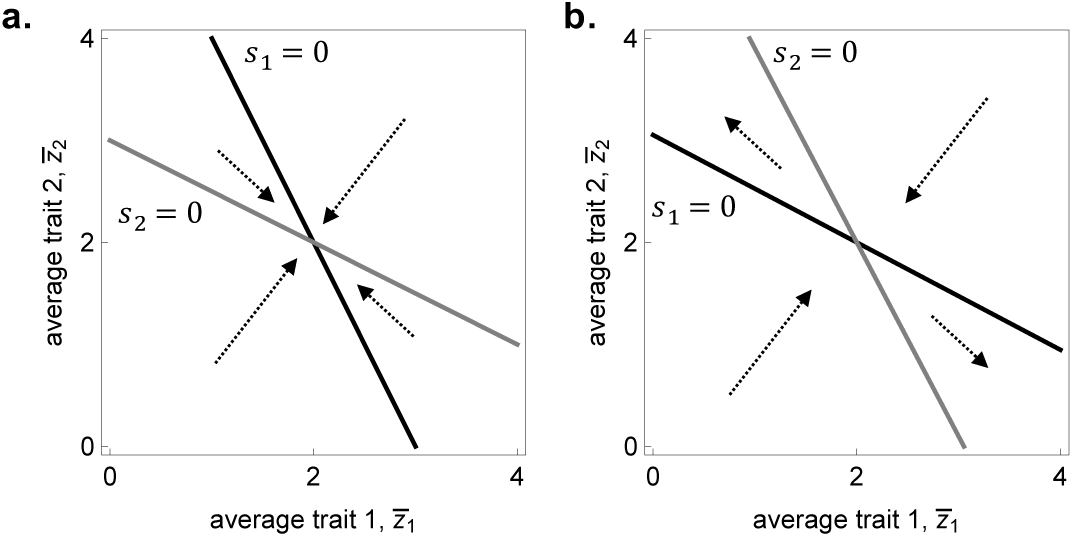
Directional selection on synergistic social traits. Qualitative dynamics of population means due to selection when equilibrium eq. (17) is: **a.** an attractor (with *b*/*N* = 3, *b*_M_/*N* = 1.5); **b.** a repeller (with *b*/*N* = 5.8, *b*_M_/*N* = 0.1). Solid lines show when the selection gradient eq. (16) on each trait vanishes (black for trait 1, 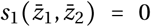; grey for trait 2, 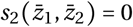; other parameters: *c* =1, *c*_M_ = 2).

##### Stabilisation of the distribution around the mean

Assuming eq. (19) holds true, whether or not the population distribution stabilises around the equilibrium trait values (eq. 17) depends on the Hessian matrix, 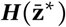. Let us start with analysing the diagonal elements of 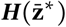, which reveal whether selection on each trait is independently stabilizing or disruptive. Substituting eq. (15) and relatedness coefficients (Appendix B.2.2) into eq. (7) for traits 1 and 2 (i.e., *a* = *b* = 1 and *a* = *b* = 2), and evaluating it at equilibrium eq. (17), we obtain that the diagonal entries of 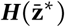 are

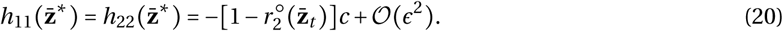

Since 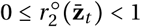, the diagonal entries of 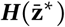 are always negative, which means that selection on each trait is stabilizing when they evolve independently from one another.

Whether selection is stabilizing when both traits co-evolve also depends on the correlational coefficient of selection, 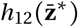. In particular, stabilizing selection requires that: (1) 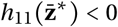 and 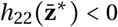; and (2) 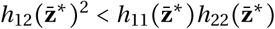, i.e., that the correlational selection coefficient is weak relative to the strength of stabilizing selection on both independent traits; this is because a 2 × 2 symmetric matrix Hessian matrix is negative-definite if and only if its diagonal entries are both negative and the off-diagonal satisfies condition (2) (e.g., Horn and Johnson, 2012). Condition (2) can equivalently be written as

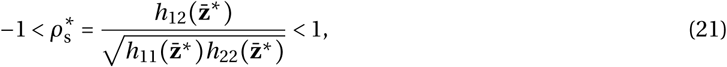

where 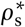 is the strength of correlational selection, relative to the strength of stabilizing selection on each independent trait at 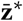. If eq. (21) does not hold, then selection is disruptive due to correlational selection.

The correlational coefficient of selection is derived by first substituting eq. (15) into eq. (7) with *a* = 1 and *b* = 2, and second, evaluating the result at equilibrium eq. (17). This yields

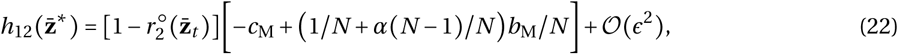

where

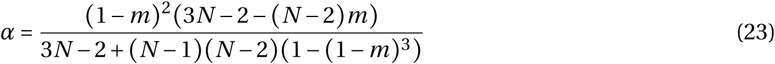

decreases as dispersal and group size increases (i.e., *α* decreases as relatedness coefficients decrease, see Figure 3a). Eq. (22) reveals that as *α* (and relatedness) increases, the within-individual association favored by selection goes from negative to positive (Figure 3b-c). This is because as relatedness increases, indirect synergistic effects become increasingly targeted by correlational selection (Figure 1b-d).

**Figure 3:**
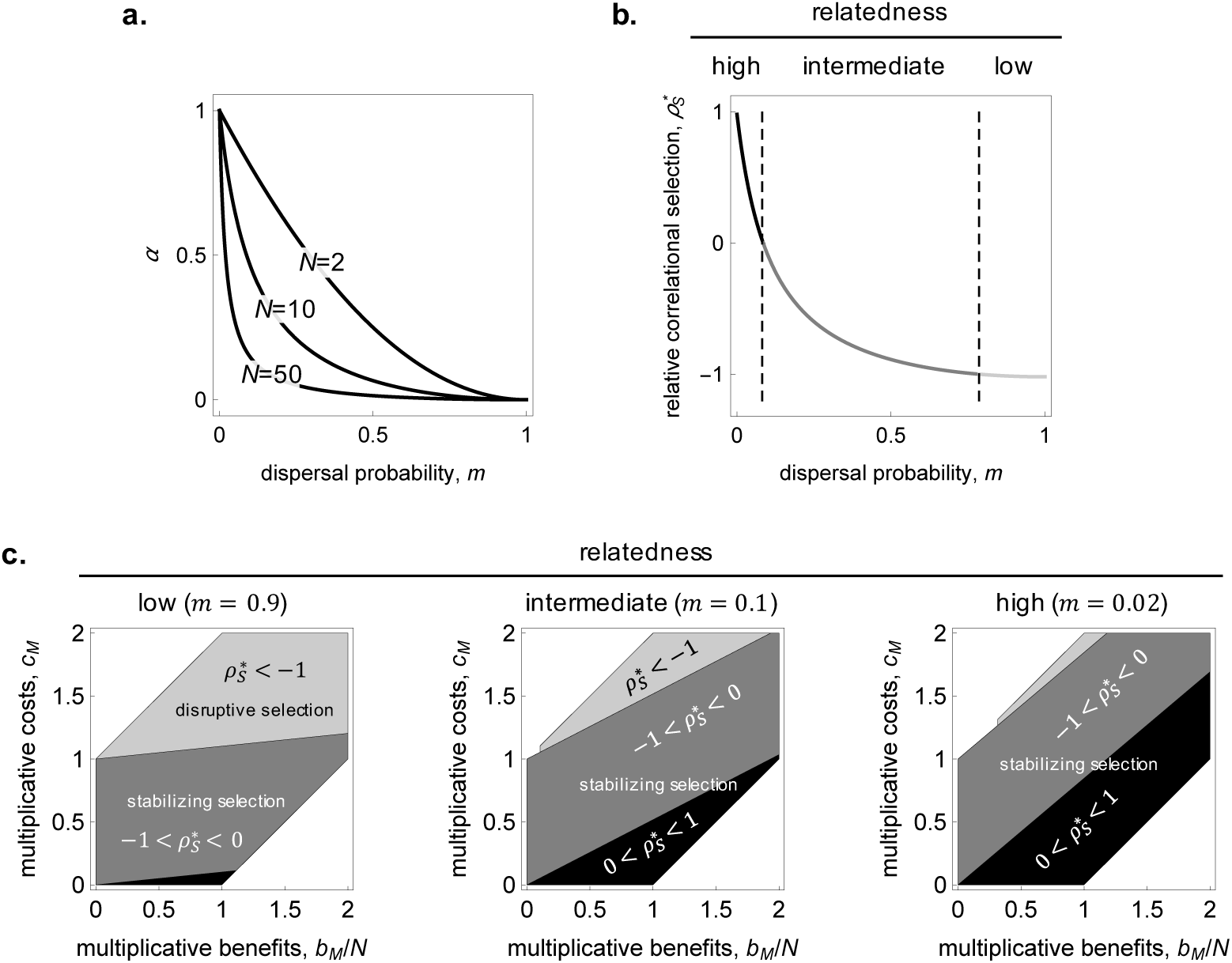
Correlational selection on synergistic social traits. **a.** Weight *α*, eq. (23), to multiplicative benefits in the coefficient of correlational selection (see eq. 22). **b.** Relative correlational selection, 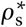 (eq. 24), as a function of dispersal *m*, with critical levels of dispersal for which: 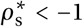 (light grey); 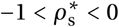 (dark grey); and 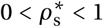 (black, with *N* = 10, *b*/*N* = 0.03, *b*_M_/*N* = 1.8, *c* = 0.8, *c*_M_ = 1). **c.** Parameter combinations (with *N* = 10, *c* = 1) for which correlational selection at the equilibrium eq. (17) is: (1) strongly negative (and causes selection to be disruptive, 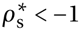 & eq. 24 does not hold, light grey regions); (2) negative (and selection is stabilizing, 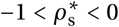 & eq. 24 holds, dark grey regions); and (3) positive (and selection is stabilizing, 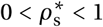 & eq. 24 holds, black regions). White regions correspond to parameter combinations under which the equilibrium is not evolutionary convergent (i.e., eq. 19 does not hold).

Substituting eqs. (20) and (22) into eq. (21), we find that selection is stabilizing around 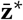 when

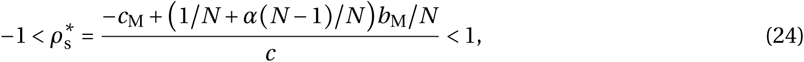

which reveals that high relatedness, or large *α*, favors stabilizing selection (Figure 3b-c, dark grey and black regions), and conversely, low relatedness, or small *α*, favors disruptive selection and thus polymorphism (when eq. 19 holds but eq. 24 does not, Figure 3b-c, light grey region). This finding is in line with a recent computational eco-evolutionary model which found that when species can evolve cross-feeding interactions, mutualistic coexistence is compromised by spatial structure and limited dispersal (Oliveira et al., 2014). This is also in line with previous results on the evolution of single traits that have found that evolutionary branching is inhibited by limited dispersal (e.g., Day, 2001, Ajar, 2003, Wakano and Lehmann, 2014, Parvinen et al., 2017). In such models and ours, limited dispersal inhibits evolutionary branching because it creates genetic correlations among competing individuals, so that a mutant cannot be as different to common types as in well-mixed population. As a result, frequency-dependent disruptive selection is weaker under limited dispersal.

##### Effect of selection on phenotypic correlation

Putting our stability analyses together (especially eqs. 17, 19, 22, and 24) and validating them using individual-based simulations (see Appendix D for details), we find that there are three possible outcomes for the phenotypic distribution once it has converged to be unimodal around the equilibrium eq. (17) due to selection: (1) when relatedness is low, correlational selection is negative and strong enough to make selection disruptive, leading to the stable coexistence of individuals specialized in producing a single public good (Figure 4a). In this instance, evolutionary dynamics follow so-called “Black queen” dynamics (Morris et al., 2012, Morris, 2015, with special reference to microorganisms): individuals first evolve to produce the same amount of leaky product that is shared among individuals, but the costly maintenance of both traits leads to specialization in a single product and the evolution of cross-feeding among types (see Rueffler et al., 2012, Vásárhelyi et al., 2015, for similar models on the evolution of specialization in well-mixed populations). (2) Over a critical level of relatedness, selection becomes stabilizing but correlational selection remains negative, which prevents evolutionary branching and thus specialization, but still results in a negative association among traits within individuals (Figure 4b). (3) Over another threshold of relatedness, correlational selection becomes positive, so that the traits become positively associated within individuals (Figure 4c). Hence, even though limited dispersal and relatedness have no bearing on the mean of the phenotypic distribution in our model (eqs. 17 and 19), indirect synergistic effects entail that relatedness has a significant influence on the shape of this distribution (which goes from being bimodal with a negative correlation under panmixia to unimodal with a positive correlation under limited dispersal, Figure 4).

**Figure 4:**
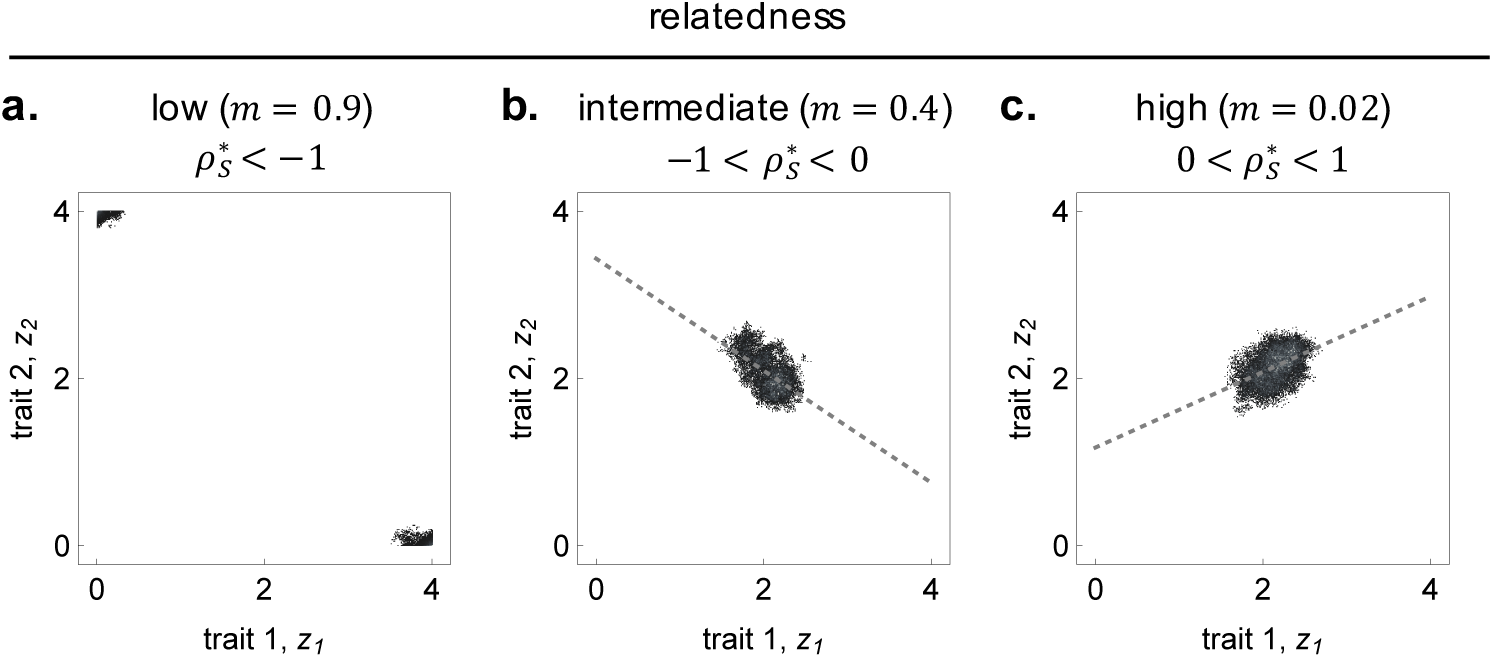
The effect of relatedness and indirect synergy on the phenotypic distribution. Equilibrium phenotypic density distribution, *p*_*t*_(**z**), of a simulated population, initially monomorphic for both traits at equilibrium 2, 2 (population composed of 1000 groups of size *N* = 10; sampled every 500 generations for 20’000 generations after 30’000 generations of evolution; other parameters: 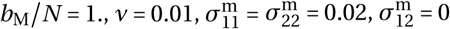; see Appendix D for details on simulations). **a**. Strong negative association with social polymorphism (with *b*/*N* = 0.2, *c* = 0.1, *c*_M_ = 1). **b**. Negative association (correlation = −0.67, *p* < 10^−10^; with *b*/*N* = 2.2, *c* = 1, *c*_M_ = 1.1). **c**. Positive association (correlation = 0.45, *p* < 10^−10;^ with *b*/*N* = 0.1, *c* = 1, *c*_M_ = .05)

##### Effect of pleiotropy on phenotypic correlation

So far, our analysis has focused on the effects of selection on the stability of jointly evolving traits (an analysis that could have equally been performed using invasion analysis; see Mullon et al., 2016, for such an approach to the joint evolution of multiple traits under limited dispersal). But selection is not the only relevant process for the way phenotypic distributions are shaped. As highlighted by the present quantitative genetic approach, the equilibrium variance-covariance matrix of the phenotypic distributions also depends on the patterns of mutation (captured by matrix ***M*** in eq. 11). In particular, pleiotropy is expected to influence the correlations among traits within individuals at an evolutionary equilibrium.

In order to investigate the joint effects of pleiotropy and correlational selection, let us assume that the variance of mutational effect on both traits is the same 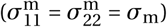, in which case the variance-covariance matrix of mutation effects can be written as

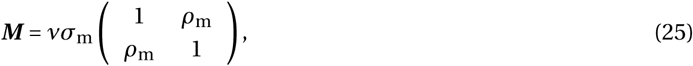

where 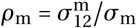 is the correlation of the effect of mutations on traits 1 and 2. The parameter −1 < *ρ*_m_ < 1 thus captures the degree of pleiotropy between both traits (when it is zero, both traits change independently due to mutation, when it is positive, they tend to change in similar ways, and when it is negative, in opposite ways).

Substituting eqs. (20), (22) and (25) into eq. (11), we find that the correlation 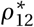 among traits 1 and 2 at equilibrium is

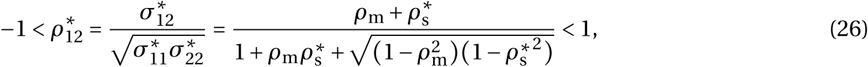

where 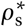 is given in eq. (21). This shows that at equilibrium, the sign of the correlation among between two traits reflects the balance, 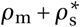, between the degree of pleiotropy, *ρ*_m_, and the relative strength of correlational selection 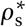 (see Figure 5a; note that eq. 26 can be directly deduced from eq. 11 whenever the variance of mutational effect on both traits is the same, 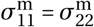, and the coefficients of disruptive selection on independent traits are equal, 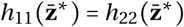, see eq. 8 of Jones et al., 2007). Since limited dispersal and relatedness has a significant influence on relative correlational selection 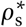 (eq. 21), it can affect the correlation 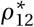 among traits in the population as much as pleiotropy, *ρ*_m_.

**Figure 5:**
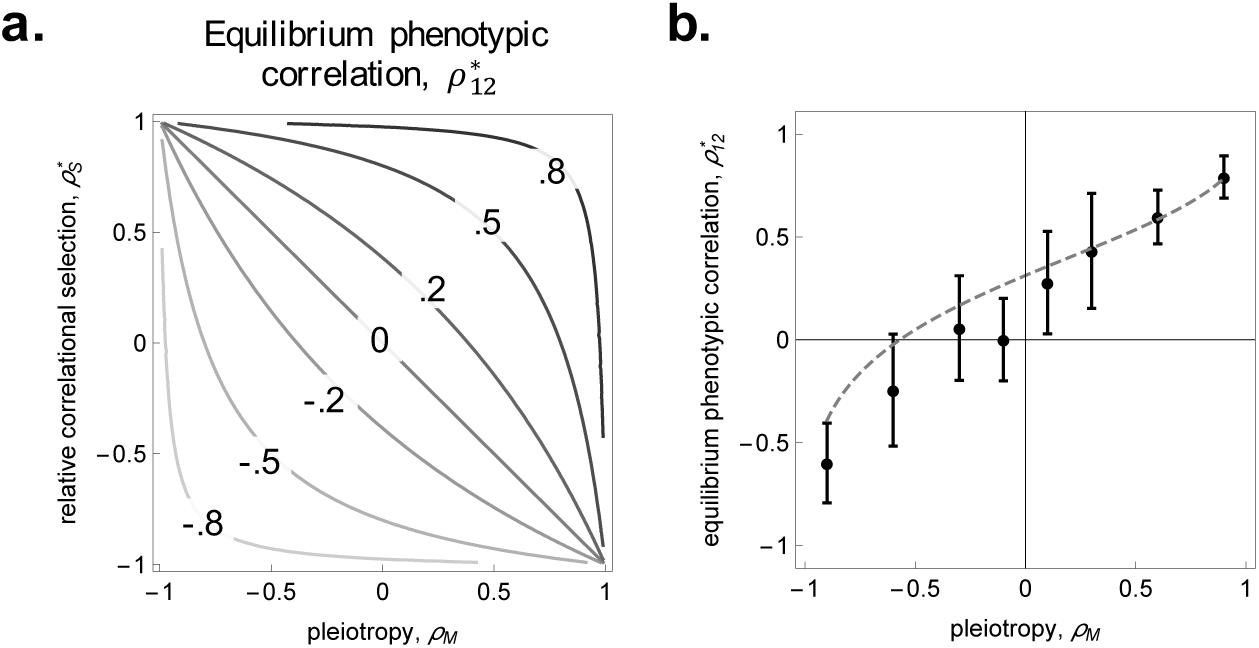
The effect of pleiotropy on phenotypic correlation. **a**. Contours of predicted phenotypic correlation among traits 1 and 2 at mutation-selection balance, 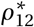, according to pleiotropy, *ρ*_m_, and the relative strength of correlational selection, 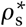 (from eq. 26). **b**. Predicted phenotypic correlation among traits 1 and 2 (dashed grey curve, from eq. 26), and corresponding observations from individual based simulations of a population initially monomorphic for 2, 2 divided among 1000 groups of size *N* = 10 (black dots, averaged correlation over 20’000 generations after 30’000 generations of evolution, error bars indicate standard deviation; other parameters: *m* = 0.05, *b*/*N* = 0.2, *b*_M_/*N* = 1, *c* = 1, *c*_M_ = 0.1, *v* = 0.01, 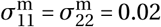; see Appendix D for details).

We additionally checked that our model captured pleiotropy correctly by comparing the phenotypic correlation among the two traits at equilibrium predicted by our model (eq. 26) and that observed in simulations for different levels of pleiotropy. We found that model predictions and observations from simulations also matched well in the presence of pleiotropy (Figure 5b).

##### Dynamics of the distribution

We further tested the accuracy of our dynamical model by comparing individual-based simulations with numerical recursions of eqs. (3). We found that simulated populations tend to have lower phenotypic variance than eqs. (3) would predict (Figure 6). This is probably due to global genetic drift, which our model ignores and which depletes phenotypic variance (as in well-mixed populations, e.g., Wakano and Iwasa, 2013, Débarre and Otto, 2016), and/or the presence of phenotypic skew, which is ignored under our assumption that the phenotypic distribution in the population is normal, but which can influence the dynamics of phenotypic variance (Appendix B, eq. B-18). Nonetheless, we observed overall a good qualitative fit between the predicted and observed dynamics of the phenotypic distribution (Figure 6). This suggests that the assumption of normality yields accurate predictions for the change of mean and variance when dispersal is limited (like in well-mixed populations, Turelli and Barton, 1994).

**Figure 6:**
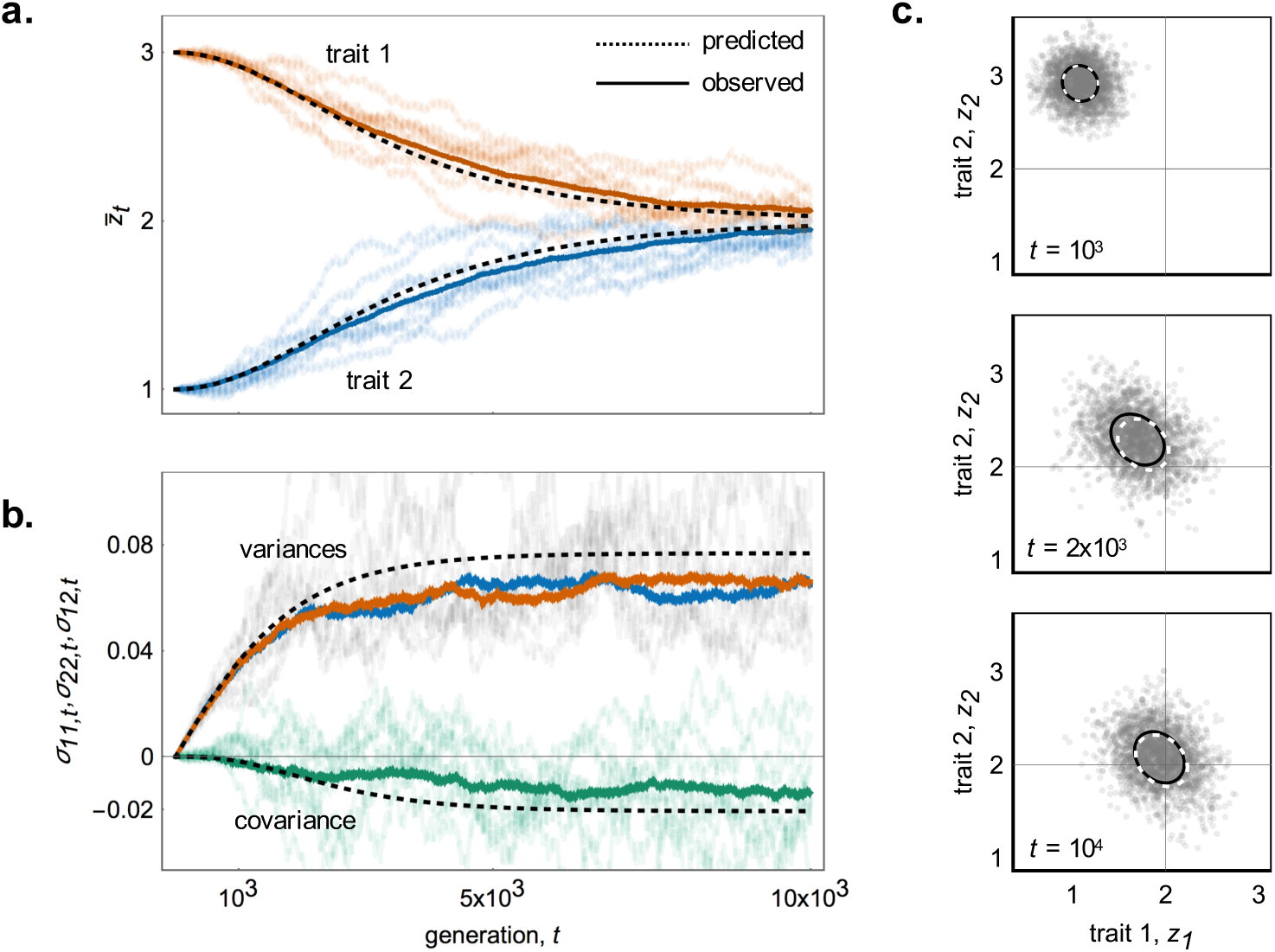
Observed and predicted evolution of the phenotypic distribution, *p*_*t*_ (z). The observed (full lines, from individual based simulations) and predicted (dashed lines, from eq. 3) evolution of the traits’ means (**a**. trait 1 in orange and 2 in blue), variances (**b**. trait 1 in orange and 2 in blue) and covariance (**b**. green) for 64 replicates (10 randomly chosen replicates in lighter shade, average over all 64 replicates in darker shade, initial population monomorphic with *z*_1_ = 3 and *z*_2_ = 1, distributed over 1000 groups of size *N* = 10, other parameters: *m* = 0.4, *b*/*N* 14.8, *b*_M_/*N* = 0.1, *c* = 5, *c*_M_ = 2.5, *v* = 0.1, 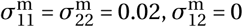; see Appendix D for details). **c**. Snapshot of the population (2’500 individuals randomly sampled across 64 replicates shown by grey points) and variance-covariance ellipses given by the (right) eigenvectors of the **G** matrix (observed across all 64 replicates in full lines and predicted in dashed), at generations: 1’000 (top panel); 2’000 (middle panel); and 10’000 (bottom panel).

## 4 Discussion

In this paper, we have modeled the evolution of the distribution of genetically-determined quantitative traits under limited dispersal, frequency-dependent selection and pleiotropic mutation. By doing so, we have generalized two classical quantitative genetics results to include limited dispersal: first for the general recurrence eq. (1) of the phenotypic distribution under the continuum of alleles model (Kimura, 1965b, Fleming, 1979, Lande, 1979, Bürger, 1986); and second for the closed dynamical system eq. (3) of the vector of means and matrix of variance-covariance when the distribution is normal (Lande, 1979, Lande and Arnold, 1983, Phillips and Arnold, 1989). In both cases, genetic structure due to limited dispersal leads to the replacement of individual fitness in classical quantitative genetics equations by lineage fitness. This is the fitness of a typical carrier of a given phenotype (randomly sampled from the lineage of all members carrying that phenotype), i.e., the average direct fitness of a phenotype, which depends on the phenotypes expressed in the whole population and how they are distributed among groups (eq. 2).

From lineage fitness, we were able to reinforce existing links between concepts of evolutionary stability and evolutionary quantitative genetics: (1) the vector of means evolves to convergence stable phenotypic values (eqs. 8-9; see Charlesworth, 1990, Iwasa et al., 1991, Taper and Case, 1992, Abrams et al., 1993a, Abrams, 2001, Lion, 2018, for well-mixed populations, and Cheverud, 1985, Queller, 199 2a,b, Frank, 1998, McGlothlin et al., 2014, Lehmann and Rousset, 2014, for limited dispersal); and (2) the distribution remains unimodal around such values when they are locally uninvadable or may become bimodal when they are invadable (eq. 10; see Sasaki and Dieckmann, 2011, Wakano and Iwasa, 2013, Débarre et al., 2014, Débarre and Otto, 2016, for well-mixed populations, and Lehmann and Rousset, 2014, Wakano and Lehmann, 2014, for the dynamics of the variance of a single trait around a singular strategy under limited dispersal). Specifically, we have shown that in a dispersal-limited population with infinitely many types, the selection gradient, which determines the change in mean trait values (eq. 3a), and the Hessian matrix, which shapes the variance-covariance matrix (eq. 3b), are respectively equal to the selection gradient vector and Hessian matrix computed from the invasion fitness of a rare mutant in an otherwise monomorphic population (i.e., eqs. 6-7 are equal to eqs. 12-13 of Mullon et al., 2016. Since the correspondence between the selection gradients of the two approaches is well-established for dispersal-limited populations (Lehmann and Rousset, 2014, for review), it may be felt that the correspondence between the Hessian matrix obtained from the evolutionary invasion analysis and that determining the change in the variance-covariance matrix is intuitive and must hold. Demonstrating this, however, required surprisingly lengthy calculations (see Appendix) showing that it is actually not obvious that when many alleles segregate in a dispersal-limited population and traits are far away from a convergence stable point, selection on phenotypic (co)variances only depends on simple pairwise and three-way probabilities of identity-by-descent. Having established these relationships, we expect them to hold under more general demographic settings (e.g., with local demographic fluctuations) and hope that simpler arguments than our present ones can be found to prove it.

The extension of evolutionary invasion analyses to a quantitative genetics model allows to specify the phenotypic distribution at a mutation-selection balance (eq. 11). In particular, it allows to study the effects of selection and mutation on the phenotypic associations that emerge among traits at equilibrium (eq. 26). Our analyses of such associations suggest that kin selection due to limited dispersal can mold phenotypic associations as much as pleiotropic mutations (eq. 26 and Fig. 5). By expressing correlational selection on traits in terms of their direct and indirect fitness effects, we gained insights into the influence of kin selection on phenotypic associations (eq. 7). Motivated by our explicit formula for the variance-covariance matrix (eq. 11) and our example (section 3.5), we complement here the discussion found in Mullon et al. (2016) (based on an invasion analysis) on the implications of kin selection for the evolution of within-individual phenotypic associations. As indicated by the decomposition of correlational selection eq. (7a), there are two ways kin selection influences such associations.

The first is through the fitness effects that traits have when co-expressed among relatives, so when traits have indirect synergistic effects (eq. 7b, Figure 1b-d). Under limited dispersal, selection favors an association among two traits within individuals, when such an association between individuals has indirect fitness benefits. Due to such kin selection effects, different levels of dispersal can lead to significantly different evolutionary outcomes for phenotypic associations, as highlighted by our example on the coevolution of two traits whose association within-individual is costly but beneficial between-individuals due to social synergy. In this example, populations with little genetic structure evolved a division of social labor, with individuals specialized in only one trait coexisting with one another (Figure 4a), but populations with strong genetic structure evolved no such specialization, with traits in fact becoming positively associated within individuals (Figure 4c). In line with our results, populations of *E. coli* that experience frequent mixing (so show little genetic structure) readily evolve cross feeding interactions in poor environments, so that different strains specialize in the production of a specific amino acid (D’Souza and Kost, 2016). By contrast, in meerkat social groups (which typically show high levels of relatedness), individuals tend to participate to all social activities, with participation to different tasks such as babysitting and pup feeding positively associated within individuals (Clutton-Brock et al., 2003). Such patterns can be explained by our results if participation to different tasks in these systems is genetically determined, at least partially.

It is also worthy of note that in our example, relatedness has a substantial influence on the shape of the phenotypic distribution but none on the mean of this distribution (Figure 4, eqs. 17 and 18). Hence, the effects of genetic structure on phenotypic evolution that we report would have gone unnoticed from the study of the dynamics of the mean only (which is the focus of the vast majority of study of quantitative genetics in family-structured populations, e.g., Cheverud, 1985, Queller, 199 2a,b, Frank, 1998, McGlothlin et al., 2014), or from the analysis of the selection gradient vector only (as done in the majority of evolutionary analyses to synergistic social traits, e.g., Gandon, 1999, Perrin and Mazalov, 2000, Reuter and Keller, 2001, Lehmann and Perrin, 2002, Rousset and Gandon, 2002, Gardner and West, 2004, Leturque and Rousset, 2004, Hochberg et al., 2008, Brown and Taylor, 2010, Kuijper and Johnstone, 2017). Overall, our example highlights that when traits have indirect synergistic effects between individuals (Figure 1b-d), relatedness is important for the way natural selection molds phenotypic associations within individuals. This consideration should be especially relevant to the evolution of specialization and the emergence of division of labor.

A relevant pair of traits likely to be influenced by such kin selection effects is costly helping and punishment, which have synergistic indirect benefits when expressed by different individuals (e.g., Raihani et al., 2012, and references therein). According to our results, kin selection should favor a positive association among helping and punishment, which interestingly, has been observed in humans (Fehr and Gächter, 2000). Another pair of traits whose evolution is likely to be influenced by their joint expression in different individuals is the production and exploitation of a public good, such as the secretion and use of siderophores by microorganisms (West et al., 2006). Under limited diffusion of siderophores and limited bacterial dispersal (Nadell et al., 2009, Kümmerli et al., 2014, Ross-Gillespie et al., 2015), we expect kin selection effects to be ecologically relevant for how secretion and use of siderophores are associated, and more generally for patterns of multi-trait social variation within microbial communities (Cordero and Polz, 2014, van Gestel et al., 2015, Özkaya et al., 2017, Schiessl et al., 2019, RodrÍguez Amor and Dal Bello, 2019).

The second way kin selection influences phenotypic associations is via the combination of the indirect effect of one trait with the effect of the other on the tendency to interact with relatives (“synergy via relatedness”, eq. 7c, Figure 1e). Specifically, selection favors an association among two traits when it results in fitness benefits being preferentially directed towards relatives or fitness costs towards non-relatives. For example, if trait *a* has positive indirect fitness effects (e.g., altruistic helping) and trait *b* decreases the tendency to interact with relatives (e.g., dispersal), then selection favors a negative correlation between traits *a* and *b* (e.g., Koella, 2000, Purcell et al., 2012, Mullon et al., 2018). We refer readers interested in this effect to Mullon et al. (2016), in which it is discussed at greater length, in particular in the context of dispersal syndromes (Edelaar and Bolnick, 2012, Ronce and Clobert, 2012).

More generally, our evolutionary perspective on phenotypic associations may be useful to empiricists who investigate correlational selection among traits in experimental or natural populations (e.g., Blows and Brooks, 2003, Blows, 2007, for reviews, and ch. 30 of Walsh and Lynch, 2018). Based on Lande and Arnold (1983) paper, the typical starting point of such studies is to perform a quadratic regression of individual fitness on the multiple traits expressed by this individual (for e.g., eq. 30.11 of Walsh and Lynch, 2018). The linear regression coefficients are collected in a vector usually denoted ***β*** with entry *β*_*a*_ interpreted as directional selection on trait *a*, and the quadratic coefficients in a matrix ***γ*** with entry *γab* interpreted as correlational selection on traits *a* and *b* (in our notation, 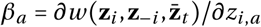 and 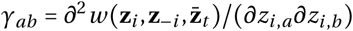). This correspondence between selection on traits and regression coefficients on individual fitness, however, is only valid in well-mixed populations. Indeed, as our analysis has shown, ***β*** and ***γ*** are respectively equal to the selection gradient 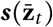 and Hessian matrix 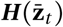, only when all relatedness coefficients are zero (eq. 6 and 7).

For populations that are genetically structured, empirical estimates of selection on multiple traits require to: first regress individual fitness on the traits of the focal individual and on those of its social partners; and second, weigh these indirect fitness effects by relatedness coefficients (according to eqs. 6 and 7). Estimates of pairwise relatedness can be obtained from *F*_ST_ statistics at neutral sites such as microsatellite loci (i.e., the genetic covariance among pairs of interacting individuals relative to the average genetic covariance in the population, Charlesworth and Charlesworth, 2010, chapter 7.1). Similarly, three-way relatedness coefficients can be estimated from comparisons between the genetic skew at neutral sites among triplets of interacting individuals and the average genetic skew in the population. While the importance of indirect fitness effects and relatedness has long been emphasized for the directional selection gradient (so considering only linear regression coefficients, ***β***, e.g., Cheverud, 1985, Queller, 1992a, b, Frank, 1998, McGlothlin et al., 2014, see also ch. 5 of Walsh and Lynch, 2018), our analysis has further quantified the relationship between correlational selection and quadratic regression coefficients (***γ***, eq. 7, Figure 1b-d), which is necessary to understand patterns of phenotypic variation within populations.

In practice, it is likely to be challenging to obtain reliable estimates of all the quadratic regression coefficients necessary to quantify the strength and direction of correlational selection (eq. 7). But our results can nevertheless be of use when designing experimental assays or interpreting collected data. For instance, our results show that for traits that underlie social or competitive behaviors, such as mating, aggression or cooperation, there is little reason to believe that a quadratic regression of an individual’s fitness on its own traits provides a full picture of correlational selection. A corollary to this is that when there is mismatch between phenotypic correlations among two traits observed in a population on one hand, and the quadratic regression coefficient on individual fitness from experimental assays on the other (e.g., Bell and Sih, 2007, Adriaenssens and Johnsson, 2012, Han and Brooks, 2013, Akçay et al., 2015), this may indicate the presence of indirect synergistic fitness effects among traits and genetic structure in the population (rather than genetic constraints as typically inferred). One first accessible step towards testing this hypothesis would be to estimate genetic relatedness among interacting individuals. A high relatedness would suggest that phenotypic correlations are influenced by indirect synergistic fitness effects, which could then be estimated though quadratic fitness regressions of fitness on partners’ traits.

Our results further provide insight into the effects of limited dispersal on how selection influences the **G** matrix of additive genetic variances-covariances (Steppan et al., 2002, Arnold et al., 2008). Previous theoretical works have studied how linkage disequilibrium, pleiotropy and epistasis influence **G** under selection (Lande, 1980, 1984, Turelli, 1985, Turelli and Barton, 1990, Revell, 2007, Jones et al., 2014), but the effects of limited dispersal on **G** have either been assessed in the absence of selection (Lande, 1992), or when selection is frequency-independent (Jones et al., 2004, Guillaume and Whitlock, 2007, Guillaume, 2011, Björklund and Gustafsson, 2015). Here, we have shown that kin selection effects due to limited dispersal are relevant for the way selection favors phenotypic associations (i.e., for correlational selection, eq. 7), which in the long run can lead to genetic correlations through genetic integration (Sinervo and Svensson, 2002, Roff and Fairbairn, 2012). Of course, our model ignores many relevant features for quantitative genetics: environmental effects, genetic dominance, genetic linkage or sexual reproduction for examples. In particular, by assuming that individuals are haploid and reproduce clonally, our model does not allow to distinguish between different possible genetic architectures such as one pleiotropic locus that determines all evolving traits *versus* one independent locus for each trait. In the latter case, correlations among traits in dispersal-limited sexuals would depend on epistatic effects between loci within individuals and genetic linkage between loci (like in well mixed populations, Lande, 1984), as well as epistatic effects between loci located in different individuals, weighted by the genetic associations between these loci within groups (which would depend on dispersal, inbreeding and linkage; see Roze and Rousset, 2008 for how to compute such associations). Incorporating these features into our framework is likely to make the analysis of selection more complicated, but it would allow to study the genetic basis of variation, such as how genetic architecture and its evolution influence trait associations (e.g., Saltz et al., 2017).

One other significant limitation to our present approach is that it assumes that the phenotypic distribution is normal. This assumption is likely to be violated under frequency-dependent selection, which can lead to skewed and complicated distributions. In particular, the normal assumption precludes investigating what happens to the phenotypic distribution once evolutionary branching has occurred (like adaptive dynamics models based on the invasion analyses of monomorphic populations). To relax this assumption would entail tracking the dynamics of higher moments of the phenotypic distribution. One possible way to retain some mathematical tractability would be to use the oligomorphic approximation proposed by Sasaki and Dieckmann (2011). This approximation decomposes a multimodal trait distribution into a sum of unimodal distributions, each corresponding to a morph. Applying Sasaki and Dieckmann (2011)’s approach, which was developed for a large and well-mixed population, to a dispersal limited one, would be an interesting avenue of future research, as well as including class-structure (e.g., age- or sex-structure).

To conclude, we have derived a quantitative genetics model to study the gradual evolution of multiple traits that experience frequency-dependent selection and pleiotropic mutations when dispersal is limited. This model has revealed that limited dispersal opens previously unattended pathways for correlational selection, through the synergistic effects of traits: (1) between interacting individuals (Figure 1b-d), due to non-random frequency-dependent interactions; and (2) via relatedness (Figure 1e), owing to preferential interactions with relatives. This suggests that limited dispersal can profoundly influence how associations between social traits emerge in response to mutation and selection. Given the ubiquity of genetic structure in natural populations (e.g., Bohonak, 1999, Charlesworth and Charlesworth, 2010, p. 310), our results can help understand a wide range of patterns of intra-specific variation in competitive or social traits (such as behavioral syndromes, Dall et al., 2004, Dingemanse et al., 2012; social niche specialization, Bergmüller and Taborsky, 2007, Montiglio et al., 2013; or social division of labour, Boehm, 2002, Wright et al., 2014), which are increasingly thought to be ecologically significant (Bolnick et al., 2011, Wolf and Weissing, 2012, Sih et al., 2012, Canestrelli et al., 2016, Chaturvedi et al., 2017, Estrela et al., 2019). More broadly, by connecting different branches of theoretical evolutionary biology, from invasion analysis to adaptive dynamics to quantitative genetics, the present framework further bolsters the notion that whatever modeling approach is taken, natural selection cannot be divorced from kin selection when dispersal is limited (Hamilton, 1964, Frank, 1998, Rousset, 2004, van Baalen M, 2013, Lehmann et al., 2016).

## Appendix

## A Phenotypic distribution dynamics

In this appendix, we derive eq. (1) of the main text.

### A.1 Process construction

We first lay the foundations of our analysis by describing how phenotypic evolution in our model population (see section 2) is represented mathematically.

#### A1.1 Markov chain

The phenotypic state, or state for short, of a group at given time point is given by the set of phenotypic values of all individuals residing in that group, {**z**_1_, …, **z**_***N***_}(where **z**_*i*_ **=** {*z*_*i*,1_, …, *z*_*i,n*_}ϵ ℝ^*n*^ is the phenotype of individual indexed *i* ϵ {1, …, *N*}). The state of each group in the population changes stochastically from one time period to the next (i.e., after one iteration of the life cycle) due to selection, mutation and dispersal. We assume that these changes can be modeled as a discrete time Markov chain on a continuous state space (as traits are continuous; see Meyn and Tweedie, 2009, for general state spaces). Because groups affect one another through dispersal, the transition kernel of a group depends on the state of all the other groups. But since there is an infinite number of groups and there is no isolation-by-distance (i.e., all groups are equally connected to one another through dispersal), the infinite set of interacting Markov chains (one for each group) can be described as a single Markov chain (for a focal group), whose kernel is a function of the expected value of the process (see Chesson, 1981, 1984, for ecological models). In other words, we can focus on the stochastic dynamics of a focal group and ignore the stochasticity stemming from groups other than the focal one.

#### A.1.2 Markov chain in terms of counting measures

Note that to describe the state of a focal group, the order of elements in {**z**_1_, …, **z**_*N*_} does not matter (because there is no class structure in our population, we do not care which specific individual carries a given phenotype within a group). What matters is how many individuals carry which phenotypes. We can thus represent the state of the focal group by a function, a *counting measure*, that counts the number of individuals with phenotypes that belong to an arbitrary set. Specifically, the counting measure, denoted *µ*, takes a subset *E* ⊆ ℝ^*n*^ and sends it to a non-negative integer by counting the number of individuals within the focal group with phenotypes that belong to *E* according to the following definition

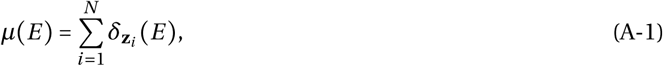

where *δ* is the dirac measure,

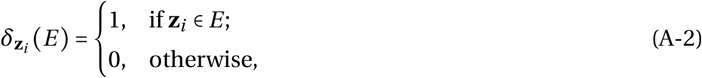

(p. 51 of Harris, 1963, p. 3 of Daley and Vere-Jones, 2003; see also pp. 228-229 of Bürger and Bomze, 1996 for further formal considerations on using counting measures to describe a population under the continuum-of-alleles model). Applied to a single phenotypic value **z =** {*z*_1_, …, *z*_*n*_} ϵ ℝ^*n*^, where *z*_*a*_ is the value of trait *a, µ*(**z**) returns the number of individuals with phenotype **z**, and applied to the whole set of possible phenotypic values, it returns group size, *µ* (ℝ ^*n*^) = *N*.

Under definition eq. (A-1), each possible state that a group can assume is uniquely determined by a specific counting measure, i.e., for each unordered set of *N* vectors in ℝ^*n*^, there exists a unique counting measure (p. 52 of Harris, 1963 and p. 7 of Daley and Vere-Jones, 2003). We can therefore study the dynamics of the state of the focal group by studying the dynamics of its equivalent counting measure (Daley and Vere-Jones, 2003, p. 13-15). So, if *S*_*t*_ denotes the (random) counting measure of a focal group at time period *t*, we can study the Markov chain {*S*_*t*_}on the space of all finite counting measures, which we write as 𝒮. This type of construction has so far primarily been used to study phenotypic evolution in populations that are well-mixed and when time is continuous (e.g., Bürger and Bomze, 1996, Oechssler and Riedel, 2001, Champagnat et al., 2006, Champagnat and Lambert, 2007; but see Morale et al., 2005, Simon, 2008, for populations in explicit space).

#### A.1.3 State dynamics

To describe the stochastic dynamics of the state of a focal group, we let

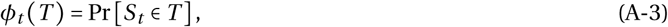

denote the probability that a focal group is in a state that belongs to a subset *T* ⊆ 𝒮 at time period *t* (equivalent to eq. 3.3 of Harris, 1963, p. 55). Since there is an infinite number of groups, *ϕ*_*t*_ (*T*) also gives the distribution of group states in the whole population. The dynamics *ϕ*_*t*_*(T*) are governed by the Markov kernel transition function,

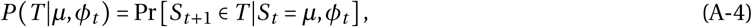

which is the probability that a group will be in a state that belongs to a subset *T* ⊆ 𝒮 at time period *t* +1, given that it was in state *µ* at time period *t* and that the population distribution of states is *ϕ*_*t*_ (this is a non-homogeneous Markov chain, eq. 6.1 of Harris, 1963, p. 60).

State dynamics, or the probability that the focal group is in a state that belongs to *T* ⊆ 𝒮 at time period *t* + 1, can then be written as

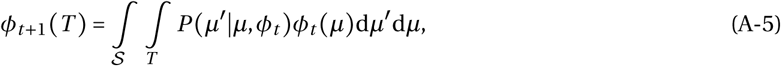

i.e., the sum of weighted probabilities of going from all states *µ* ϵ 𝒮 to states *µ′* ϵ *T*. Because one iteration of the life cycle (from *t* to *t*+ 1) encompasses many events, like selection, mutation, and dispersal, the transition kernel for our model is difficult to characterize (studies like Champagnat et al., 2006, are capable of constructing explicit transition kernels by considering time steps small enough so that only one event can occur per step). To model the evolutionary process in a more practical way, we will focus on the dynamics of the distribution of phenotypes across the entire population rather than on the dynamics of the distribution of group states *ϕ*_*t*_ (*T*).

### A.2 Recurrence for the phenotypic distribution

The distribution of phenotypes across the entire population at time *t* is given by the density function

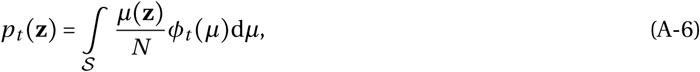

where *µ*(**z)/***N* is the frequency of individuals with phenotype **z** within a group in state *µ* (recall that all groups have the same size *N*). Using eq. (A-5), the phenotypic distribution at time period *t* +1 can be written as

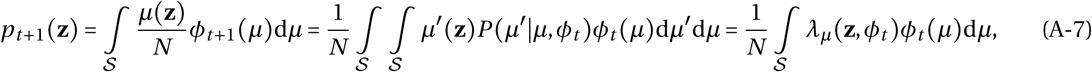

where

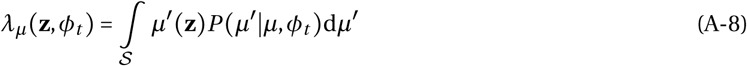

is the expected number of individuals with phenotype **z** residing in a focal group at time *t* +1, given that this focal group was in state *µ* at time *t* (and the population state distribution was *ϕ*_*t*_). We can decompose this expected number as

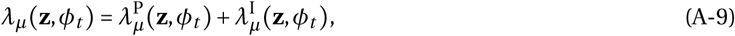

where 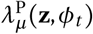 is the expected number of philopatric individuals (i.e., surviving adults or offspring that have remained in their natal group) and 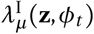 is the expected number of immigrant offspring (i.e., coming from other groups) with phenotype **z**. We aim to express these expected numbers in terms of the fitness of individuals at time *t*.

#### A.2.1 Individual fitness

The number, 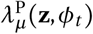, of philopatric individuals with phenotype **z** can be expressed in terms of fitness components of individuals at time *t* as

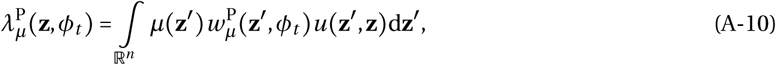

where 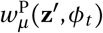 is philopatric fitness: it is the expected number of offspring produced by a *single* individual (including itself if it survives) bearing **z′** a time *t* (given that it resides in a state *µ* group); and *u* (**z′**, **z**) is the p.d.f. for the event that the offspring produced by an individual with phenotype **z′** has phenotype **z**. Note that we assume that surviving adults and offspring mutate alike. While this is relevant to unicellular organisms, an application specific to multicellular organisms would require distinguishing between two components of philopatric fitness: adult survival and offspring production. This would only complicate eq. (1) but would not affect our other results presented in the main text (eq. 3 onwards) as we later assume that mutations are rare, so that the chances of mutating during an individual’s lifetime would be unlikely (Appendix B.1.2).

Likewise, we can write the expected number of immigrant offspring as

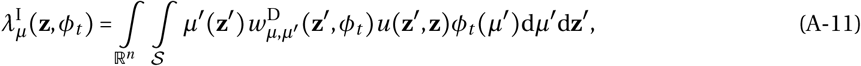

where 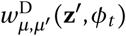 is the expected number of successful emigrant offspring of a *single* individual with phenotype **z′**, given that it resides in a group in state *µ′* ϵ 𝒮, and that the colonized group (i.e., the group the offspring lands in) was in state *µ* at time *t*.

Substituting eqs. (A-10) and (A-11) into eq. (A-9), which is in turn substituted into eq. (A-7), the phenotypic distribution at *t* +1 reads as

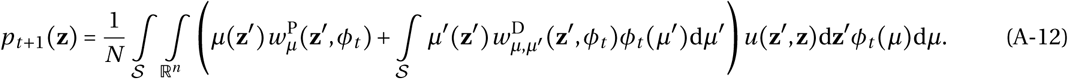

By exchanging integral variables *µ* and *µ′* in the second summand within brackets, we obtain

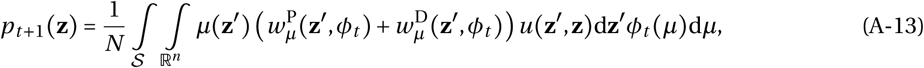

where

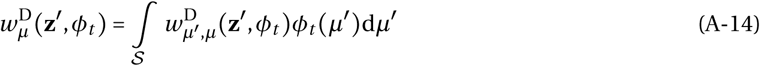

is the expected number of successful dispersing offspring produced by an individual with phenotype **z**′, given that this individual resides in a group in state *µ* at time *t*.

Individual fitness is then defined as

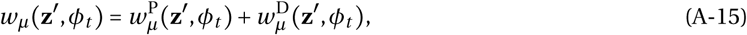

which gives the expected number of successful offspring produced by an individual with phenotype **z**′, given that this individual resides in a group in state *µ* at time *t* (and the population state distribution was *ϕ*_*t*_). In terms of this individual fitness function, the phenotypic distribution at time *t* +1 (eq. A-13) reads as

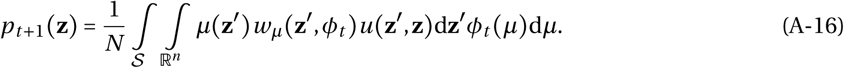

#### A.2.2 Lineage fitness

To go from eq. (A-16) to eq. (1) of the main text, let us define

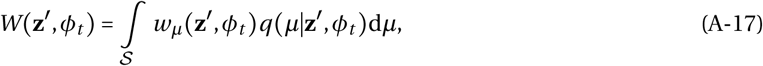

where

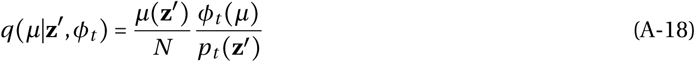

is the p.d.f. for the event that an individual resides in a group in state *µ* at time *t* given that this individual bears phenotype **z**′. In other words, *q* (*µ*|**z**′, *ϕ*_*t*_) gives the probability that an individual, randomly sampled at time *t* from the collection of individuals with phenotype **z**′ in the population (the “**z**′-lineage”), resides in a group in a state *µ*. As such, *W*(**z**′, *ϕ*_*t*_) (eq. A-17) is the expected fitness of a member of the **z**′-lineage at time *t* (where expectation is taken over all possible groups this member can belong to) and a multi-allelic version of *lineage fitness* (Mullon et al., 2016, Lehmann et al., 2016).

Substituting eq. (A-17) into eq. (A-16), we obtain that the individual phenotypic density distribution is

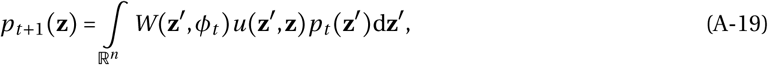

which combines the forces of mutation and selection on phenotypic change. To start disentangling these, note that when the probability of a mutation is independent from parental phenotype, the p.d.f. for the event that the offspring of an individual with phenotype **z**′ has phenotype **z** can be expressed as

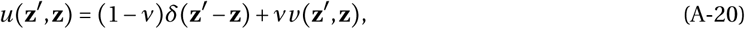

where *v* is the probability that an offspring has a mutant phenotype (i.e., 1– *u* (**z, z) =** *v* for all **z**), *δ* (**z**′ – **z**) is the Dirac delta function, and *v*(**z**′, **z**) is the conditional probability of mutating from **z**′ to **z** given that a mutation has occurred. So the first term of eq. (A-20) captures the event of no mutation, in which case the offspring has the same phenotype than its parent, and the second term captures the event of a mutation. Substituting eq. (A-20) into eq. (A-19), we finally obtain eq. (1) in the main text, as required.

## B The dynamics of trait means and variance-covariance

Here, we derive eqs. (3)-(7) of the main text, which govern the closed dynamics of trait means and variance-covariance. As mentioned in the main text, this derivation hinges upon several assumptions that we detail below.

### B.1 Weak selection and mutation

#### B.1.1 Weak selection

We first assume that the phenotypic distribution, *p*_*t*_(**z**), is peaked around the population mean 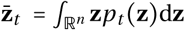 (i.e., the phenotypic variance is small). We can thus approximate lineage fitness, *W* (**z**, *ϕ*_*t*_), as a second-order Taylor expansion around 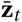 We do so in Appendix C, in which we show that lineage fitness can be written as

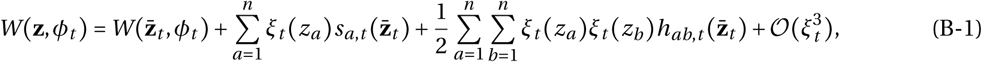

where

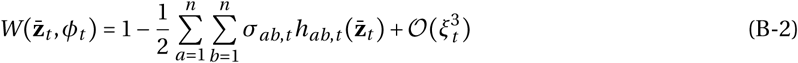

is the lineage fitness of the average phenotype 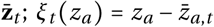 denotes the difference between a value *z*_*a*_ and the average trait value 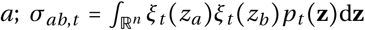 is the (co)variance among traits *a* and *b* in the population; 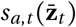 is the first-order effect of change in trait *a* away from 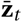 on lineage fitness (i.e., 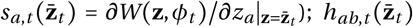 is the second-order effect of a joint change in traits *a* and *b* away from 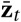 on lineage fitness (i.e., 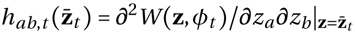); and *ξ*_*t*_ is the maximum deviation between individual trait value in the population and the population mean trait value at time *t*. We detail the first- and second-order effects below.

The first-order effect is given by

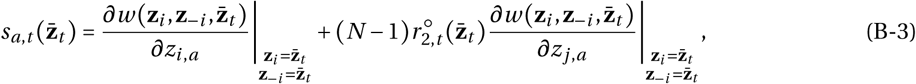

where individual fitness, 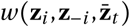, is written as in the main text eq. (5) and 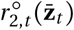 is a neutral time dependent coefficient of pairwise relatedness (i.e., the probability that two individuals sampled at random within a group at time *t* are identical-by-descent in the absence of selection, see section C.1.2 in Appendix C for more details).

The second-order effect is given by

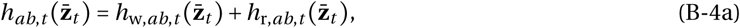

With

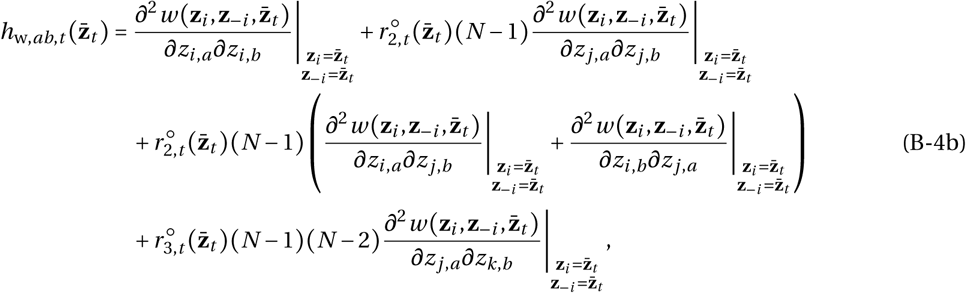

and

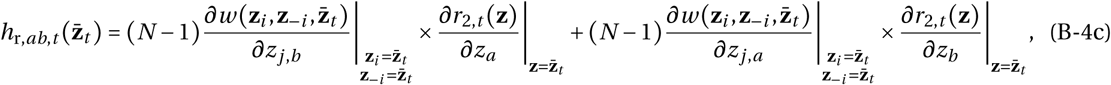

where 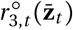 is the neutral time-dependent three-way relatedness (i.e., the probability that three individuals sampled at random within a group at time *t* are identical-by-descent in the absence of selection, see eq. C-44 for details); and 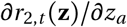 is the marginal effect of a change in trait *a* on time-dependent pairwise relatedness (i.e., the effect of trait *a* on the probability that a neighbor of a focal individual with phenotype **z** carries a phenotype that is identical-by-descent to that of the focal at time *t*, see section C.1.2 in Appendix C for more details).

The first (eq. B-3) and second (eq. B-4) order effects are the same as the selection gradient (eq. 6) and correlational selection (eq. 7) of the main text, respectively, with the exception that relatedness coefficients 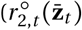, 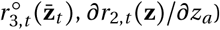 are time-dependent in eqs. (B-3) and (B-4) and independent in eqs. (6) and (7). We will specify in section B.2.2 below how we can get rid of this time dependence, but first, we need to make a further assumption.

#### B.1.2 Weak mutation

Our next assumption is that mutations are rare, with the probability of mutating, *v*, of the order 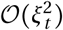. Under this assumption, note that 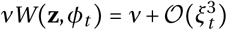 (from eqs. B-1-B-2). We can therefore rewrite eq. (1) of the main text as

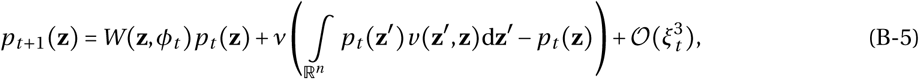

where the first term captures the effects of selection only, and the next term, the effects of mutation only. Eq. (B-5) takes the same functional form as classical recurrence for the phenotypic distribution in well-mixed populations when selection and mutation are weak (under the continuum-of-alleles model, e.g., eq. 1 of Bürger, 1986; for fluctuating population size, see eq. 4.9 of Champagnat et al., 2006), but with lineage, *W* (**z**, *ϕ*_*t*_), instead of individual fitness. Next, we use eq. (B-5) to derive recurrence equations for the changes in mean trait values and the phenotypic variance-covariance matrix over one time period.

#### B.1.3 Dynamics of the mean trait values

By definition, the change in the mean of trait *a* over one time period is

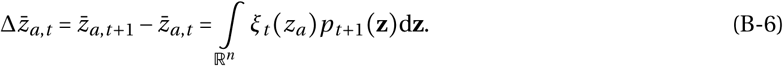

Substituting eq. (B-5) into eq. (B-6), we obtain

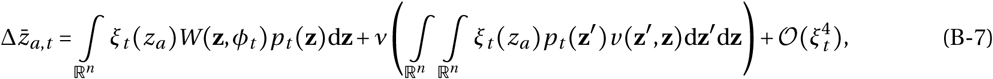

But since the effects of mutation are assumed to be unbiased, we have

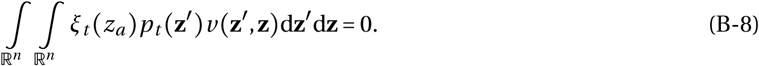

Eq. (B-7) then reduces to

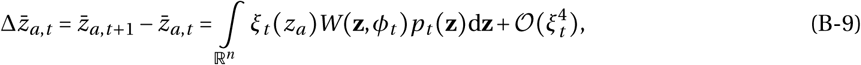

which corresponds to the first term of the Price equation: the change in average trait value in a population is equal to the covariance between trait and fitness (Price, 1970; see eq. 3 of Frank, 1997).

Substituting (eq. B-1) into eq. (B-9), we obtain that the change in the mean of trait *a* is,

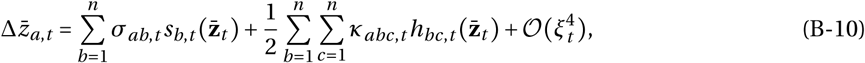

which depends on the skew,

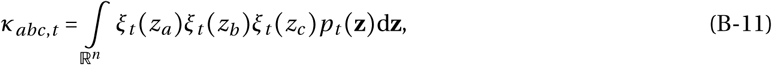

in the population at time period *t* (in line with e.g., eq. 8a of Wakano and Iwasa, 2013 and eq. A20 b of Débarre and Otto, 2016 in well-mixed populations; eq. 17 of Wakano and Lehmann, 2014 for the island model).

#### B.1.4 Dynamics of the phenotypic variance-covariance

By definition, the change in the (co)variance (within individuals) between two traits *a* and *b* over one time period is

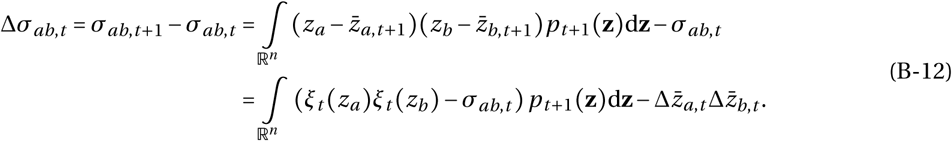

Substituting eq. (B-5) into the above, we obtain

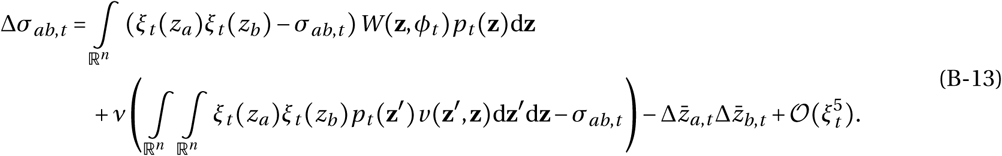

The bracketed term in the second line of eq. (B-13), which captures the effects of mutations, can be simplified by first writing out the product of deviations in terms of parental phenotype as

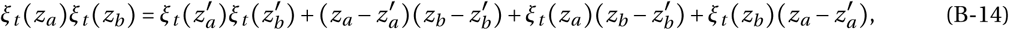

and second, by noting that since mutations are assumed to be unbiased, the covariance between parental phenotype and mutation effect is zero:

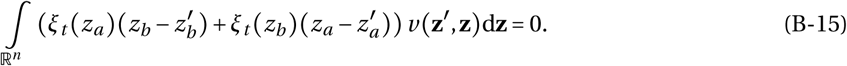

Using eqs. (B-14)-(B-15), the effect of mutations in eq. (B-13) can then be written as

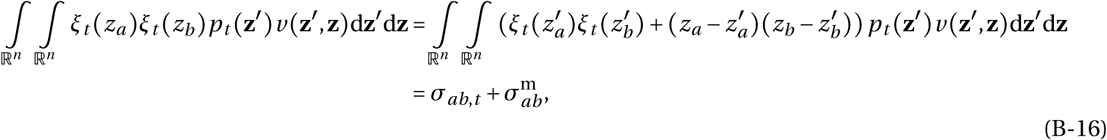

where we have defined, 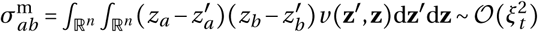, as the (co)variance in mutational effects on traits *a* and *b*. Substituting eq. (B-16) into eq. (B-13), we obtain that the change in the (co)variance between two traits *a* and *b* over one time period is

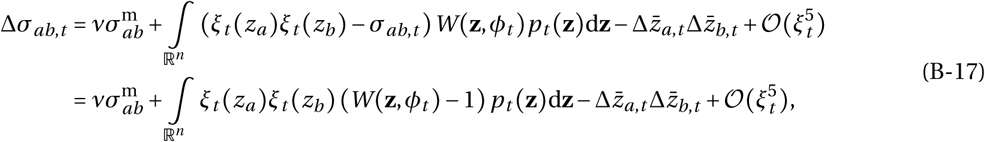

where to go from the first to the second line, we have used the fact that mean lineage fitness is one: 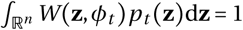 (since the population size is constant).

Substituting eq. (B-1), and the change in mean, eq. (B-10), into eq. (B-17), we obtain after some re-arrangements that the one-generational change in phenotypic (co)variance between traits *a* and *b* is

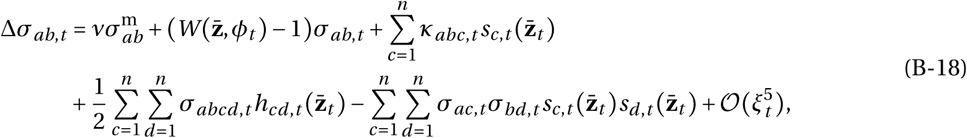

which depends on the fourth central moment of the phenotypic distribution,

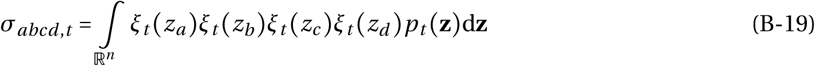

(in line with e.g., eq. 8b of Wakano and Iwasa, 2013 and eq. A24 b of Débarre and Otto, 2016 in well-mixed populations; eqs. B1-B8 of Wakano and Lehmann, 2014 for the island model with a single trait).

### B.2 Closure assumptions

Finally, we close the dynamical system for the means and (co)variances (given by eqs. B-10 & B-18). We achieve this closure in two steps.

#### B.2.1 Normal closure

First, we assume that the phenotypic distribution, *p*_*t*_(**z**), is normal. Under this assumption, the skew in the phenotypic distribution is zero, *κ* _*abc,t*_ = 0, and the fourth central moments can be expressed in terms of the (co)variances, *σ*_*abcd,t*_ *= σ*_*ab,t*_ *σ*_*cd,t*_+ *σ*_*ac,t*_ *σ*_*bd,t*_+ *σ*_*ad,t*_ *σ*_*bc,t*_. Substituting these relationships into eqs (B-10) and (B-18), we obtain that the one-generational changes in means and covariances are respectively given by

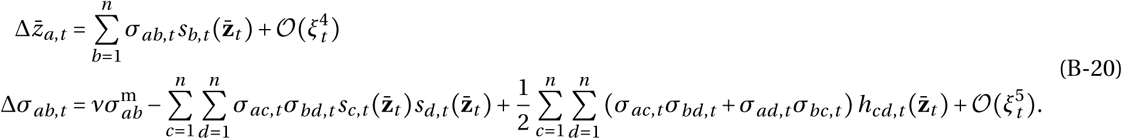

Since we make no assumption about the order of the fitness effects of traits (i.e., 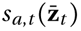 and 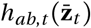 can be of order 𝒪(1)), the magnitude of a one-generational change in mean trait *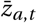* and (co)variance *σ*_*ab,t*_ are respectively of order 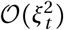 and 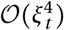. In vector and matrix form, eq. (B-20) corresponds to eq. (3) of the main text, except that in eq. (B-20), the selection coefficients depend on time *t* (due to time-dependent relatedness coefficients, 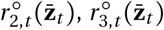, and 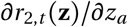. We get rid off of this dependency and finally achieve closure in the next section.

#### B.2.2 Quasi-equilibrium

Our second step to close the dynamical system eq. (B-20) is to assume that dispersal is strong enough (relative to selection) so that genetic associations between individuals within groups reach their steady-state values before any significant changes has occurred in the phenotypic distribution, *p*_*t*_ **z**, at the population level. This *quasi-equilibrium* assumption, which is frequently used in population genetic and social evolution theory (e.g., Kimura, 1965a, Nagylaki, 1993, Kirkpatrick et al., 2002, Roze and Rousset, 2005, 2008) is in line with our assumption that selection is weak. It entails that we can evaluate 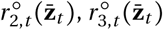, and 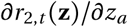 in eqs. (B-3)-(B-4) at their quasi-equilibrium, i.e., we take the limits 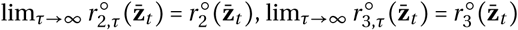, and 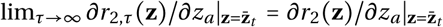, while holding *p*_*t*_(**z**) constant (we thus denote by 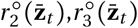, and 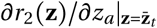, the steady-state values of neutral pairwise relatedness, neutral three-way relatedness, and the first-order perturbation of pairwise relatedness, respectively). Substituting these steady-states into the selection coefficients eqs. (B-3)-(B-4) (now independent of time so written as 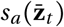 and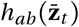, which are in turn substituted into eq. (B-20), we finally obtain the closed dynamical eqs. (3) of the main text.

##### Computing relatedness coefficients

Computing relatedness coefficients under neutrality (i.e., 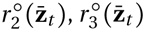) is standard in population genetics (e.g., Karlin, 1968, Rousset, 2004). When generations are non-overlapping (i.e., a Wright-Fisher life cycle), for example, the relevant relatedness coefficients for our approach are given by

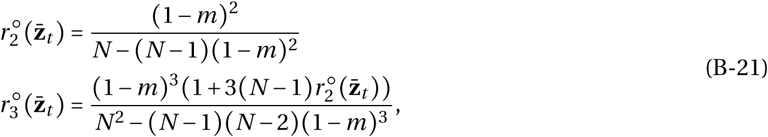

(e.g., eqs. 12a & 12b of Ohtsuki, 2010; see also Table 1 of Mullon et al., 2016 for the Moran model). Calculating the first-order effect of selection on pairwise relatedness, 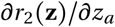, however, is more complicated. Under the quasi-equilibrium assumption, a perturbation of genetic associations between individuals will depend on first-order perturbations of individual fitness and neutral relatedness coefficients (see Roze and Rousset, 2008 for a general treatment, in particular their eq. 67). So far, the first-order effect of selection on pairwise relatedness, 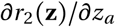, has been explicitly derived for two standard life-cycles, the semelparous Wright-Fisher life-cycle (in which all adults die after reproduction; see eq. 18 of Ajar, 2003 and eq. 28 of Wakano and Lehmann, 2014) and the iteroparous birth-death Moran life-cycle (in which a single adult dies after reproduction in each group; see eq. 14 of Mullon et al., 2016). In both cases, the effect of selection on relatedness can be written as

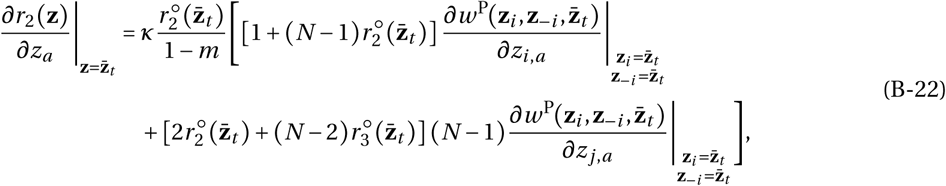

where 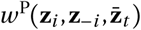 is the expected number of offspring of a focal individual (with phenotype **z**_*i*_, with neighbours **z**_−*i*_, and individuals in other groups with phenotype 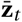) that successfully establish in their natal group; and *κ* = 2 for the Wright-fisher and *κ* = 1 for the Moran life cycle (watch out for an unfortunate typo in eq. 15 of Mullon et al., 2018, which has “*κ* = *N*” under the Moran life cycle).

## C Second-order approximation of lineage fitness

Here, we derive the second-order Taylor expansion of lineage fitness, *W*(**z**, *ϕ*_*t*_), around the population mean phenotype **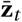** (i.e., we derive eqs. B-1 – B-4 of Appendix B). Let us first recall the definition of lineage fitness,

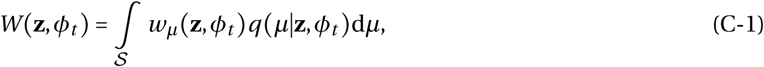

where *w*_*μ*_ (**z**, *ϕ*_*t*_) is the fitness of an individual with phenotype **z** in a group in state *µ*. Our approach is to develop a second-order Taylor expansion of the individual fitness function *w*_*μ*_ (**z**, *ϕ*_*t*_), which we then plug back into eq. (C-1) to average it over group composition, *q(µ*|**z**, *ϕ*_*t*_), and thus obtain lineage fitness.

Our starting point is to rewrite individual fitness as

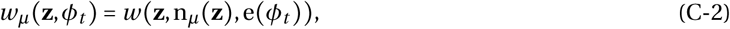

i.e., as a function that depends explicitly on all relevant phenotypes in the population: (1) **z** ∊ ℝ^*n*^, the phenotype of the focal individual (the individual whose fitness is under scrutiny); (2) n_*μ*_ (**z**) ∊ ℝ ^*(N−*1)×*n*^, the set of phenotypes of the *N −*1 neighbors of the focal individual; and (3) e(*ϕ*_*t*_) ∈ ℝ^*N×n*^, the set of *N* phenotypes from a representative (or average) group other than the one the focal resides in (which depends on the population state (*ϕ*_*t*_).

From these dependencies, the Taylor expansion of individual fitness, *w*(**z**, n_*µ*_ **(z**), e(*ϕ*_*t*_)), around **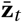** has the generic form,

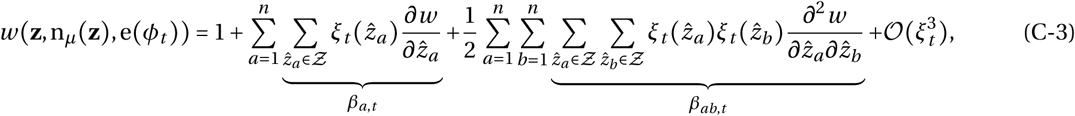

where 1 is individual fitness in a monomorphic population (i.e., when all individuals have phenotype **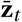**); **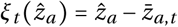** denotes the difference between a value **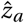** and the average trait value *a* (and *ξ*_*t*_ in **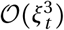** is the maximum deviation between individual trait value in the population and the population mean trait value at time *t*); and the set *Ƶ* = {**z**} ⋃ n_*µ*_ (**z**) ⋃ e (*ϕ*_*t*_) collects the phenotypes of the focal, its neighbors, and those in other groups (so that it has 2*N* elements). The term *β*_*a,t*_ in eq. (C-3) collects the marginal effects of a change in trait *a* on focal fitness: it sums the marginal effects of changing trait *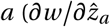*, where here and hereafter all derivatives are evaluated when all individuals have mean phenotype *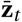*) across all individuals that belong to *Ƶ*. Similarly, *β*_*ab,t*_ collects the interaction effects of changes in traits *a* and *b* on focal fitness (summing the interaction effects of changes in traits *a* and *b* within, and between, all individuals that are in *Ƶ*).

We will develop these marginal (*β*_*a,t*_ in section C.1) and interaction (*β*_*ab,t*_ in section C.2) effects on individual fitness, and then average them over the group distribution an individual can reside in, *q*(*µ*|**z**, *ϕ*_*t*_), to obtain lineage fitness (eq. C-1). But first, note that lineage fitness in a monomorphic population is,

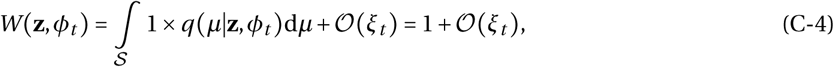

i.e., also one.

### C.1 Marginal effects

We first develop the marginal effects, *β*_*a,t*_, of varying trait *a* on individual fitness. To distinguish the effects of varying the trait in different individuals, we will use the symbols

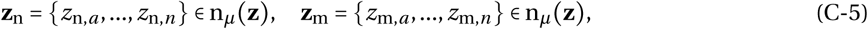

to denote the phenotypes of two distinct individuals from the focal group (and distinct from the focal individual), and

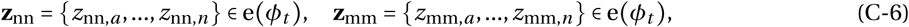

to denote the phenotypes of two distinct individuals from a group different to the focal.

With these notations, *β*_*a,t*_ can be decomposed into

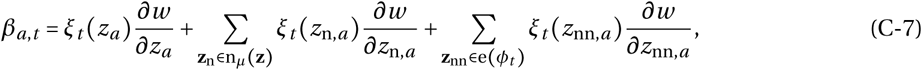

where the first, second and third summands capture the marginal effect of varying the trait in: the focal; its neighbors; and individual in other groups, respectively. Because the fitness function *w* (**z**, n_*μ*_ (**z**), e(*ϕ*_*t*_)) is invariant to permutations of elements within the sets n_*µ*_ **z** and e *ϕ*_*t*_ (i.e., it does not matter to individual fitness which precise neighbor or individual from another group expresses which phenotype), the derivatives in eq. (C-7) can be taken out of their sums,

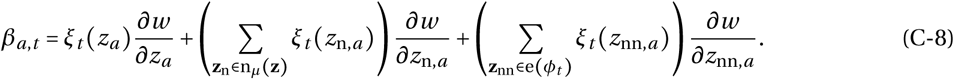

In order to make the sums in eq. (C-8) more convenient for averaging over *q*(*µ* | **z**, *ϕ*_*t*_), we seek to express them in terms of the counting measure *µ* **z** that counts the number of individuals with phenotype **z** in a group in state *µ* (see eq. A-2 in Appendix A). This is easily achieved for the last sum in eq. (C-8), which turns out to be zero:

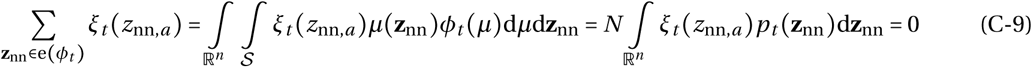

(this reflects that since there is an infinite number of groups, the average deviation from the mean in groups other than the focal is zero).

For the first sum between parenthesis in eq. (C-8), we define the conditional counting measure,

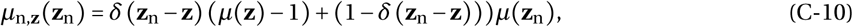

where *δ* (.) is the Dirac-Delta function, so that *µ*_n, **z**_ (**z**_n_) counts the number of neighbors of the focal that have phenotype **z**_n_, given that the focal individual has phenotype **z**. With eq. (C-10), we can write the first sum of eq. (C-8) as

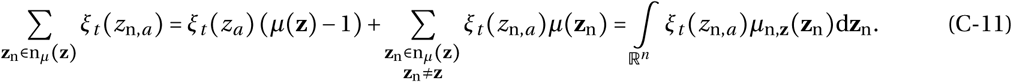

Substituting eqs. (C-9) & (C-11) into eq. (C-8) then gives

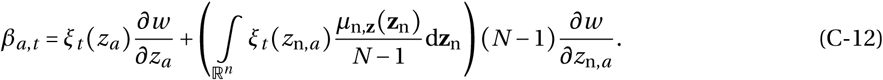

#### C.1.1 Average marginal effects

We proceed to average the marginal effects of a change in one trait, *β*_*a,t*_, over the group distribution an individual can reside in, *q* (*µ*|**z**, *ϕ*_*t*_), which is necessary to obtain lineage fitness (eq. C-1). From eq. (C-12), this average can be written as,

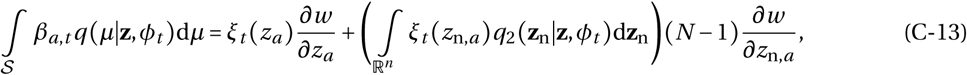

where we have defined

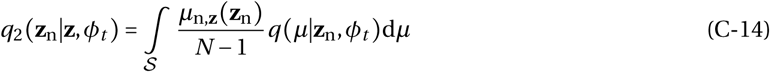

as the p.d.f. for the event of sampling an individual with phenotype **z**_n_ within a group, given that within this group a focal individual with phenotype **z** has already been sampled (and removed).

#### C.1.2 Pairwise relatedness

The p.d.f. *q*_2_ (**z**_n_|**z**, *ϕ*_*t*_) can be connected to the notion of pairwise relatedness by noting that two neighbors with the same phenotype may have a common ancestor who resided in the same group, which in the infinite island model is equivalent to the event that these individuals are identical-by-descent (IBD, e.g., Rousset, 2002). To make this connection explicit, we decompose *q*_2_ (**z**_n_ |**z**, *ϕ*_*t*_) as,

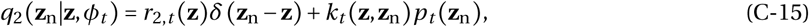

where *r*_2,*t*_ (**z**) is the conditional p.d.f for the event that, given a focal individual has phenotype **z** at time *t*, a randomly sampled individual among its neighbors is IBD to this focal (this p.d.f. depends on the whole phenotypic distribution, *ϕ*_*t*_, which is captured by the time index in *r*_2,*t*_ (**z**)). We refer to *r*_2,*t*_ (**z**) as pairwise relatedness. The two summands in eq. (C-15) respectively capture two complementary events: the sampled neighbor is either (1) IBD with the focal (and thus must have the same phenotype as the focal, **z**_n_ = **z**); or (2) not IBD with the focal and has phenotype **z**_n_ (which may or may not be equal to **z**). We have written the p.d.f. for this latter event as *k*_*t*_ (**z, z**_n_)*p*_*t*_ (**z**_n_), where *p*_*t*_ (**z**_n_) is the marginal probability of sampling an individual with phenotype **z**_n_ from the global population. The function *k*_*t*_ (**z, z**_n_) can therefore be viewed as the multiplicative effect of having already sampled an individual with phenotype **z** in the group on this marginal probability.

In our endeavour to obtain an expression for lineage fitness up to the order 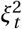, we seek to express *q* (**z**_n_ | **z**, *ϕ*_*t*_) to the order of *ξ*_*t*_ (as it multiplies *ξ*_*t*_ (*z*_*n,a*_), which is of order *ξ*_*t*_, in eq. C-13). We thus Taylor expand both *r*_2,*t*_ (**z**) and *k*_*t*_ (**z, z**_n_) in eq. (C-15) to the first-order around 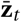, and obtain

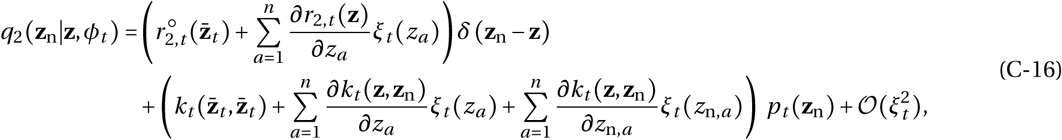

where 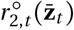 is the probability that two randomly sampled individuals within a group at time *t* are IBD under neutrality (i.e., when the population is monomorphic for 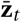).

Next, we use the fact that from the definition of *q*_2_ (**z**_n_ | **z**, *ϕ*_*t*_) (eq. C-14), *q*_2_ (**z**_n_ | **z**, *ϕ*_*t*_)*p*_*t*_ (**z**) is the (unconditional) p.d.f. for the event of sampling one individual with phenotype **z** and another with **z**_n_ *without* replacement from a group. Thus, three identities must hold:

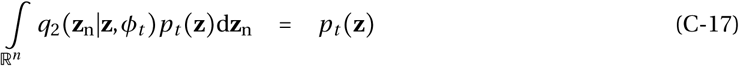

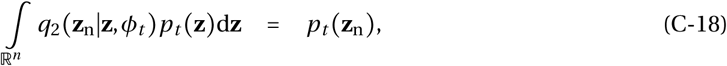

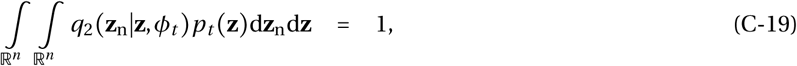

which can be used to express *k*_*t*_ (**z**, **z**_n_) in terms of *r*_2,*t*_ **(z**). In fact, substituting eq. (C-16) into eq. (C-19), we obtain

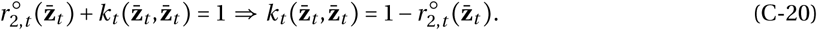

Substituting eq. (C-20) into eq. (C-16), which is in turn substituted into eq. (C-17) yields,

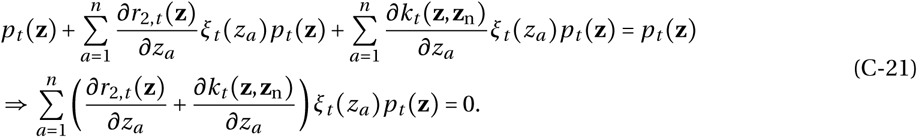

As the above equality holds for all possible *ξ*_*t*_ (*z*_*a*_) and for all *a* = 1, …, *n*, we must have

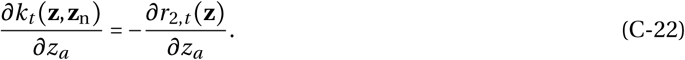

Substituting eq. (C-20) into eq. (C-16), which is in turn substituted into (C-18) and using a similar argument, we obtain that,

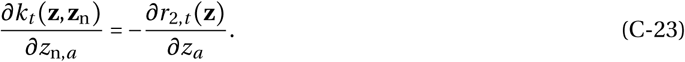

Plugging eqs. (C-20), (C-22) and (C-23) into eq. (C-16) then gives us

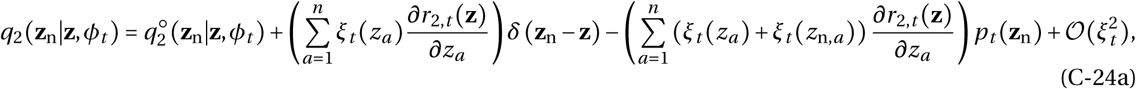

Where

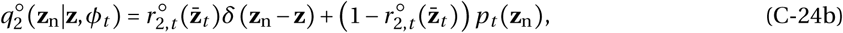

is the neutral (conditional) p.d.f. for the event of sampling an individual with phenotype **z**_n_ within the neigh-borhood of an individual with phenotype **z**.

##### Regression definition of relatedness

It is noteworthy that the definition of pairwise relatedness as the probability of IBD between two randomly sampled individuals within a group that we use, aligns with the “regression definition of relatedness” (e.g., Grafen, 1985, eq. 2.13 of Frank, 1998). Under this latter definition, relatedness is the regression of neighbor phenotype on focal phenotype: it is the ratio of phenotypic covariance among neighbors,

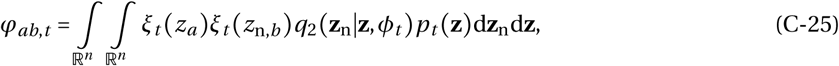

to the covariance within individuals, 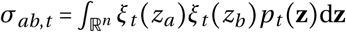. Substituting eq. (C-24) into eq. (C-25) to compute this ratio, we obtain

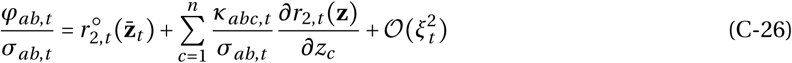

where 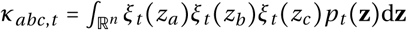 is the skew between traits *a, b* and *c* within individuals. Our eq. (C-26) is thus in line with the result that in the infinite island model and under neutrality, relatedness as a regression coefficient is equal to the probability of identity-by-descent (i.e., 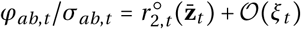, Rousset, 2002; for equivalent expressions to eq. C-26, see for e.g. eq. 7 of Queller, 1992a; eq. 2.13 of Frank, 1998; eq. 11 of Wakano and Lehmann, 2014).

##### Average marginal effects

We can now return to our calculation of the average marginal effects of a change in one trait (eq. C-13). Substituting eq. (C-24) into eq. (C-13), we in fact obtain

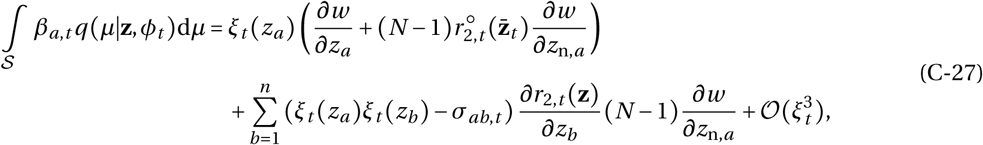

for the average marginal effect of trait *a*.

### C.2 Interaction effects

We now tackle the interaction effects on focal fitness, *β*_*ab,t*_ (eq. C-3), which we will then average over *q*(*μ*|**z**, *ϕ*_*t*_). Using notation eqs. (C-5)-(C-6), we first decompose *β*_*ab,t*_ as

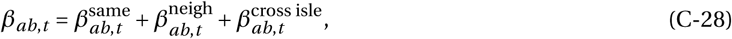

Where

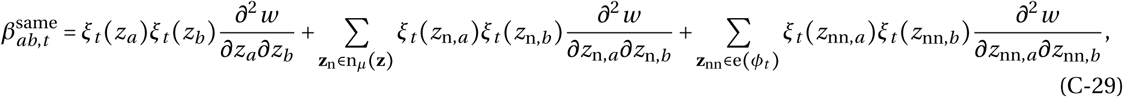

collects the interaction effects of traits *a* and *b* within individuals,

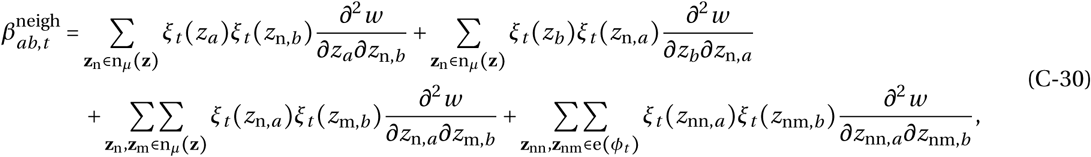

collects the interaction effects between neighbors, and

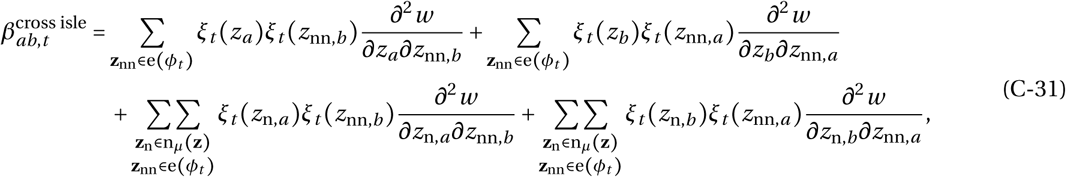

the interaction effects between individuals between groups. We proceed to specify each of these in terms of counting measures and average them over *q*(*μ*|**z**, *ϕ*_*t*_) sequentially.

#### C.2.1 Interaction effects within individuals

We start with interaction effects within individuals, 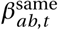. Using eq. (C-10), eq. (C-29) can be expressed as

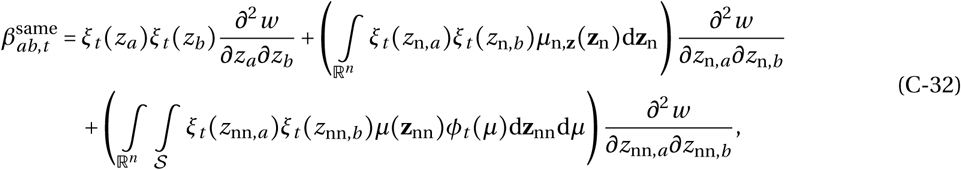

where the first summand consists of the interaction effect within the focal individual, the second within neighbors of the focal, and the third within individuals from other groups than the focal. To simplify these, we use the fact that because the population size remains constant, we have

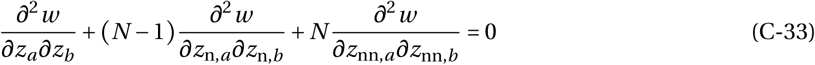

(see eq. B.13 of Wakano and Lehmann, 2014); and that the third term of eq. (C-32) can be expressed as

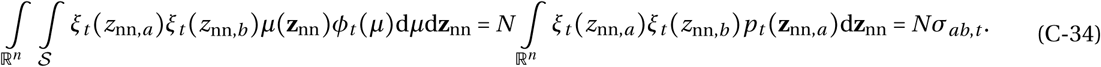

Substituting eqs. (C-33) and (C-34) into eq. (C-32), we find that the interaction effects of *a* and *b* within individuals can be written as

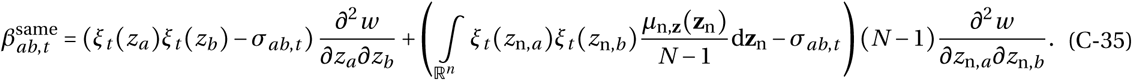

Averaging these interaction effects within individuals over *q* (*μ*|**z**, *ϕ*_*t*_) then gives

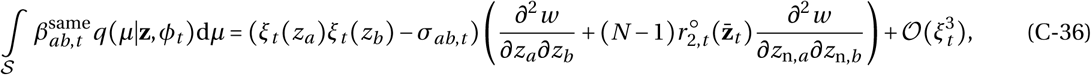

where we used definition eq. (C-14) and eq. (C-24).

#### C.2.2 Interaction effects between neighbors

Let us now turn to interaction effects between neighbors, 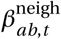 (eq. C-30). It is composed of four summands: the first two capture the interaction effects between the focal and its neighbors; the third, between two neighbors of the focal; and the fourth, between two neighbors from another group than the focal. We consider these separately below.

##### Interaction effects between the focal and its neighbors

The first summand of 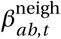 (eq. C-30) corresponds to the interaction effect between trait *a* in the focal and trait *b* in its neighbors. Using eq. (C-10), it can be expressed as

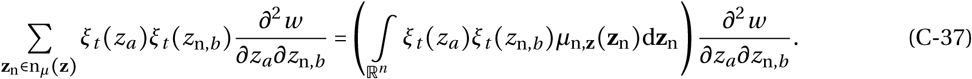

Averaging eq. (C-37) over *q* (*μ*|**z**, *ϕ*_*t*_) then reads as

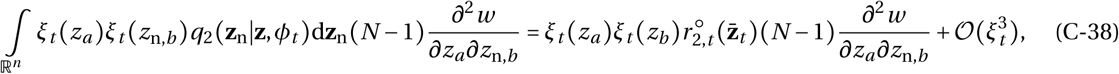

where we used eq. (C-24). Similarly, averaging the second summand of 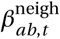 (eq. C-30), which corresponds to the interaction effect between trait *b* in the focal and trait *a* in its neighbors, yields

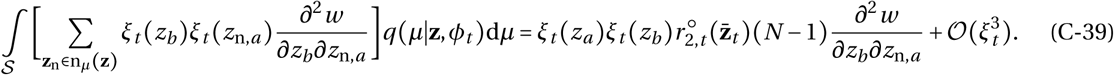

##### Interaction effects between neighbors of the focal

The third summand of 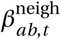 (eq. C-30) collects the interaction effects between neighbors of the focal. To express these in terms of the counting measure *μ* **z** and average them over *q*(*µ*|**z**, *ϕ*_*t*_), we introduce one further conditional counting measure,

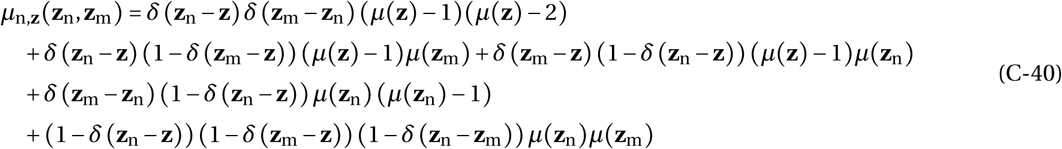

which counts the number of unordered pairs of neighbors that have phenotypes **z**_n_ and **z**_m_, given that the focal individual has phenotype **z**. Integrated over ℝ^*n*^, the first line of eq. (C-40) counts the number of pairs of neighbors that have focal phenotype **z**; the second line, the number of pairs of neighbors in which only one has the focal phenotype **z**; the third line, the number of pairs in which both neighbors have the same phenotype that is different to the the focal; and the fourth line, the number of pairs in which neighbors have phenotypes that are different to one another, and to the focal.

Using eq. (C-40), we can then re-write the third summand of 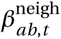 as

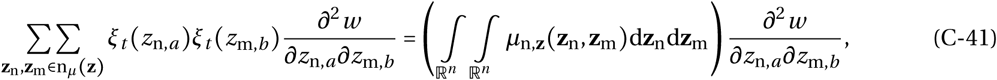

which averaged over *q*(*µ*|**z**, *ϕ*_*t*_) reads as

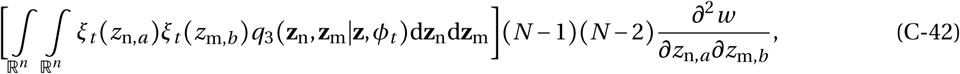

where

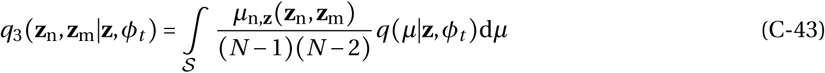

is the conditional p.d.f. for the event of sampling two individuals without replacement, one with phenotype **z**_n_ and another with phenotype **z**_m_ in a group, given that a focal individual with phenotype **z** has already been sampled in that group at time *t*. This p.d.f. can be connected to notions of relatedness as individuals that have the same phenotype may be IBD. In fact, using a coalescence argument, we can rewrite *q*_3_ (**z**_n_, **z**_m_|**z**, *ϕ*_*t*_) as

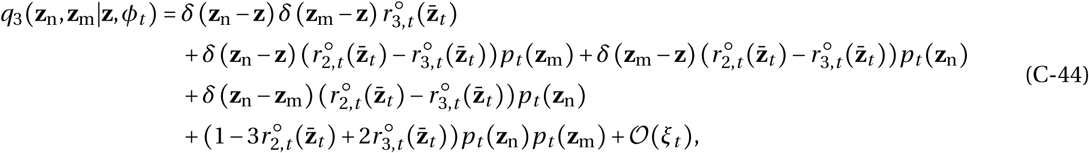

where 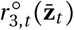 is the probability of sampling three individuals without replacement from a group are IBD in the absence of selection at time *t* (i.e., three-way relatedness). Each summand of eq. (C-44) capture a different possible relationship among the sampled (**z**_n_ and **z**_m_) and focal (**z**) phenotype. The first line of eq. (C-44) captures the event of sampling two individuals that are both IBD with the focal (and thus with the same phenotype as the focal, **z = z**_n_ **= z**_m_); the next line, the event of sampling only one individual IBD with the focal (and thus with the same phenotype: the two summands respectively capture the cases **z = z**_n_ and **z = z**_m_); the third line, the event of sampling two individuals that are IBD together (and thus with phenotypes **z**_n_ = **z**_m_) but not with the focal; and the last line, the event of sampling two individuals that are not IBD with one another or with the focal.

Substituting eq. (C-44) into eq. (C-42) then gives us

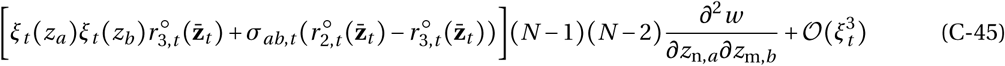

for the average interaction effects between neighbors of the focal.

##### Interaction effects between neighbors from other groups

The fourth and final summand of 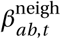 (eq. C-30), which collects the interaction effects between neighbors from other groups, can be expressed using eq. (C-10) as

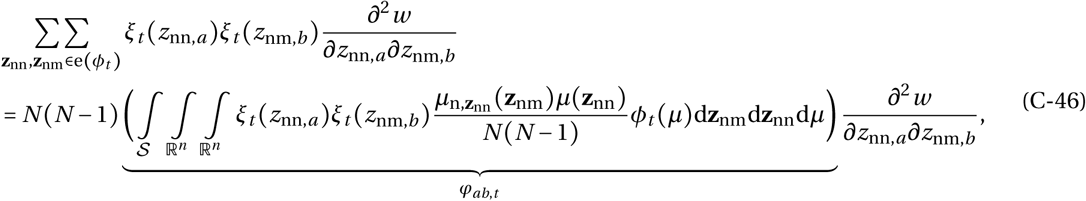

where *ϕ*_*ab,t*_ is the covariance among traits *a* and *b* between individuals within groups (see eq. C-25). We can then use eq. (C-26) to specify this covariance and obtain,

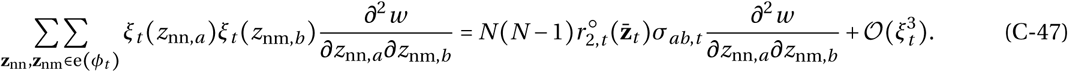

Since this expression does not depend on group composition *µ*, it will be invariant to averaging over *q*(*µ*|**z**, *ϕ*_*t*_).

##### Average interaction effects between neighbors

Hence, the average interaction effects between neighbors 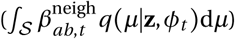, which is given by the sum of eqs. (C-38), (C-39), (C-45) and (C-47), reads as

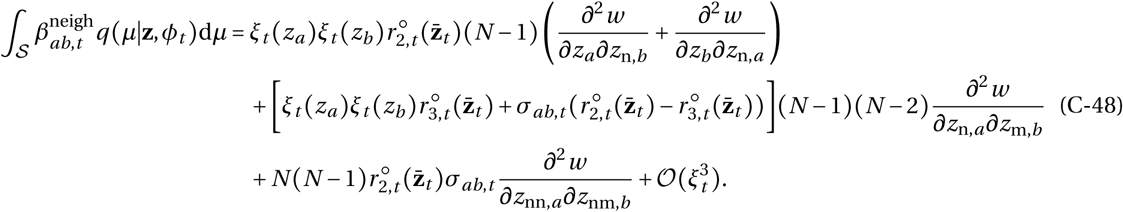

This can be further simplified by using the fact than since the total population size is constant, the following holds,

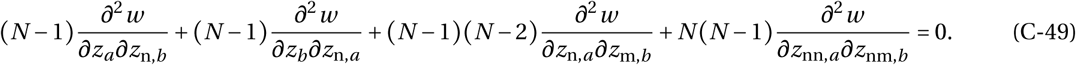

(see eq. B.14 of Wakano and Lehmann, 2014). Substituting for 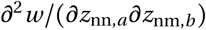 using eq. (C-49) into eq. (C-48) finally gives

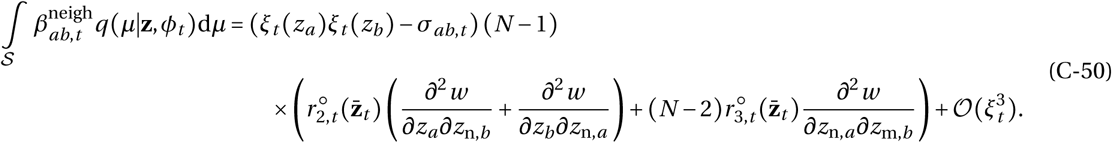

#### C.2.3 Interaction effects between individuals between groups

The final relevant interaction effect, 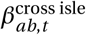 (eq. C-31), is the interaction between individuals that belong to different groups. However, it is straightforward to show that these effects vanish in the infinite island model of dispersal. For example, consider the first summand of eq. (C-31), which measures the effect of a change in trait *a* of the focal and a trait *b* in an individual from a group other than the focal:

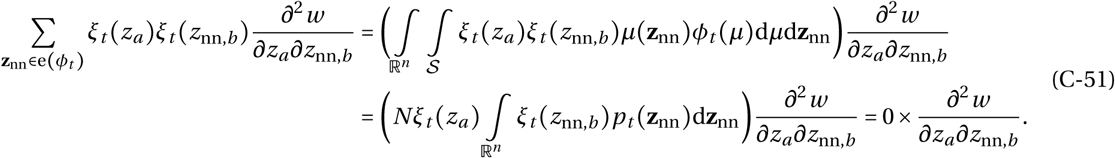

Similar arguments show that all the other summands of 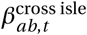 are also zero.

### C.3 Putting it all together

Summing eqs. (C-4), (C-27), (C-36), (C-50), we finally obtain the second order expansion of lineage fitness,

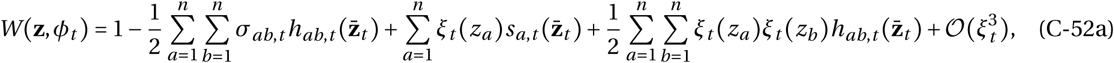

where

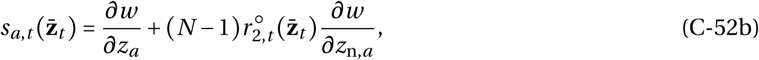

and

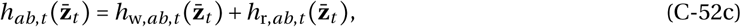

with

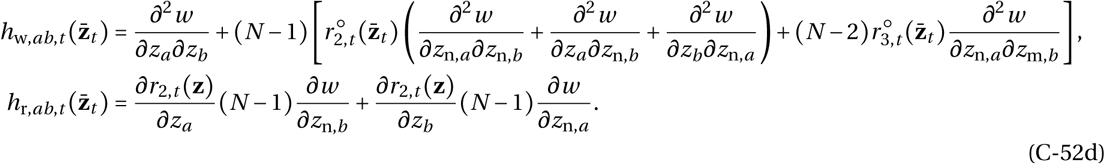

Eq. (C-52) is equivalent to eq. B-1 – B-4 of Appendix B, in which we write the derivatives of individual fitness *w*(**z**, n_*µ*_(**z**), e(*ϕ*_*t*_)) with respect to *z*_*a*_, *z*_n,*a*_, and *z*_nn,*a*_, in terms of the derivatives of the individual fitness function 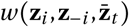 (eq. 5 of main text) with respect to *z*_*i,a*_, *z*_*j,a*_, and *z*_*k,a*_, respectively; and add evaluation signs to all derivatives at the population mean 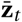.

## D Individual-based simulations

We performed individual based simulations for a population composed of *N*_d_ groups, each populated by *N* individuals, using Mathematica 11.0.1.0 (Wolfram Research, 2016). Starting with a monomorphic population, we track the evolution of the multidimensional phenotypic distribution under the constant influx of mutations. Each individual *i* ∈ {1, …, *N*_d_*N*} is characterized by two traits (*z*_*i*,1_, *z*_*i*,2_). At the beginning of a generation, we calculate the fecundity *f*_*i*_ of each individual according to its traits and those of its neighbors (using eq. 14). Then, we form the next generation of adults by sampling *N* individuals in each group with replacement according to parental fecundity, but to capture limited dispersal, the fecundity of each individual from the parental generation is weighted according to whether or not they belong to the group on which the breeding spot is filled: if an individual belongs to the same group in which a breeding spot is filled, its weighted fecundity is *f*_*i*_(1 − *m*), where *m* is the dispersal probability; if it belongs to another group, its weighted fecundity is *f*_*i*_ *m/*(*N*_d_ − 1) (as a disperser is equally likely to reach any other group, it lands with probability 1/(*N*_d_ − 1) in a focal group). Once an individual is chosen to fill the breeding spot, it mutates with probability *v*, in which case we add to parental values a perturbation that is sampled from a multivariate normal distribution with mean (0,0) and variance-covariance matrix 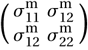. The resulting phenotypic values are truncated to remain between 0 and 4. We repeat the procedure for a fixed number of generations (see Figures for parameter values).

## Footnotes

1 When the size of the population fluctuates, 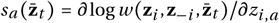, due to normalization of focal fitness with respect to mean fitness (see section 3.1; and eq. A6 of Iwasa et al., 1991 for how this holds when selection is frequency-dependent). If the size of the population fluctuates but selection is frequency-independent, then the selection gradient can be expressed as the derivative of the log of mean fitness in the population with respect to the trait under scrutiny (e.g., eq. 7b of Lande, 1979)

