## Supplementary Information for "An evolutionary quantitative genetics model for phenotypic (co)variance under limited dispersal, with an application to socially synergistic traits" (from another version of this preprint: version 2, 2019-03-02)

### Online Supplementary Information for *A quantitative genetics approach to the evolution of phenotypic (co)variance under limited dispersal, with an application to socially synergistic traits*

Charles Mullon*[1] and Laurent Lehmann[1]

[1] *Department of Ecology and Evolution, University of Lausanne, 1015 Lausanne, Switzerland*

In this supplementary information, we derive the second-order Taylor expansion of lineage fitness, $W(\mathbf{z},\phi_t)$, around the population mean phenotype $\bar{\mathbf{z}}_t$ (i.e., we derive eqs. B-1 – B-4 of Appendix B). Let us first recall the definition of lineage fitness,

$$W(\mathbf{z},\phi_t) = \int_{\mathcal{S}} w_\mu(\mathbf{z},\phi_t) q_t(\mu|\mathbf{z}) \mathrm{d}\mu, \tag{SI-1}$$

where $w_\mu(\mathbf{z},\phi_t)$ is the fitness of an individual with phenotype $\mathbf{z}$ in a group in state $\mu$. Our approach is to develop a second-order Taylor expansion of the individual fitness function $w_\mu(\mathbf{z},\phi_t)$, which we then plug back into eq. (SI-1) to average it over group composition, $q_t(\mu|\mathbf{z})$, and thus obtain lineage fitness.

Our starting point is to rewrite individual fitness as

$$w_\mu(\mathbf{z},\phi_t) = w(\mathbf{z}, \mathrm{n}_\mu(\mathbf{z}), \mathrm{e}(\phi_t)), \tag{SI-2}$$

i.e., as a function that depends explicitly on all relevant phenotypes in the population: (1) $\mathbf{z} \in \mathbb{R}^n$, the phenotype of the focal individual (the individual whose fitness is under scrutiny); (2) $\mathrm{n}_\mu(\mathbf{z}) \in \mathbb{R}^{(N-1)\times n}$, the set of phenotypes of the $N-1$ neighbours of the focal individual; and (3) $\mathrm{e}(\phi_t) \in \mathbb{R}^{N\times n}$, the set of $N$ phenotypes from a representative (or average) group other than the one the focal resides in (which depends on the population state $\phi_t$).

From these dependencies, the Taylor expansion of individual fitness, $w(\mathbf{z}, \mathrm{n}_\mu(\mathbf{z}), \mathrm{e}(\phi_t))$, around $\bar{\mathbf{z}}_t$ has the generic form,

$$w(\mathbf{z}, \mathrm{n}_\mu(\mathbf{z}), \mathrm{e}(\phi_t)) = 1 + \sum_{a=1}^{n} \underbrace{\sum_{\hat{z}_a \in \mathcal{Z}} \xi_t(\hat{z}_a) \frac{\partial w}{\partial \hat{z}_a}}_{\beta_{a,t}} + \frac{1}{2} \sum_{a=1}^{n} \sum_{b=1}^{n} \underbrace{\sum_{\hat{z}_a \in \mathcal{Z}} \sum_{\hat{z}_b \in \mathcal{Z}} \xi_t(\hat{z}_a) \xi_t(\hat{z}_b) \frac{\partial^2 w}{\partial \hat{z}_a \partial \hat{z}_b}}_{\beta_{ab,t}} + \mathcal{O}(\xi_t^3), \tag{SI-3}$$

where 1 is individual fitness in a monomorphic population (i.e., when all individuals have phenotype $\bar{\mathbf{z}}_t$); $\xi_t(\hat{z}_a) = \hat{z}_a - \bar{z}_{a,t}$ denotes the difference between a value $\hat{z}_a$ and the average trait value $a$ (and $\xi_t$ in $\mathcal{O}(\xi_t^3)$ is the maximum deviation between individual trait value in the population and the population mean trait value at time $t$); and the set $\mathcal{Z} = \{\mathbf{z}\} \cup \mathrm{n}_\mu(\mathbf{z}) \cup \mathrm{e}(\phi_t)$ collects the phenotypes of the focal, its neighbours, and those in other groups (so that it has $2N$ elements). The term $\beta_{a,t}$ in eq. (SI-3) collects the marginal effects of a change in trait $a$ on focal fitness: it sums the marginal effects of changing trait $a$ ($\partial w/\partial \hat{z}_a$, where here and hereafter all derivatives are evaluated when all individuals have mean phenotype $\bar{\mathbf{z}}_t$) across all individuals that belong to $\mathcal{Z}$. Similarly, $\beta_{ab,t}$ collects the interaction effects of changes in traits $a$ and $b$ on focal fitness (summing the interaction effects of changes in traits $a$ and $b$ within, and between, all individuals that are in $\mathcal{Z}$).

*Electronic address: ``

We will develop these marginal ($\beta_{a,t}$) and interaction ($\beta_{ab,t}$) effects on individual fitness, and then average them over the group distribution an individual can reside in, $q_t(\mu|\mathbf{z})$. But first, note that lineage fitness in a monomorphic population is,

$$W(\mathbf{z},\phi_t) = \int_{\mathcal{S}} 1 \times q_t(\mu|\mathbf{z})\mathrm{d}\mu + \mathcal{O}(\xi_t) = 1 + \mathcal{O}(\xi_t), \tag{SI-4}$$

i.e., also one.

### 1 Marginal effects

We here develop the marginal effects, $\beta_{a,t}$, of varying trait $a$ on individual fitness and average them over $q_t(\mu|\mathbf{z})$. To distinguish the effects of varying the trait in different individuals, we will use the symbols

$$\mathbf{z}_{\mathrm{n}} = \{z_{\mathrm{n},a},...,z_{\mathrm{n},n}\} \in \mathrm{n}_\mu(\mathbf{z}), \quad \mathbf{z}_{\mathrm{m}} = \{z_{\mathrm{m},a},...,z_{\mathrm{m},n}\} \in \mathrm{n}_\mu(\mathbf{z}), \tag{SI-5}$$

to denote the phenotypes of two distinct individuals from the focal group (and distinct from the focal individual), and

$$\mathbf{z}_{\mathrm{nn}} = \{z_{\mathrm{nn},a},...,z_{\mathrm{nn},n}\} \in \mathrm{e}(\phi_t), \quad \mathbf{z}_{\mathrm{mm}} = \{z_{\mathrm{mm},a},...,z_{\mathrm{mm},n}\} \in \mathrm{e}(\phi_t), \tag{SI-6}$$

to denote the phenotypes of two distinct individuals from a group different to the focal.

With these notations, $\beta_{a,t}$ can be decomposed into

$$\beta_{a,t} = \xi_t(z_a)\frac{\partial w}{\partial z_a} + \sum_{\mathbf{z}_{\mathrm{n}}\in\mathrm{n}_\mu(\mathbf{z})} \xi_t(z_{\mathrm{n},a})\frac{\partial w}{\partial z_{\mathrm{n},a}} + \sum_{\mathbf{z}_{\mathrm{nn}}\in\mathrm{e}(\phi_t)} \xi_t(z_{\mathrm{nn},a})\frac{\partial w}{\partial z_{\mathrm{nn},a}}, \tag{SI-7}$$

where the first, second and third summands capture the marginal effect of varying the trait in: the focal; its neighbours; and individual in other groups, respectively. Because the fitness function $w(\mathbf{z},\mathrm{n}_\mu(\mathbf{z}),\mathrm{e}(\phi_t))$ is invariant to permutations of elements within the sets $\mathrm{n}_\mu(\mathbf{z})$ and $\mathrm{e}(\phi_t)$ (i.e., it does not matter to individual fitness which precise neighbour or individual from another group expresses which phenotype), the derivatives in eq. (SI-7) can be taken out of their sums,

$$\beta_{a,t} = \xi_t(z_a)\frac{\partial w}{\partial z_a} + \left(\sum_{\mathbf{z}_{\mathrm{n}}\in\mathrm{n}_\mu(\mathbf{z})} \xi_t(z_{\mathrm{n},a})\right)\frac{\partial w}{\partial z_{\mathrm{n},a}} + \left(\sum_{\mathbf{z}_{\mathrm{nn}}\in\mathrm{e}(\phi_t)} \xi_t(z_{\mathrm{nn},a})\right)\frac{\partial w}{\partial z_{\mathrm{nn},a}}. \tag{SI-8}$$

In order to make the sums in eq. (SI-8) more convenient for averaging over $q_t(\mu|\mathbf{z})$, we seek to express them in terms of the counting measure $\mu(\mathbf{z})$ that counts the number of individuals with phenotype $\mathbf{z}$ in a group in state $\mu$ (see eq. A-2 in Appendix A). This is easily achieved for the second sum in eq. (SI-8), which turns out to be zero:

$$\sum_{\mathbf{z}_{\mathrm{nn}}\in\mathrm{e}(\phi_t)} \xi_t(z_{\mathrm{nn},a}) = \int_{\mathbb{R}^n}\int_{\mathcal{S}} \xi_t(z_{\mathrm{nn},a})\mu(\mathbf{z}_{\mathrm{nn}})\phi_t(\mu)\mathrm{d}\mu\mathrm{d}\mathbf{z}_{\mathrm{nn}} = N\int_{\mathbb{R}^n} \xi_t(z_{\mathrm{nn},a})p_t(\mathbf{z}_{\mathrm{nn}})\mathrm{d}\mathbf{z}_{\mathrm{nn}} = 0 \tag{SI-9}$$

(this reflects that since there is an infinite number of groups, the average deviation from the mean in groups other than the focal is zero).

For the first sum between parenthesis in eq. (SI-8), we define the conditional counting measure,

$$\mu_{\mathrm{n},\mathbf{z}}(\mathbf{z}_{\mathrm{n}}) = \delta(\mathbf{z}_{\mathrm{n}} - \mathbf{z})(\mu(\mathbf{z}) - 1) + (1 - \delta(\mathbf{z}_{\mathrm{n}} - \mathbf{z}))\mu(\mathbf{z}_{\mathrm{n}}), \tag{SI-10}$$

where $\delta(.)$ is the Dirac-Delta function, so that $\mu_{\mathrm{n},\mathbf{z}}(\mathbf{z}_{\mathrm{n}})$ counts the number of neighbours of the focal that have phenotype $\mathbf{z}_{\mathrm{n}}$, given that the focal individual has phenotype $\mathbf{z}$. With eq. (SI-10), we can write the first sum of eq. (SI-8) as

$$\sum_{\mathbf{z}_{\mathrm{n}}\in\mathrm{n}_\mu(\mathbf{z})} \xi_t(z_{\mathrm{n},a}) = \xi_t(z_a)(\mu(\mathbf{z}) - 1) + \sum_{\substack{\mathbf{z}_{\mathrm{n}}\in\mathrm{n}_\mu(\mathbf{z}) \\ \mathbf{z}_{\mathrm{n}}\neq\mathbf{z}}} \xi_t(z_{\mathrm{n},a})\mu(\mathbf{z}_{\mathrm{n}}) = \int_{\mathbb{R}^n} \xi_t(z_{\mathrm{n},a})\mu_{\mathrm{n},\mathbf{z}}(\mathbf{z}_{\mathrm{n}})\mathrm{d}\mathbf{z}_{\mathrm{n}}. \tag{SI-11}$$

Substituting eqs. (SI-9) & (SI-11) into eq. (SI-8) then gives

$$\beta_{a,t} = \xi_t(z_a)\frac{\partial w}{\partial z_a} + \left(\int_{\mathbb{R}^n} \xi_t(z_{\mathrm{n},a})\frac{\mu_{\mathrm{n},\mathbf{z}}(\mathbf{z}_\mathrm{n})}{N-1}\mathrm{d}\mathbf{z}_\mathrm{n}\right)(N-1)\frac{\partial w}{\partial z_{\mathrm{n},a}}. \tag{SI-12}$$

#### 1.1 Average marginal effects

We proceed to average the marginal effects of a change in one trait, $\beta_{a,t}$, over the group distribution an individual can reside in, $q_t(\mu|\mathbf{z})$, which is necessary to obtain lineage fitness (eq. SI-1). From eq. (SI-12), this average can be written as,

$$\int_{\mathcal{S}} \beta_{a,t} q_t(\mu|\mathbf{z})\mathrm{d}\mu = \xi_t(z_a)\frac{\partial w}{\partial z_a} + \left(\int_{\mathbb{R}^n} \xi_t(z_{\mathrm{n},a}) q_{2,t}(\mathbf{z}_\mathrm{n}|\mathbf{z})\mathrm{d}\mathbf{z}_\mathrm{n}\right)(N-1)\frac{\partial w}{\partial z_{\mathrm{n},a}}, \tag{SI-13}$$

where we have defined

$$q_{2,t}(\mathbf{z}_\mathrm{n}|\mathbf{z}) = \int_{\mathcal{S}} \frac{\mu_{\mathrm{n},\mathbf{z}}(\mathbf{z}_\mathrm{n})}{N-1} q_t(\mu|\mathbf{z}_\mathrm{n})\mathrm{d}\mu \tag{SI-14}$$

as the p.d.f. for the event of sampling an individual with phenotype $\mathbf{z}_\mathrm{n}$ within a group, given that within this group a focal individual with phenotype $\mathbf{z}$ has already been sampled (and removed).

##### 1.1.1 Pairwise relatedness

The p.d.f. $q_{2,t}(\mathbf{z}_\mathrm{n}|\mathbf{z})$ can be connected to the notion of pairwise relatedness by noting that two neighbours with the same phenotype may have a common ancestor who resided in the same group, which in the infinite island model is equivalent to the event that these individuals are identical-by-descent (IBD, e.g., Rousset, 2002). To make this connection explicit, we decompose $q_{2,t}(\mathbf{z}_\mathrm{n}|\mathbf{z})$ as,

$$q_{2,t}(\mathbf{z}_\mathrm{n}|\mathbf{z}) = r_{2,t}(\mathbf{z})\delta(\mathbf{z}_\mathrm{n} - \mathbf{z}) + k_t(\mathbf{z},\mathbf{z}_\mathrm{n})p_t(\mathbf{z}_\mathrm{n}), \tag{SI-15}$$

where $r_{2,t}(\mathbf{z})$ is the conditional p.d.f for the event that, given a focal individual has phenotype $\mathbf{z}$ at time $t$, a randomly sampled individual among its neighbours is IBD to this focal (this p.d.f. depends on the whole phenotypic distribution, $\phi_t$, which is captured by the time index in $r_{2,t}(\mathbf{z})$). We refer to $r_{2,t}(\mathbf{z})$ as pairwise relatedness. The two summands in eq. (SI-15) respectively capture two complementary events: the sampled neighbour is either (1) IBD with the focal (and thus must have the same phenotype as the focal, $\mathbf{z}_\mathrm{n} = \mathbf{z}$); or (2) not IBD with the focal and has phenotype $\mathbf{z}_\mathrm{n}$ (which may or may not be equal to $\mathbf{z}$). We have written the p.d.f. for this latter event as $k_t(\mathbf{z},\mathbf{z}_\mathrm{n})p_t(\mathbf{z}_\mathrm{n})$, where $p_t(\mathbf{z}_\mathrm{n})$ is the marginal probability of sampling an individual with phenotype $\mathbf{z}_\mathrm{n}$ from the global population. The function $k_t(\mathbf{z},\mathbf{z}_\mathrm{n})$ can therefore be viewed as the multiplicative effect of having already sampled an individual with phenotype $\mathbf{z}$ in the group on this marginal probability.

In our endeavour to obtain an expression for lineage fitness up to the order $\xi_t^2$, we seek to express $q_{2,t}(\mathbf{z}_\mathrm{n}|\mathbf{z})$ to the order of $\xi_t$ (as it multiplies $\xi_t(z_{\mathrm{n},a})$, which is of order $\xi_t$, in eq. SI-13). We thus Taylor expand both $r_{2,t}(\mathbf{z})$ and $k_t(\mathbf{z},\mathbf{z}_\mathrm{n})$ in eq. (SI-15) to the first-order around $\bar{\mathbf{z}}_t$, and obtain

$$\begin{aligned} q_{2,t}(\mathbf{z}_\mathrm{n}|\mathbf{z}) = &\left(r^\circ_{2,t}(\bar{\mathbf{z}}_t) + \sum_{a=1}^{n}\frac{\partial r_{2,t}(\mathbf{z})}{\partial z_a}\xi_t(z_a)\right)\delta(\mathbf{z}_\mathrm{n} - \mathbf{z}) \\ &+ \left(k_t(\bar{\mathbf{z}}_t,\bar{\mathbf{z}}_t) + \sum_{a=1}^{n}\frac{\partial k_t(\mathbf{z},\mathbf{z}_\mathrm{n})}{\partial z_a}\xi_t(z_a) + \sum_{a=1}^{n}\frac{\partial k_t(\mathbf{z},\mathbf{z}_\mathrm{n})}{\partial z_{\mathrm{n},a}}\xi_t(z_{\mathrm{n},a})\right) p_t(\mathbf{z}_\mathrm{n}) + \mathcal{O}(\xi_t^2), \end{aligned} \tag{SI-16}$$

where $r^\circ_{2,t}(\bar{\mathbf{z}}_t)$ is the probability that two randomly sampled individuals within a group at time $t$ are IBD under neutrality (i.e., when the population is monomorphic for $\bar{\mathbf{z}}_t$).

Next, we use the fact that from the definition of $q_{2,t}(\mathbf{z}_\mathrm{n}|\mathbf{z})$ (eq. SI-14), $q_{2,t}(\mathbf{z}_\mathrm{n}|\mathbf{z})p_t(\mathbf{z})$ is the (unconditional) p.d.f. for the event of sampling one individual with phenotype $\mathbf{z}$ and another with $\mathbf{z}_\mathrm{n}$ *without* replacement from a group. Thus, three identities must hold:

$$\begin{aligned}\int_{\mathbb{R}^n} q_{2,t}(\mathbf{z}_\mathrm{n}|\mathbf{z})p_t(\mathbf{z})\mathrm{d}\mathbf{z}_\mathrm{n} &= p_t(\mathbf{z}) &\text{(SI-17)}\\ \int_{\mathbb{R}^n} q_{2,t}(\mathbf{z}_\mathrm{n}|\mathbf{z})p_t(\mathbf{z})\mathrm{d}\mathbf{z} &= p_t(\mathbf{z}_\mathrm{n}), &\text{(SI-18)}\\ \int_{\mathbb{R}^n}\int_{\mathbb{R}^n} q_{2,t}(\mathbf{z}_\mathrm{n}|\mathbf{z})p_t(\mathbf{z})\mathrm{d}\mathbf{z}_\mathrm{n}\mathrm{d}\mathbf{z} &= 1, &\text{(SI-19)}\end{aligned}$$

which can be used to express $k_t(\mathbf{z},\mathbf{z}_\mathrm{n})$ in terms of $r_{2,t}(\mathbf{z})$. In fact, substituting eq. (SI-16) into eq. (SI-19), we obtain

$$r^\circ_{2,t}(\bar{\mathbf{z}}_t) + k_t(\bar{\mathbf{z}}_t,\bar{\mathbf{z}}_t) = 1 \Rightarrow k_t(\bar{\mathbf{z}}_t,\bar{\mathbf{z}}_t) = 1 - r^\circ_{2,t}(\bar{\mathbf{z}}_t). \tag{SI-20a}$$

Similarly, substituting eq. (SI-20a) into eq. (SI-16), which is in turn substituted into eqs. (SI-20b) and (SI-20c), yields,

$$p_t(\mathbf{z}) + \sum_{a=1}^{n}\frac{\partial r_{2,t}(\mathbf{z})}{\partial z_a}\xi_t(z_a)p_t(\mathbf{z}) + \sum_{a=1}^{n}\frac{\partial k_t(\mathbf{z},\mathbf{z}_\mathrm{n})}{\partial z_a}\xi_t(z_a)p_t(\mathbf{z}) = p_t(\mathbf{z}) \Rightarrow \frac{\partial k_t(\mathbf{z},\mathbf{z}_\mathrm{n})}{\partial z_a} = \frac{\partial r_{2,t}(\mathbf{z})}{\partial z_a} \tag{SI-20b}$$

and

$$p_t(\mathbf{z}_\mathrm{n}) + \sum_{a=1}^{n}\frac{\partial r_{2,t}(\mathbf{z})}{\partial z_a}\xi_t(z_{\mathrm{n},a})p_t(\mathbf{z}_\mathrm{n}) + \sum_{a=1}^{n}\frac{\partial k_t(\mathbf{z},\mathbf{z}_\mathrm{n})}{\partial z_{\mathrm{n},a}}\xi_t(z_{\mathrm{n},a})p_t(\mathbf{z}_\mathrm{n}) = p_t(\mathbf{z}_\mathrm{n}) \Rightarrow \frac{\partial k_t(\mathbf{z},\mathbf{z}_\mathrm{n})}{\partial z_{\mathrm{n},a}} = \frac{\partial r_{2,t}(\mathbf{z})}{\partial z_a}, \tag{SI-20c}$$

respectively. Finally, substituting eqs. (SI-20) into eq. (SI-16), we obtain,

$$q_{2,t}(\mathbf{z}_\mathrm{n}|\mathbf{z}) = q^\circ_{2,t}(\mathbf{z}_\mathrm{n}|\mathbf{z}) + \left(\sum_{a=1}^{n}\xi_t(z_a)\frac{\partial r_{2,t}(\mathbf{z})}{\partial z_a}\right)\delta(\mathbf{z}_\mathrm{n}-\mathbf{z}) - \left(\sum_{a=1}^{n}(\xi_t(z_a)+\xi_t(z_{\mathrm{n},a}))\frac{\partial r_{2,t}(\mathbf{z})}{\partial z_a}\right)p_t(\mathbf{z}_\mathrm{n}) + \mathcal{O}(\xi_t^2), \tag{SI-21a}$$

where

$$q^\circ_{2,t}(\mathbf{z}_\mathrm{n}|\mathbf{z}) = r^\circ_{2,t}(\bar{\mathbf{z}}_t)\delta(\mathbf{z}_\mathrm{n}-\mathbf{z}) + \left(1-r^\circ_{2,t}(\bar{\mathbf{z}}_t)\right)p_t(\mathbf{z}_\mathrm{n}), \tag{SI-21b}$$

is the neutral (conditional) p.d.f. for the event of sampling an individual with phenotype $\mathbf{z}_\mathrm{n}$ within the neighbourhood of an individual with phenotype $\mathbf{z}$.

##### 1.1.2 Regression definition of relatedness

It is noteworthy that the definition of pairwise relatedness as the probability of IBD between two randomly sampled individuals within a group that we use, aligns with the "regression definition of relatedness" (e.g., Grafen, 1985, eq. 2.13 of Frank, 1998). Under this latter definition, relatedness is the regression of neighbour phenotype on focal phenotype: it is the ratio of phenotypic covariance among neighbours,

$$\varphi_{ab,t} = \int_{\mathbb{R}^n}\int_{\mathbb{R}^n}\xi_t(z_a)\xi_t(z_{\mathrm{n},b})q_{2,t}(\mathbf{z}_\mathrm{n}|\mathbf{z})p_t(\mathbf{z})\mathrm{d}\mathbf{z}_\mathrm{n}\mathrm{d}\mathbf{z}, \tag{SI-22}$$

to the covariance within individuals, $\sigma_{ab,t} = \int_{\mathbb{R}^n}\xi_t(z_a)\xi_t(z_b)p_t(\mathbf{z})\mathrm{d}\mathbf{z}$. Substituting eq. (SI-21) into eq. (SI-22) to compute this ratio, we obtain

$$\frac{\varphi_{ab,t}}{\sigma_{ab,t}} = r^\circ_{2,t}(\bar{\mathbf{z}}_t) + \sum_{c=1}^{n}\frac{\kappa_{abc,t}}{\sigma_{ab,t}}\frac{\partial r_{2,t}(\mathbf{z})}{\partial z_c} + \mathcal{O}(\xi_t^2) \tag{SI-23}$$

where $\kappa_{abc,t} = \int_{\mathbb{R}^n}\xi_t(z_a)\xi_t(z_b)\xi_t(z_c)p_t(\mathbf{z})\mathrm{d}\mathbf{z}$ is the skew between traits $a$, $b$ and $c$ within individuals. Our eq. (SI-23) is thus in line with the result that in the infinite island model and under neutrality, relatedness as a regression coefficient is equal to the probability of identity-by-descent (i.e., $\varphi_{ab,t}/\sigma_{ab,t} = r^\circ_{2,t}(\bar{\mathbf{z}}_t) + \mathcal{O}(\xi_t)$, Rousset, 2002; for equivalent expressions to eq. SI-23, see for e.g. eq. 7 of Queller, 1992; eq. 2.13 of Frank, 1998; eq. 11 of Wakano and Lehmann, 2014).

##### 1.1.3 Average marginal effects

We can now return to our calculation of the average marginal effects of a change in one trait (eq. SI-13). Substituting eq. (SI-21) into eq. (SI-13), we in fact obtain

$$\begin{aligned}\int_{\mathcal{S}} \beta_{a,t} q_t(\mu|\mathbf{z})\mathrm{d}\mu &= \xi_t(z_a)\left(\frac{\partial w}{\partial z_a} + (N-1) r^{\circ}_{2,t}(\bar{\mathbf{z}}_t)\frac{\partial w}{\partial z_{\mathrm{n},a}}\right) \\ &+ \sum_{b=1}^{n} \left(\xi_t(z_a)\xi_t(z_b) - \sigma_{ab,t}\right)\frac{\partial r_{2,t}(\mathbf{z})}{\partial z_b}(N-1)\frac{\partial w}{\partial z_{\mathrm{n},a}} + \mathcal{O}(\xi_t^3),\end{aligned} \tag{SI-24}$$

for the average marginal effect of trait $a$.

#### 2 Interaction effects

We now tackle the interaction effects on focal fitness, $\beta_{ab,t}$ (eq. SI-3), which we will then average over $q_t(\mu|\mathbf{z})$. Using notation eqs. (SI-5)-(SI-6), we first decompose $\beta_{ab,t}$ as

$$\beta_{ab,t} = \beta^{\text{same}}_{ab,t} + \beta^{\text{neigh}}_{ab,t} + \beta^{\text{cross isle}}_{ab,t}, \tag{SI-25}$$

where

$$\beta^{\text{same}}_{ab,t} = \xi_t(z_a)\xi_t(z_b)\frac{\partial^2 w}{\partial z_a \partial z_b} + \sum_{\mathbf{z}_{\mathrm{n}} \in \mathrm{n}_\mu(\mathbf{z})} \xi_t(z_{\mathrm{n},a})\xi_t(z_{\mathrm{n},b})\frac{\partial^2 w}{\partial z_{\mathrm{n},a}\partial z_{\mathrm{n},b}} + \sum_{\mathbf{z}_{\mathrm{nn}} \in \mathrm{e}(\phi_t)} \xi_t(z_{\mathrm{nn},a})\xi_t(z_{\mathrm{nn},b})\frac{\partial^2 w}{\partial z_{\mathrm{nn},a}\partial z_{\mathrm{nn},b}}, \tag{SI-26}$$

collects the interaction effects of traits $a$ and $b$ within individuals,

$$\begin{aligned}\beta^{\text{neigh}}_{ab,t} &= \sum_{\mathbf{z}_{\mathrm{n}} \in \mathrm{n}_\mu(\mathbf{z})} \xi_t(z_a)\xi_t(z_{\mathrm{n},b})\frac{\partial^2 w}{\partial z_a \partial z_{\mathrm{n},b}} + \sum_{\mathbf{z}_{\mathrm{n}} \in \mathrm{n}_\mu(\mathbf{z})} \xi_t(z_b)\xi_t(z_{\mathrm{n},a})\frac{\partial^2 w}{\partial z_b \partial z_{\mathrm{n},a}} \\ &+ \sum\sum_{\mathbf{z}_{\mathrm{n}},\mathbf{z}_{\mathrm{m}} \in \mathrm{n}_\mu(\mathbf{z})} \xi_t(z_{\mathrm{n},a})\xi_t(z_{\mathrm{m},b})\frac{\partial^2 w}{\partial z_{\mathrm{n},a}\partial z_{\mathrm{m},b}} + \sum\sum_{\mathbf{z}_{\mathrm{nn}},\mathbf{z}_{\mathrm{nm}} \in \mathrm{e}(\phi_t)} \xi_t(z_{\mathrm{nn},a})\xi_t(z_{\mathrm{nm},b})\frac{\partial^2 w}{\partial z_{\mathrm{nn},a}\partial z_{\mathrm{nm},b}},\end{aligned} \tag{SI-27}$$

collects the interaction effects between neighbours, and

$$\begin{aligned}\beta^{\text{cross isle}}_{ab,t} &= \sum_{\mathbf{z}_{\mathrm{nn}} \in \mathrm{e}(\phi_t)} \xi_t(z_a)\xi_t(z_{\mathrm{nn},b})\frac{\partial^2 w}{\partial z_a \partial z_{\mathrm{nn},b}} + \sum_{\mathbf{z}_{\mathrm{nn}} \in \mathrm{e}(\phi_t)} \xi_t(z_b)\xi_t(z_{\mathrm{nn},a})\frac{\partial^2 w}{\partial z_b \partial z_{\mathrm{nn},a}} \\ &+ \sum\sum_{\substack{\mathbf{z}_{\mathrm{n}} \in \mathrm{n}_\mu(\mathbf{z}) \\ \mathbf{z}_{\mathrm{nn}} \in \mathrm{e}(\phi_t)}} \xi_t(z_{\mathrm{n},a})\xi_t(z_{\mathrm{nn},b})\frac{\partial^2 w}{\partial z_{\mathrm{n},a}\partial z_{\mathrm{nn},b}} + \sum\sum_{\substack{\mathbf{z}_{\mathrm{n}} \in \mathrm{n}_\mu(\mathbf{z}) \\ \mathbf{z}_{\mathrm{nn}} \in \mathrm{e}(\phi_t)}} \xi_t(z_{\mathrm{n},b})\xi_t(z_{\mathrm{nn},a})\frac{\partial^2 w}{\partial z_{\mathrm{n},b}\partial z_{\mathrm{nn},a}},\end{aligned} \tag{SI-28}$$

the interaction effects between individuals between groups. We proceed to specify each of these in terms of counting measures and average them over $q_t(\mu|\mathbf{z})$ sequentially.

##### 2.1 Interaction effects within individuals

We start with interaction effects within individuals, $\beta^{\text{same}}_{ab,t}$. Using eq. (SI-10), eq. (SI-26) can be expressed as

$$\begin{aligned}\beta^{\text{same}}_{ab,t} &= \xi_t(z_a)\xi_t(z_b)\frac{\partial^2 w}{\partial z_a \partial z_b} + \left(\int_{\mathbb{R}^n} \xi_t(z_{\mathrm{n},a})\xi_t(z_{\mathrm{n},b})\mu_{\mathrm{n},\mathbf{z}}(\mathbf{z}_{\mathrm{n}})\mathrm{d}\mathbf{z}_{\mathrm{n}}\right)\frac{\partial^2 w}{\partial z_{\mathrm{n},a}\partial z_{\mathrm{n},b}} \\ &+ \left(\int_{\mathbb{R}^n}\int_{\mathcal{S}} \xi_t(z_{\mathrm{nn},a})\xi_t(z_{\mathrm{nn},b})\mu(\mathbf{z}_{\mathrm{nn}})\phi_t(\mu)\mathrm{d}\mathbf{z}_{\mathrm{nn}}\mathrm{d}\mu\right)\frac{\partial^2 w}{\partial z_{\mathrm{nn},a}\partial z_{\mathrm{nn},b}},\end{aligned} \tag{SI-29}$$

where the first summand consists of the interaction effect within the focal individual, the second within neighbours of the focal, and the third within individuals from other groups than the focal. To simplify these, we use the fact that because the population size remains constant, we have

$$\frac{\partial^2 w}{\partial z_a \partial z_b} + (N-1)\frac{\partial^2 w}{\partial z_{\mathrm{n},a}\partial z_{\mathrm{n},b}} + N\frac{\partial^2 w}{\partial z_{\mathrm{nn},a}\partial z_{\mathrm{nn},b}} = 0 \tag{SI-30}$$

(see eq. B.13 of Wakano and Lehmann, 2014); and that the third term of eq. (SI-29) can be expressed as

$$\int_{\mathbb{R}^n}\int_{\mathcal{S}} \xi_t(z_{\mathrm{nn},a})\xi_t(z_{\mathrm{nn},b})\mu(\mathbf{z}_{\mathrm{nn}})\phi_t(\mu)\mathrm{d}\mu\mathrm{d}\mathbf{z}_{\mathrm{nn}} = N\int_{\mathbb{R}^n} \xi_t(z_{\mathrm{nn},a})\xi_t(z_{\mathrm{nn},b})p_t(\mathbf{z}_{\mathrm{nn},a})\mathrm{d}\mathbf{z}_{\mathrm{nn}} = N\sigma_{ab,t}. \tag{SI-31}$$

Substituting eqs. (SI-30) and (SI-31) into eq. (SI-29), we find that the interaction effects of $a$ and $b$ within individuals can be written as

$$\beta^{\mathrm{same}}_{ab,t} = (\xi_t(z_a)\xi_t(z_b) - \sigma_{ab,t})\frac{\partial^2 w}{\partial z_a\partial z_b} + \left(\int_{\mathbb{R}^n} \xi_t(z_{\mathrm{n},a})\xi_t(z_{\mathrm{n},b})\frac{\mu_{\mathrm{n},\mathbf{z}}(\mathbf{z}_\mathrm{n})}{N-1}\mathrm{d}\mathbf{z}_\mathrm{n} - \sigma_{ab,t}\right)(N-1)\frac{\partial^2 w}{\partial z_{\mathrm{n},a}\partial z_{\mathrm{n},b}}. \tag{SI-32}$$

Averaging these interaction effects within individuals over $q_t(\mu|\mathbf{z})$ then gives

$$\int_{\mathcal{S}} \beta^{\mathrm{same}}_{ab,t} q_t(\mu|\mathbf{z})\mathrm{d}\mu = (\xi_t(z_a)\xi_t(z_b) - \sigma_{ab,t})\left(\frac{\partial^2 w}{\partial z_a\partial z_b} + (N-1)r^{\circ}_{2,t}(\bar{\mathbf{z}}_t)\frac{\partial^2 w}{\partial z_{\mathrm{n},a}\partial z_{\mathrm{n},b}}\right) + \mathcal{O}(\xi_t^3), \tag{SI-33}$$

where we used definition eq. (SI-14) and eq. (SI-21).

#### 2.2 Interaction effects between neighbours

Let us now turn to interaction effects between neighbours, $\beta^{\mathrm{neigh}}_{ab,t}$ (eq. SI-27). It is composed of four summands: the first two capture the interaction effects between the focal and its neighbours; the third, between two neighbours of the focal; and the fourth, between two neighbours from another group than the focal. We consider these separately below.

##### 2.2.1 Interaction effects between the focal and its neighbours

The first summand of $\beta^{\mathrm{neigh}}_{ab,t}$ (eq. SI-27) corresponds to the interaction effect between trait $a$ in the focal and trait $b$ in its neighbours. Using eq. (SI-10), it can be expressed as

$$\sum_{\mathbf{z}_\mathrm{n}\in \mathrm{n}_\mu(\mathbf{z})} \xi_t(z_a)\xi_t(z_{\mathrm{n},b})\frac{\partial^2 w}{\partial z_a\partial z_{\mathrm{n},b}} = \left(\int_{\mathbb{R}^n} \xi_t(z_a)\xi_t(z_{\mathrm{n},b})\mu_{\mathrm{n},\mathbf{z}}(\mathbf{z}_\mathrm{n})\mathrm{d}\mathbf{z}_\mathrm{n}\right)\frac{\partial^2 w}{\partial z_a\partial z_{\mathrm{n},b}}. \tag{SI-34}$$

Averaging eq. (SI-34) over $q_t(\mu|\mathbf{z})$ then reads as

$$\int_{\mathbb{R}^n} \xi_t(z_a)\xi_t(z_{\mathrm{n},b})q_{2,t}(\mathbf{z}_\mathrm{n}|\mathbf{z})\mathrm{d}\mathbf{z}_\mathrm{n}(N-1)\frac{\partial^2 w}{\partial z_a\partial z_{\mathrm{n},b}} = \xi_t(z_a)\xi_t(z_b)r^{\circ}_{2,t}(\bar{\mathbf{z}}_t)(N-1)\frac{\partial^2 w}{\partial z_a\partial z_{\mathrm{n},b}} + \mathcal{O}(\xi_t^3), \tag{SI-35}$$

where we used eq. (SI-21). Similarly, averaging the second summand of $\beta^{\mathrm{neigh}}_{ab,t}$ (eq. SI-27), which corresponds to the interaction effect between trait $b$ in the focal and trait $a$ in its neighbours, yields

$$\int_{\mathcal{S}}\left[\sum_{\mathbf{z}_\mathrm{n}\in \mathrm{n}_\mu(\mathbf{z})} \xi_t(z_b)\xi_t(z_{\mathrm{n},a})\frac{\partial^2 w}{\partial z_b\partial z_{\mathrm{n},a}}\right]q_t(\mu|\mathbf{z})\mathrm{d}\mu = \xi_t(z_a)\xi_t(z_b)r^{\circ}_{2,t}(\bar{\mathbf{z}}_t)(N-1)\frac{\partial^2 w}{\partial z_b\partial z_{\mathrm{n},a}} + \mathcal{O}(\xi_t^3). \tag{SI-36}$$

##### 2.2.2 Interaction effects between neighbours of the focal

The third summand of $\beta^{\mathrm{neigh}}_{ab,t}$ (eq. SI-27) collects the interaction effects between neighbours of the focal. To express these in terms of the counting measure $\mu(\mathbf{z})$ and average them over $q_t(\mu|\mathbf{z})$, we introduce one further conditional counting measure,

$$\begin{aligned}\mu_{\mathrm{n},\mathbf{z}}(\mathbf{z}_{\mathrm{n}},\mathbf{z}_{\mathrm{m}}) &= \delta(\mathbf{z}_{\mathrm{n}}-\mathbf{z})\,\delta(\mathbf{z}_{\mathrm{m}}-\mathbf{z}_{\mathrm{n}})\,(\mu(\mathbf{z})-1)(\mu(\mathbf{z})-2)\\ &+\delta(\mathbf{z}_{\mathrm{n}}-\mathbf{z})\,(1-\delta(\mathbf{z}_{\mathrm{m}}-\mathbf{z}))\,(\mu(\mathbf{z})-1)\mu(\mathbf{z}_{\mathrm{m}})+\delta(\mathbf{z}_{\mathrm{m}}-\mathbf{z})\,(1-\delta(\mathbf{z}_{\mathrm{n}}-\mathbf{z}))\,(\mu(\mathbf{z})-1)\mu(\mathbf{z}_{\mathrm{n}})\\ &+\delta(\mathbf{z}_{\mathrm{m}}-\mathbf{z}_{\mathrm{n}})\,(1-\delta(\mathbf{z}_{\mathrm{n}}-\mathbf{z}))\,\mu(\mathbf{z}_{\mathrm{n}})\,(\mu(\mathbf{z}_{\mathrm{n}})-1)\\ &+(1-\delta(\mathbf{z}_{\mathrm{n}}-\mathbf{z}))\,(1-\delta(\mathbf{z}_{\mathrm{m}}-\mathbf{z}))\,(1-\delta(\mathbf{z}_{\mathrm{n}}-\mathbf{z}_{\mathrm{m}}))\,\mu(\mathbf{z}_{\mathrm{n}})\mu(\mathbf{z}_{\mathrm{m}})\end{aligned} \tag{SI-37}$$

which counts the number of unordered pairs of neighbours that have phenotypes $\mathbf{z}_{\mathrm{n}}$ and $\mathbf{z}_{\mathrm{m}}$, given that the focal individual has phenotype $\mathbf{z}$. Integrated over $\mathbb{R}^n$, the first line of eq. (SI-37) counts the number of pairs of neighbours that have focal phenotype $\mathbf{z}$; the second line, the number of pairs of neighbours in which only one has the focal phenotype $\mathbf{z}$; the third line, the number of pairs in which both neighbours have the same phenotype that is different to the the focal; and the fourth line, the number of pairs in which neighbours have phenotypes that are different to one another, and to the focal.

Using eq. (SI-37), we can then re-write the third summand of $\beta_{ab,t}^{\mathrm{neigh}}$ as

$$\sum_{\mathbf{z}_{\mathrm{n}},\mathbf{z}_{\mathrm{m}}\in\mathrm{n}_\mu(\mathbf{z})}\sum \xi_t(z_{\mathrm{n},a})\xi_t(z_{\mathrm{m},b})\frac{\partial^2 w}{\partial z_{\mathrm{n},a}\partial z_{\mathrm{m},b}} = \left(\int_{\mathbb{R}^n}\int_{\mathbb{R}^n}\mu_{\mathrm{n},\mathbf{z}}(\mathbf{z}_{\mathrm{n}},\mathbf{z}_{\mathrm{m}})\mathrm{d}\mathbf{z}_{\mathrm{n}}\mathrm{d}\mathbf{z}_{\mathrm{m}}\right)\frac{\partial^2 w}{\partial z_{\mathrm{n},a}\partial z_{\mathrm{m},b}}, \tag{SI-38}$$

which averaged over $q_t(\mu|\mathbf{z})$ reads as

$$\left[\int_{\mathbb{R}^n}\int_{\mathbb{R}^n}\xi_t(z_{\mathrm{n},a})\xi_t(z_{\mathrm{m},b})q_{3,t}(\mathbf{z}_{\mathrm{n}},\mathbf{z}_{\mathrm{m}}|\mathbf{z})\mathrm{d}\mathbf{z}_{\mathrm{n}}\mathrm{d}\mathbf{z}_{\mathrm{m}}\right](N-1)(N-2)\frac{\partial^2 w}{\partial z_{\mathrm{n},a}\partial z_{\mathrm{m},b}}, \tag{SI-39}$$

where

$$q_{3,t}(\mathbf{z}_{\mathrm{n}},\mathbf{z}_{\mathrm{m}}|\mathbf{z}) = \int_{\mathcal{S}}\frac{\mu_{\mathrm{n},\mathbf{z}}(\mathbf{z}_{\mathrm{n}},\mathbf{z}_{\mathrm{m}})}{(N-1)(N-2)}q_t(\mu|\mathbf{z})\mathrm{d}\mu \tag{SI-40}$$

is the conditional p.d.f. for the event of sampling two individuals without replacement, one with phenotype $\mathbf{z}_{\mathrm{n}}$ and another with phenotype $\mathbf{z}_{\mathrm{m}}$ in a group, given that a focal individual with phenotype $\mathbf{z}$ has already been sampled in that group at time $t$. This p.d.f. can be connected to notions of relatedness as individuals that have the same phenotype may be IBD. In fact, using a coalescence argument (see eq. B.10 of Wakano and Lehmann, 2014 for e.g.), we can rewrite $q_{3,t}(\mathbf{z}_{\mathrm{n}},\mathbf{z}_{\mathrm{m}}|\mathbf{z})$ as

$$\begin{aligned}q_{3,t}(\mathbf{z}_{\mathrm{n}},\mathbf{z}_{\mathrm{m}}|\mathbf{z}) = {}&\delta(\mathbf{z}_{\mathrm{n}}-\mathbf{z})\,\delta(\mathbf{z}_{\mathrm{m}}-\mathbf{z})\,r_{3,t}^{\circ}(\bar{\mathbf{z}}_t)\\ &+\delta(\mathbf{z}_{\mathrm{n}}-\mathbf{z})\,(r_{2,t}^{\circ}(\bar{\mathbf{z}}_t)-r_{3,t}^{\circ}(\bar{\mathbf{z}}_t))p_t(\mathbf{z}_{\mathrm{m}})+\delta(\mathbf{z}_{\mathrm{m}}-\mathbf{z})\,(r_{2,t}^{\circ}(\bar{\mathbf{z}}_t)-r_{3,t}^{\circ}(\bar{\mathbf{z}}_t))p_t(\mathbf{z}_{\mathrm{n}})\\ &+\delta(\mathbf{z}_{\mathrm{n}}-\mathbf{z}_{\mathrm{m}})\,(r_{2,t}^{\circ}(\bar{\mathbf{z}}_t)-r_{3,t}^{\circ}(\bar{\mathbf{z}}_t))p_t(\mathbf{z}_{\mathrm{n}})\\ &+(1-3r_{2,t}^{\circ}(\bar{\mathbf{z}}_t)+2r_{3,t}^{\circ}(\bar{\mathbf{z}}_t))p_t(\mathbf{z}_{\mathrm{n}})p_t(\mathbf{z}_{\mathrm{m}})+\mathcal{O}(\xi_t),\end{aligned} \tag{SI-41}$$

where $r_{3,t}^{\circ}(\bar{\mathbf{z}}_t)$ is the probability of sampling three individuals without replacement from a group are IBD in the absence of selection at time $t$ (i.e., three-way relatedness). Each summand of eq. (SI-41) capture a different possible relationship among the sampled ($\mathbf{z}_{\mathrm{n}}$ and $\mathbf{z}_{\mathrm{m}}$) and focal ($\mathbf{z}$) phenotype. The first line of eq. (SI-41) captures the event of sampling two individuals that are both IBD with the focal (and thus with the same phenotype as the focal, $\mathbf{z}=\mathbf{z}_{\mathrm{n}}=\mathbf{z}_{\mathrm{m}}$); the next line, the event of sampling only one individual IBD with the focal (and thus with the same phenotype: the two summands respectively capture the cases $\mathbf{z}=\mathbf{z}_{\mathrm{n}}$ and $\mathbf{z}=\mathbf{z}_{\mathrm{m}}$); the third line, the event of sampling two individuals that are IBD together (and thus with phenotypes $\mathbf{z}_{\mathrm{n}}=\mathbf{z}_{\mathrm{m}}$) but not with the focal; and the last line, the event of sampling two individuals that are not IBD with one another or with the focal.

Substituting eq. (SI-41) into eq. (SI-39) then gives us

$$\left[\xi_t(z_a)\xi_t(z_b)r_{3,t}^{\circ}(\bar{\mathbf{z}}_t)+\sigma_{ab,t}(r_{2,t}^{\circ}(\bar{\mathbf{z}}_t)-r_{3,t}^{\circ}(\bar{\mathbf{z}}_t))\right](N-1)(N-2)\frac{\partial^2 w}{\partial z_{\mathrm{n},a}\partial z_{\mathrm{m},b}}+\mathcal{O}(\xi_t^3) \tag{SI-42}$$

for the average interaction effects between neighbours of the focal.

##### 2.2.3 Interaction effects between neighbours from other groups

The fourth and final summand of $\beta^{\text{neigh}}_{ab,t}$ (eq. SI-27), which collects the interaction effects between neighbours from other groups, can be expressed using eq. (SI-10) as

$$
\begin{aligned}
&\sum\sum_{\mathbf{z}_{\text{nn}},\mathbf{z}_{\text{nm}}\in e(\phi_t)} \xi_t(z_{\text{nn},a})\xi_t(z_{\text{nm},b})\frac{\partial^2 w}{\partial z_{\text{nn},a}\partial z_{\text{nm},b}} \\
&= N(N-1)\underbrace{\left(\int_{\mathcal{S}}\int_{\mathbb{R}^n}\int_{\mathbb{R}^n} \xi_t(z_{\text{nn},a})\xi_t(z_{\text{nm},b})\frac{\mu_{\text{n},\mathbf{z}_{\text{nn}}}(\mathbf{z}_{\text{nm}})\mu(\mathbf{z}_{\text{nn}})}{N(N-1)}\phi_t(\mu)\text{d}\mathbf{z}_{\text{nm}}\text{d}\mathbf{z}_{\text{nn}}\text{d}\mu\right)}_{\varphi_{ab,t}}\frac{\partial^2 w}{\partial z_{\text{nn},a}\partial z_{\text{nm},b}},
\end{aligned}
\tag{SI-43}
$$

where $\varphi_{ab,t}$ is the covariance among traits $a$ and $b$ between individuals within groups (see eq. SI-22). We can then use eq. (SI-23) to specify this covariance and obtain,

$$
\sum\sum_{\mathbf{z}_{\text{nn}},\mathbf{z}_{\text{nm}}\in e(\phi_t)} \xi_t(z_{\text{nn},a})\xi_t(z_{\text{nm},b})\frac{\partial^2 w}{\partial z_{\text{nn},a}\partial z_{\text{nm},b}} = N(N-1)r^{\circ}_{2,t}(\bar{\mathbf{z}}_t)\sigma_{ab,t}\frac{\partial^2 w}{\partial z_{\text{nn},a}\partial z_{\text{nm},b}} + \mathcal{O}(\xi_t^3). \tag{SI-44}
$$

Since this expression does not depend on group composition $\mu$, it will be invariant to averaging over $q_t(\mu|\mathbf{z})$.

##### 2.2.4 Average interaction effects between neighbours.

Hence, the average interaction effects between neighbours ($\int_{\mathcal{S}}\beta^{\text{neigh}}_{ab,t}q_t(\mu|\mathbf{z})\text{d}\mu$), which is given by the sum of eqs. (SI-35), (SI-36), (SI-42) and (SI-44), reads as

$$
\begin{aligned}
\int_{\mathcal{S}}\beta^{\text{neigh}}_{ab,t}q_t(\mu|\mathbf{z})\text{d}\mu &= \xi_t(z_a)\xi_t(z_b)r^{\circ}_{2,t}(\bar{\mathbf{z}}_t)(N-1)\left(\frac{\partial^2 w}{\partial z_a\partial z_{\text{n},b}}+\frac{\partial^2 w}{\partial z_b\partial z_{\text{n},a}}\right) \\
&+\left[\xi_t(z_a)\xi_t(z_b)r^{\circ}_{3,t}(\bar{\mathbf{z}}_t)+\sigma_{ab,t}(r^{\circ}_{2,t}(\bar{\mathbf{z}}_t)-r^{\circ}_{3,t}(\bar{\mathbf{z}}_t))\right](N-1)(N-2)\frac{\partial^2 w}{\partial z_{\text{n},a}\partial z_{\text{m},b}} \\
&+N(N-1)r^{\circ}_{2,t}(\bar{\mathbf{z}}_t)\sigma_{ab,t}\frac{\partial^2 w}{\partial z_{\text{nn},a}\partial z_{\text{nm},b}}+\mathcal{O}(\xi_t^3).
\end{aligned}
\tag{SI-45}
$$

This can be further simplified by using the fact than since the total population size is constant, the following holds,

$$
(N-1)\frac{\partial^2 w}{\partial z_a\partial z_{\text{n},b}}+(N-1)\frac{\partial^2 w}{\partial z_b\partial z_{\text{n},a}}+(N-1)(N-2)\frac{\partial^2 w}{\partial z_{\text{n},a}\partial z_{\text{m},b}}+N(N-1)\frac{\partial^2 w}{\partial z_{\text{nn},a}\partial z_{\text{nm},b}}=0. \tag{SI-46}
$$

(see eq. B.14 of Wakano and Lehmann, 2014 for argument). Substituting for $\partial^2 w/(\partial z_{\text{nn},a}\partial z_{\text{nm},b})$ using eq. (SI-46) into eq. (SI-45) finally gives

$$
\begin{aligned}
\int_{\mathcal{S}}\beta^{\text{neigh}}_{ab,t}q_t(\mu|\mathbf{z})\text{d}\mu &= (\xi_t(z_a)\xi_t(z_b)-\sigma_{ab,t})(N-1) \\
&\times\left(r^{\circ}_{2,t}(\bar{\mathbf{z}}_t)\left(\frac{\partial^2 w}{\partial z_a\partial z_{\text{n},b}}+\frac{\partial^2 w}{\partial z_b\partial z_{\text{n},a}}\right)+(N-2)r^{\circ}_{3,t}(\bar{\mathbf{z}}_t)\frac{\partial^2 w}{\partial z_{\text{n},a}\partial z_{\text{m},b}}\right)+\mathcal{O}(\xi_t^3).
\end{aligned}
\tag{SI-47}
$$

#### 2.3 Interaction effects between individuals between groups

The final relevant interaction effect, $\beta^{\text{cross isle}}_{ab,t}$ (eq. SI-28), is the interaction between individuals that belong to different groups. However, it is straightforward to show that these effects vanish in the infinite island model of dispersal. For example, consider the first summand of eq. (SI-28), which measures the effect of a change in trait

$a$ of the focal and a trait $b$ in an individual from a group other than the focal:

$$\sum_{\mathbf{z}_{\mathrm{nn}}\in \mathrm{e}(\phi_t)} \xi_t(z_a)\xi_t(z_{\mathrm{nn},b})\frac{\partial^2 w}{\partial z_a \partial z_{\mathrm{nn},b}} = \left(\int_{\mathbb{R}^n}\int_{\mathcal{S}} \xi_t(z_a)\xi_t(z_{\mathrm{nn},b})\mu(\mathbf{z}_{\mathrm{nn}})\phi_t(\mu)\mathrm{d}\mu\mathrm{d}\mathbf{z}_{\mathrm{nn}}\right)\frac{\partial^2 w}{\partial z_a \partial z_{\mathrm{nn},b}}$$
$$= \left(N\xi_t(z_a)\int_{\mathbb{R}^n} \xi_t(z_{\mathrm{nn},b})p_t(\mathbf{z}_{\mathrm{nn}})\mathrm{d}\mathbf{z}_{\mathrm{nn}}\right)\frac{\partial^2 w}{\partial z_a \partial z_{\mathrm{nn},b}} = 0 \times \frac{\partial^2 w}{\partial z_a \partial z_{\mathrm{nn},b}}. \quad \text{(SI-48)}$$

Similar arguments show that all the other summands of $\beta_{ab,t}^{\text{cross isle}}$ are also zero.

### 3 Putting it all together

Summing eqs. (SI-4), (SI-24), (SI-33), (SI-47), we finally obtain the second order expansion of lineage fitness,

$$W(\mathbf{z},\phi_t) = 1 - \frac{1}{2}\sum_{a=1}^{n}\sum_{b=1}^{n}\sigma_{ab,t}h_{ab,t}(\bar{\mathbf{z}}_t) + \sum_{a=1}^{n}\xi_t(z_a)s_{a,t}(\bar{\mathbf{z}}_t) + \frac{1}{2}\sum_{a=1}^{n}\sum_{b=1}^{n}\xi_t(z_a)\xi_t(z_b)h_{ab,t}(\bar{\mathbf{z}}_t) + \mathcal{O}(\xi_t^3), \quad \text{(SI-49a)}$$

where

$$s_{a,t}(\bar{\mathbf{z}}_t) = \frac{\partial w}{\partial z_a} + (N-1)r_{2,t}^{\circ}(\bar{\mathbf{z}}_t)\frac{\partial w}{\partial z_{\mathrm{n},a}}, \quad \text{(SI-49b)}$$

and

$$h_{ab,t}(\bar{\mathbf{z}}_t) = h_{\mathrm{w},ab,t}(\bar{\mathbf{z}}_t) + h_{\mathrm{r},ab,t}(\bar{\mathbf{z}}_t), \quad \text{(SI-49c)}$$

with

$$h_{\mathrm{w},ab,t}(\bar{\mathbf{z}}_t) = \frac{\partial^2 w}{\partial z_a \partial z_b} + (N-1)\left[r_{2,t}^{\circ}(\bar{\mathbf{z}}_t)\left(\frac{\partial^2 w}{\partial z_{\mathrm{n},a}\partial z_{\mathrm{n},b}} + \frac{\partial^2 w}{\partial z_a \partial z_{\mathrm{n,b}}} + \frac{\partial^2 w}{\partial z_b \partial z_{\mathrm{n},a}}\right) + (N-2)r_{3,t}^{\circ}(\bar{\mathbf{z}}_t)\frac{\partial^2 w}{\partial z_{\mathrm{n},a}\partial z_{\mathrm{m},b}}\right],$$
$$h_{\mathrm{r},ab,t}(\bar{\mathbf{z}}_t) = \frac{\partial r_{2,t}(\mathbf{z})}{\partial z_a}(N-1)\frac{\partial w}{\partial z_{\mathrm{n},b}} + \frac{\partial r_{2,t}(\mathbf{z})}{\partial z_b}(N-1)\frac{\partial w}{\partial z_{\mathrm{n},a}}. \quad \text{(SI-49d)}$$

Eq. (SI-49) is equivalent to eq. B-1 – B-4 of the main text, in which we write the derivatives of individual fitness $w(\mathbf{z},\mathrm{n}_{\mu}(\mathbf{z}),\mathrm{e}(\phi_t))$ with respect to $z_a$, $z_{\mathrm{n},a}$, and $z_{\mathrm{nn},a}$, in terms of the derivatives of the individual fitness function $w(\mathbf{z}_i,\mathbf{z}_{-i},\bar{\mathbf{z}}_t)$ (eq. 5 of main text) with respect to $z_{i,a}$, $z_{j,a}$, and $z_{k,a}$, respectively; and add evaluation signs to all derivatives at the population mean $\bar{\mathbf{z}}_t$.
